# Whole-Brain Causal Discovery Using fMRI

**DOI:** 10.1101/2023.08.12.553067

**Authors:** Fahimeh Arab, AmirEmad Ghassami, Hamidreza Jamalabadi, Megan A. K. Peters, Erfan Nozari

## Abstract

Despite significant research, discovering causal relationships from fMRI remains a challenge. Popular methods such as Granger Causality and Dynamic Causal Modeling fall short in handling contemporaneous effects and latent common causes. Methods from causal structure learning literature can address these limitations but often scale poorly with network size and need acyclicity. In this study, we first provide a taxonomy of existing methods and compare their accuracy and efficiency on simulated fMRI from simple topologies. This analysis demonstrates a pressing need for more accurate and scalable methods, motivating the design of Causal discovery for Large-scale Low-resolution Time-series with Feedback (CaLLTiF). CaLLTiF is a constraint-based method that uses conditional independence between contemporaneous and lagged variables to extract causal relationships. On simulated fMRI from the macaque connectome, CaLLTiF achieves significantly higher accuracy and scalability than all tested alternatives. From resting-state human fMRI, CaLLTiF learns causal connectomes that are highly consistent across individuals, show clear top-down flow of causal effect from attention and default mode to sensorimotor networks, exhibit Euclidean distance-dependence in causal interactions, and are highly dominated by contemporaneous effects. Overall, this work takes a major step in enhancing causal discovery from whole-brain fMRI and defines a new standard for future investigations.

**AUTHOR SUMMARY:** Discovering causal relationships from fMRI data is challenging due to contemporaneous effects and latent causes. Popular methods like Granger Causality and Dynamic Causal Modeling struggle with these issues, especially in large networks. To address this, we introduce CaLLTiF, a scalable method that uses both lagged and contemporaneous variables to identify causal relationships. CaLLTiF outperforms various existing techniques in accuracy and scalability on simulated fMRI data. When applied to human resting-state fMRI, it reveals consistent and biologically-plausible patterns across individuals, with a clear top-down causal flow from attention and default mode networks to sensorimotor areas. Overall, this work advances the field of causal discovery in large-scale fMRI studies.

## INTRODUCTION

A major step in the global drive for understanding the brain (Adams et al., 2020; Amunts et al., 2016; Jorgenson et al., 2015; Okano, Miyawaki, & Kasai, 2015; Poo et al., 2016) is to move beyond correlations and understand the causal relationships among internal and external factors – a process often referred to as *causal discovery* (Assaad, Devijver, & Gaussier, 2022; Glymour, Zhang, & Spirtes, 2019). When possible, causal discovery can be greatly simplified by intervening in one variable and observing the effect in others. However, such interventions are often costly and/or infeasible, necessitating the significantly more challenging task of causal discovery from purely observational data.

A particularly rich set of observational data for the brain comes from functional MRI (fMRI) (Goense & Logothetis, 2008; Winder AT & PJ, 2017). The whole-brain coverage allowed by fMRI is valuable for causal discovery not only because it allows for purely data-driven and unbiased discovery of potentially unexpected causal relationships (Bressler & Menon, 2010; Bullmore & Sporns, 2012; Fornito & Bullmore, 2015), but also because of the great extent to which the presence of unobserved variables can complicate delineating causal adjacencies and orientations (Entner & Hoyer, 2010; Gerhardus & Runge, 2020; Hasan, Hossain, & Gani, 2023; Malinsky & Spirtes, 2018). Nevertheless, many characteristics of fMRI also make causal discovery challenging, including its large dimensionality, low temporal resolution, and indirect reflection of underlying neural processes (Ramsey et al., 2010).

This has motivated a large and growing body of literature on causal discovery from fMRI. A common approach for causal discovery using neuroimaging and neurophysiology data is Granger Causality (GC) (Seth, Barrett, & Barnett, 2015). GC has a long history in neuroscience (Barnett & Seth, 2014; Seth et al., 2015), but also has well-known limitations, including its lack of ability to account for contemporaneous causal relationships and the presence of latent nodes (see Supplementary Note 1 for a formal definition of ‘contemporaneous’ causal effects). The former is particularly important for fMRI. The temporal resolution in fMRI is typically within a few hundred milliseconds to several seconds (Huettel, Song, & McCarthy, 2009), which is about one order of magnitude slower than the time that it takes for neural signals to travel across the brain (Nunez & Srinivasan, 2006; Sutton & Begleiter, 1979; Sutton, Braren, Zubin, & John, 1965). Therefore, from one fMRI sample to the next, there is enough time for causal effects to flow between almost all pairs of nodes in the network (cf. a related in-depth discussion in (Nozari, Pasqualetti, & Cortés, 2019, Appendix A)). Such fast sub-TR interactions demonstrate themselves as causal effects that appear to be “contemporaneous” and can even be cyclic, making causal discovery significantly more challenging (cf. Supplementary Note 1). Similar to GC, Dynamic Causal Modeling (DCM) has also been widely used with fMRI data (K. Friston, Harrison, & Penny, 2003; K. Friston et al., 2019; K. J. Friston, Kahan, Biswal, & Razi, 2014; Stephan & Roebroeck, 2012) and fundamentally relies on the temporal order of a generative dynamical model to infer causation from correlations, making it similarly unable to account for contemporaneous causal relationships (K. Friston, Moran, & Seth, 2013; K. J. Friston, 2011; Logothetis, 2008).

Discovering causal relationships without reliance on time has been the subject of extensive research in the causal inference literature (Glymour et al., 2019; Pearl, 1988, 2009a, 2009b; Spirtes, Glymour, & Scheines, 2000; Spirtes & Zhang, 2016). A wide range of algorithmic solutions have been proposed (Chickering, 2002a; Glymour et al., 2019; Henry & Gates, 2017; Meek, 1995, 1997; Pearl, 2009a; Ramsey et al., 2010; Shimizu, Hoyer, Hyvärinen, & Kerminen, 2006; Smith et al., 2011; Spirtes & Glymour, 1991; Spirtes, Meek, & Richardson, 1995), which are often classified based on their methodology into constraint-based (Dawid, 1979; Pearl, 1988, 2009b), noise-based (Shimizu, 2014; Shimizu et al., 2006), and score-based (Chickering, 2002b; Heckerman, Geiger, & Chickering, 1995). However, it has remained largely unknown which of these algorithms are best suited for whole-brain fMRI causal discovery and how they perform relative to one another in terms of accuracy and scalability.

In this study, we first discuss and compare existing causal discovery algorithms for their suitability for whole-brain fMRI, demonstrate a large gap between what causal discovery for fMRI needs and what existing algorithms can achieve, propose CaLLTiF to address this gap, and demonstrate its higher accuracy and scalability on synthetic and real fMRI. Unless otherwise noted, all references to the words ‘graph’/‘network’ and ‘cycle’ mean a directed graph and directed cycle, respectively.

## RESULTS

### A Taxonomy of Causal Discovery for Whole-Brain fMRI

A vast array of algorithmic solutions exist for learning causal graphs from observational data, but not all are suitable for fMRI data. We selected a subset of state-of-the-art algorithms suitable for whole-brain fMRI data based on four criteria: (1) ability to learn cycles, (2) ability to learn contemporaneous effects, (3) assuming complete coverage of relevant variables in observed data, and (4) linearity (see Discussions). Table 1 shows several state-of-the-art methods that satisfy criteria (1)-(4). Multivariate Granger Causality (MVGC) (Barnett & Seth, 2014; Granger, 1969) does not satisfy criteria (2), but we still included it in our subsequent analyses due to its popularity in neuroscience (Ding, Chen, & Bressler, 2006; Goebel, Roebroeck, Kim, & Formisano, 2003; Liao et al., 2010; Roebroeck, Formisano, & Goebel, 2005). On the other hand, we excluded LiNG (Ramsey et al., 2018) from further analysis since it is considered by its proposers as generally inferior to the hybrid FASK algorithm (Sanchez-Romero et al., 2019). We also chose FASK for implementation over GANGO (Rawls, Kummerfeld, Mueller, Ma, & Zilverstand, 2022), a similarly hybrid method with the additional caveat of not having a unified publicly available code distribution.

**Table 1:** List of causal discovery methods suitable for use with whole-brain fMRI, divided by methodological category (constraint-, noise-, and score-based). All these methods (1) allow for cycles, (2) allow for contemporaneous effects, (3) assume complete coverage of relevant variables in observed data, and (4) learn linear relationships. The FASK algorithm is fundamentally hybrid and therefore listed as both constraint-based and noise-based.

| Category<br>Type | Constraint-based | Noise-based | Score-based |
| --- | --- | --- | --- |
| Time-series | PCMCI (Runge, Nowack, Kretschmer, Flaxman, & Sejdinovic, 2019),<br>PCMCI<sup>+</sup> (Runge, 2020) | VARLiNGAM (Hyvärinen, Zhang, Shimizu, & Hoyer, 2010) | DYNOTEARS (Pamfil et al., 2020) |
| Cross-sectional<br>with cycles | FASK (Sanchez-Romero et al., 2019) | FASK (Sanchez-Romero et al., 2019), LiNG (Lacerda, Spirtes, Ramsey, & Hoyer, 2008) | DGlearn (Ghassami, Yang, Kiyavash, & Zhang, 2020) |

We compared the accuracy of the resulting list of algorithms (MVGC, PCMCI, PCMCI^+^, VARLiNGAM, DYNOTEARS, FASK, and DGlearn) using simulated fMRI data from a benchmark of simple (5-10 nodes) networks introduced in (Sanchez-Romero et al., 2019). The ground truth graphs are shown in Figure 1a, and details on the fMRI time series generation for each node in these graphs are provided under Methods. To evaluate the success of each algorithm, we treated the causal discovery problem as a binary classification problem for each directed edge and calculated the resulting F1 score, both for the directed graphs as well as their underlying undirected graphs (see Methods for details). Figure 1b illustrates the distribution of F1 scores for all algorithms, combined across nine simple networks. The results show that the PCMCI algorithm achieved significantly higher median F1 score compared to all other algorithms over the directed graphs (all Cohen’s d *>* 0.23 and *p <* 10*^−^*^4^, pairwise one-sided Wilcoxon signed-rank test) and compared to all but DYNOTEARS over the underlying undirected graphs (Supplementary Figure 3, all Cohen’s d *>* 0.44 and *p <* 10*^−^*^29^, pairwise one-sided Wilcoxon signed-rank test. Also see Supplementary Figures 4 and 5 for precision and recall). The PCMCI algorithm also has the smallest computational complexity on simple networks, as seen from Figure 1c. Furthermore, our results indicate that FASK, DGlearn, and PCMCI^+^ (at their best values of hyperparameters) do not scale well with network size, forcing us to exclude them from further analysis as we move on to larger networks (see Supplementary Figures 13, 14, and 15).

**Figure 1:**
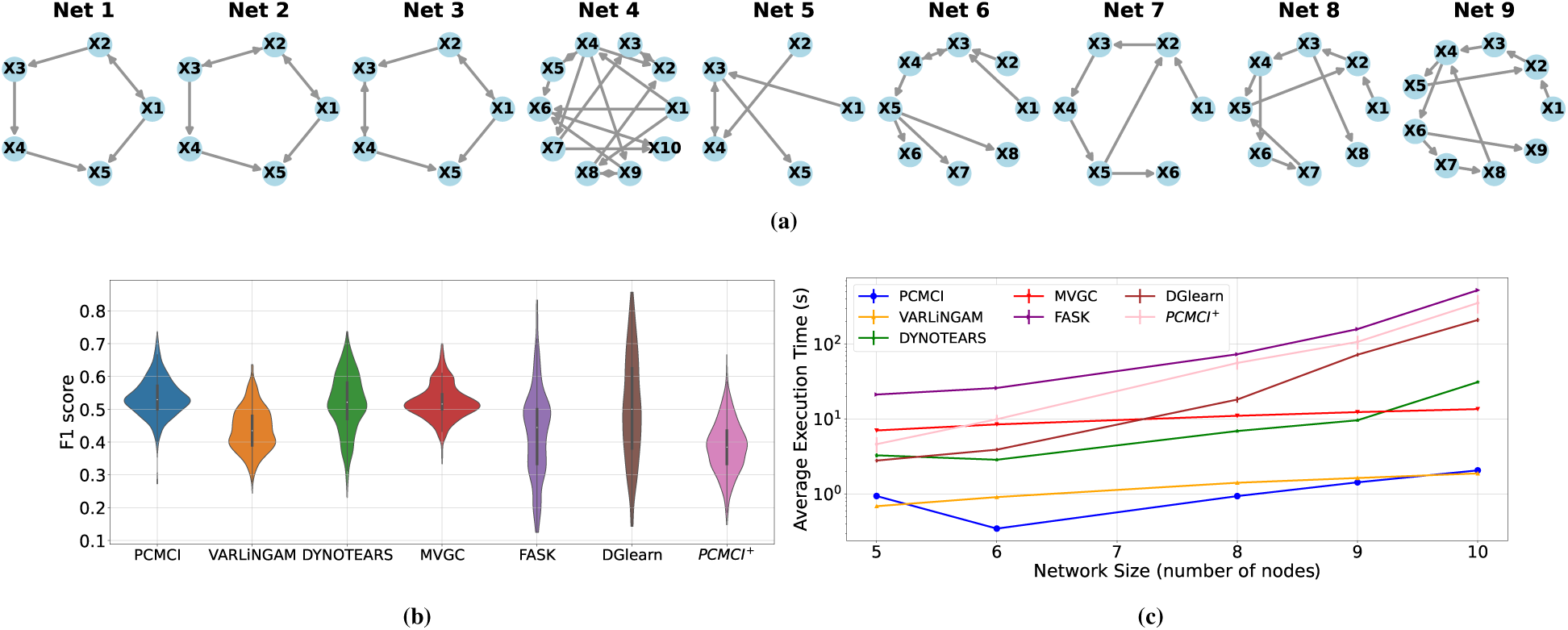
Results of comparing several state-of-the-art causal discovery algorithms over a benchmark of simulated fMRI (Sanchez-Romero et al., 2019) generated from simple networks with 5-10 nodes each. **(a)** Ground truth graphs of the simple networks in the benchmark. Despite all being small-scale, the graphs vary widely in their density, number of cycles, etc. **(b)** F1 scores of seven state-of-the-art algorithms (six from Table 1 and MVGC) for correctly identifying the full (directed) graphs. All methods are evaluated using optimized values of their respective hyperparameters (see Methods). The benchmark data includes 60 repetitions of fMRI data from each of the 9 graphs, so each violin plot is based on 540 F1 score samples. The PCMCI method achieves the highest median F1 score, both directed and undirected (see Supplementary Figure 3). **(c)** The mean execution time (averaged over all 60 repetitions) of different algorithms as a function of network size. Note the logarithmic scaling of the vertical axis. Error bars, though hardly visible, show 1 s.e.m.

Next, we compared the remaining four algorithms (PCMCI, MVGC, DYNOTEARS, and VARLiNGAM) on a larger-scale, more realistic simulated benchmark. The graph shown in Figure 2a, referred to as the Macaque_SmallDegree network, derived from the original Macaque_Full anatomical connectome and pruned to achieve average in-degree and out-degree of 1.8, consists of 28 nodes and 52 directed edges (Sanchez-Romero et al., 2019) but the generative model used to simulate fMRI data from this graph remains the same (see Methods for details). The distributions of F1 scores are shown in Figure 2b. PCMCI and MVGC achieved very similar success in learning both the directed graph and its underlying undirected graph, while significantly outperforming DYNOTEARS and VARLiNGAM. A similar result is obtained when comparing adjacency F1 scores for detecting the network’s underlying undirected graph (Supplementary Figure 17, also see Supplementary Figures 18 and 19 for precision and recall). As far as execution time is concerned, however, MVGC showed a significant advantage over PCMCI (Figure 2c). Therefore, despite its simplistic nature, MVGC was found most successful in causal discovery from Macaque_SmallDegree fMRI data (but also see Figure 3).

**Figure 2:**
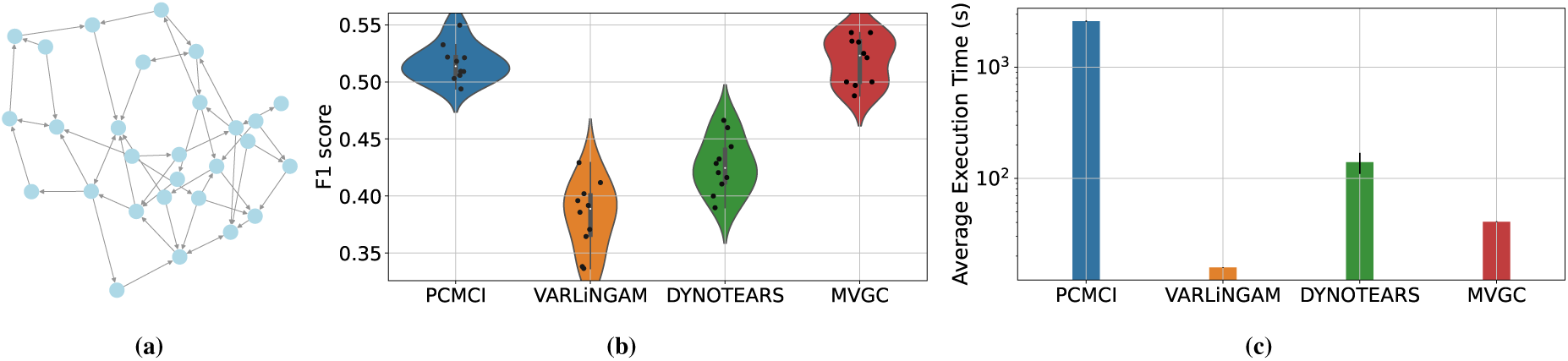
Comparing the scalable subset of algorithms from Figure 1 over simulated fMRI data from the Macaque SmallDegree bench-mark (Sanchez-Romero et al., 2019). **(a)** ground-truth Macaque SmallDegree network. **(b)** F1 scores of identifying the directed graph. Each distribution consists of 10 F1 scores calculated based on 10 repetitions of simulated data from the same underlying graph. **(c)** Mean execution times for each method (error bars show one standard deviation).

**Figure 3:**
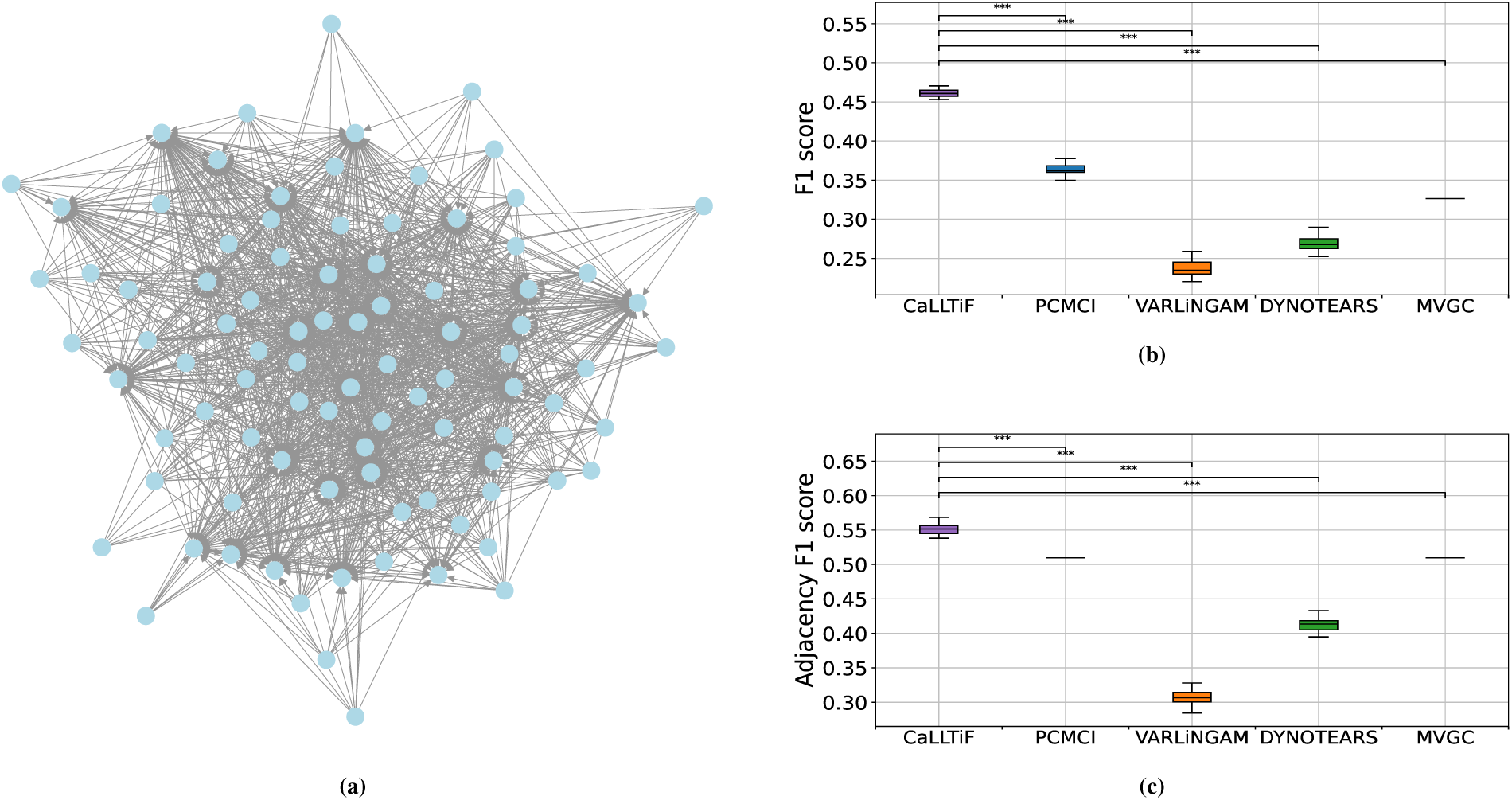
Comparisons between the proposed algorithm (CaLLTiF) and state-of-the-art alternatives over simulated fMRI from the Macaque Full connectome. **(a)** ground-truth Macaque Full network (See Supplementary Figure 25 for the heat map of the directed connectivity matrix). **(b,c)** Distributions of F1 scores for CaLLTiF and state-of-the-art alternatives in discovering the directed graph (b) and its underlying undirected graph (c). For all repetitions, the best performance of MVGC occurs at *α* = 0.5 which returns a complete graph, hence the point distributions for MVGC. *^∗∗∗^* denotes *p <* 0.001. All statistical comparisons are performed using a one-sided Wilcoxon signed-rank test. In all boxplots, the center line represents the median, the box spans the interquartile range (IQR), and the whiskers extend up to 1.5 times the IQR from the box limits.

### CaLLTiF: A New Causal Discovery Algorithm for Whole-Brain fMRI

The best-performing algorithms on Macaque_SmallDegree, i.e., PCMCI and MVGC, suffer from three main drawbacks: (1) poor scalability (only for PCMCI), (2) inability to learn directed contemporaneous effects (PCMCI only learns undirected contemporaneous effects while MVGC learns none), and (3) having sparsity-controlling hyperparameters that are subjectively selected in the absence of ground-truth graphs. In this section, we describe the design of a new algorithm based on PCMCI that mitigates these drawbacks and demonstrate its superior performance over existing methods.

Our first modification to PCMCI is with regard to scalability and computational complexity. The computational complexity of PCMCI depends heavily on the value of its ‘PC Alpha’ hyperparameter, which controls the sparsity of the set of potential common causes on which the algorithm conditions when checking the conditional independence of each pair of nodes (Supplementary Figures 16a and 37). Higher values of PC Alpha make these sets denser and accordingly decrease statistical power in the subsequent conditional independence tests, *ultimately conditioning on all other nodes (and all of their lags) when PC Alpha = 1*. Nevertheless, interestingly, our experiments on the Macaque_SmallDegree data show that the maximum achievable accuracy of PCMCI (i.e., F1 score maximized over Alpha Level for each fixed value of PC Alpha) monotonically increases with PC Alpha, reaching its maximum at PC Alpha = 1 (Supplementary Figures 16b and 16c). Therefore, while this may seem to cause a trade-off between accuracy and scalability, it is in fact an opportunity for maximizing both. At PC Alpha = 1, the PC part of PCMCI (a.k.a. the S1 algorithm in (Runge et al., 2019)) is theoretically guaranteed to return a complete conditioning set for all pairs of nodes, and can thus be skipped entirely. The PC part is further responsible for the poor scalability of PCMCI. Thus its removal significantly improves the computational efficiency of the resulting algorithm *without compromising accuracy* (cf. Discussions for a potential explanation of why conditioning on all other nodes may improve accuracy despite lowering statistical power).

Our second modification addresses the lack of directed contemporaneous causal effects (see Introduction for why these effects are particularly important in fMRI). By default, MVGC returns no contemporaneous edges and PCMCI returns *◦−◦* ones which only indicate the presence of significant partial correlations but does not resolve between *→*, *→*, or ⇆. However, we know from decades of tract tracing studies that reciprocal connections are significantly more common than unidirectional connections in the primate brain (Felleman & Van Essen, 1991; Markov et al., 2014; Tigges, Spatz, & Tigges, 1973). Therefore, we replace all *◦−◦* edges returned by PCMCI by the more likely choice of ⇆. The only exception comes from (the often minority of) pairs of nodes that have a *lagged* directed edge between them (i.e., an edge of the form *X_i_*(*t − τ*) *→ X_j_*(*t*) for *τ >* 0, see Methods), in which case we leave the direction of the contemporaneous effect between them the same as their lagged effect.

Figure 3 shows how the resulting CaLLTiF algorithm performs on a synthetic fMRI dataset generated from the significantly larger macaque structural connectome called Macaque_Full with 91 nodes and 1,615 ground-truth edges (Figure 3a, see also Methods). CaLLTiF has a significantly higher F1 score compared to PCMCI, VARLiNGAM, DYNOTEARS, and MVGC in its discovery of the directed graph (Figure 3b, all Cohen’s *d >* 15, *p <* 10*^−^*^6^, one-sided Wilcoxon signed-rank test) as well as its underlying undirected graph (Figure 3c, all Cohen’s *d >* 7, *p <* 10*^−^*^6^, one-sided Wilcoxon signed-rank test). Precisions and recalls are also shown in Supplementary Figures 26 and 27. We also compared CaLLTiF (and PCMCI) against a middle-ground ‘Mixed-PCMCI’ variant where the *◦−◦* edges returned by PCMCI are used only in the computation of adjacency F1 score (Supplementary Figures 28-30, see also Methods). Mixed-PCMCI benefits from contemporaneous effects as much as CaLLTiF in terms of adjacency F1 score, but not so in terms of full F1 score, further motivating the inclusion of directed contemporaneous connections as done in CaLLTiF. Detailed performances of all compared algorithms are provided in Supplementary Figures 29-36.

Finally, the third aspect in which CaLLTiF departs from PCMCI is the choice of sparsity-controlling hyperparameter ‘Alpha Level’. Most, if not all, algorithms for causal discovery have at least one hyperparameter (often a threshold) that controls the sparsity of the resulting graphs. Different from PC Alpha described earlier, Alpha Level in PCMCI is the standard type-I error bound in determining statistical significance in *each* partial correlation test (cf. Supplementary Figure 16). By default, Alpha Level is selected subjectively, based on domain knowledge and expected level of sparsity. However, in CaLLTiF, we select Alpha Level objectively based on a novel method for correction for multiple comparisons (see Methods) that occur when collapsing a time-series graph over lagged variables into a final summary graph. This step is critical, particularly in the absence of ground-truth connectivity, to ensure that we have statistical confidence in every edge of the final summary graph returned by CaLLTiF.

In summary, CaLLTiF starts by constructing an extended time-lagged graph among all the variables *X_i_*(*t − τ*)*, i* = 1*, . . ., n* and all lags *τ* = 0, 1*, . . ., τ*_max_. To establish a causal link between any pair of variables *X_i_*(*t − τ*) and *X_j_*(*t*), CaLLTiF performs a conditional independence test (using linear partial correlation) between *X_i_*(*t − τ*) and *X_j_*(*t*), conditioned on all other *lagged* variables (*X_k_*(*t − s*)*, s* = 1*, . . ., τ*_max_). A causal link is established if the null hypothesis of conditional independence is rejected at a significance threshold ‘Alpha Level’. By default, ‘Alpha Level’ is selected based on CaLLTiF’s type I error control, but it can also be optimized in simulated data using ground-truth knowledge. If *τ >* 0, the direction of the edge is clearly *X_i_*(*t − τ*) *→ X_j_*(*t*). When *τ* = 0, CaLLTiF returns *X_i_*(*t*)*⇆X_j_*(*t*) if no other edges exist between *X_i_* and *X_j_* at higher lags, and places the edge(s) consistent with the corresponding lagged direction(s) otherwise. Finally, the extended time-lagged graph is collapsed into a summary graph by taking an OR operation for each edge across all lags. For further details about CaLLTiF, see Methods. A pseudocode for CaLLTiF is given in Algorithm 1 and a formal analysis of its computational complexity can be found in Supplementary Note 4.

### Causal Discovery from Resting-State Human fMRI

We next applied CaLLTiF on resting-state fMRI from 200 subjects from the Human Connectome Project (HCP) (See Methods). Each scan from each subject was parcellated into 100 cortical and 16 subcortical regions. CaLLTiF was then performed on all four resting-state scans for each subject, resulting in one causal graph per individual.

#### Learned causal graphs are highly consistent across subjects

Despite individual differences, a remarkably common causal connectome emerged across subjects. Figure 4a shows the average causal graph among the subjects and Figure 4b shows the intersection graph that contained the edges *common across all subjects*. Due to the binary nature of individual graphs, the former can also be viewed as a matrix of probabilities, where entry (*i, j*) shows the probability of region *i* causing region *j* across all subjects. As a result of the significant commonalities that exist in the causal graphs among subjects, the average graph has a bimodal distribution, with the vast majority of average weights being close to either 0 or 1. These extreme values of average weights can also be seen as a measure of the confidence of the algorithm in the presence or lack of most edges, and have a clear contrast (*p* = 0, Kolmogorov-Smirnov test) with the weights of the average of randomized surrogate graphs generated *independently* across subjects (Supplementary Figure 44). In the absence of a ground truth causal connectivity for direct comparison, such strong commonalities among subjects serve as an alternative measure of validation and provide insights into the general patterns and characteristics of the causal relationships in a resting brain.

**Figure 4:**
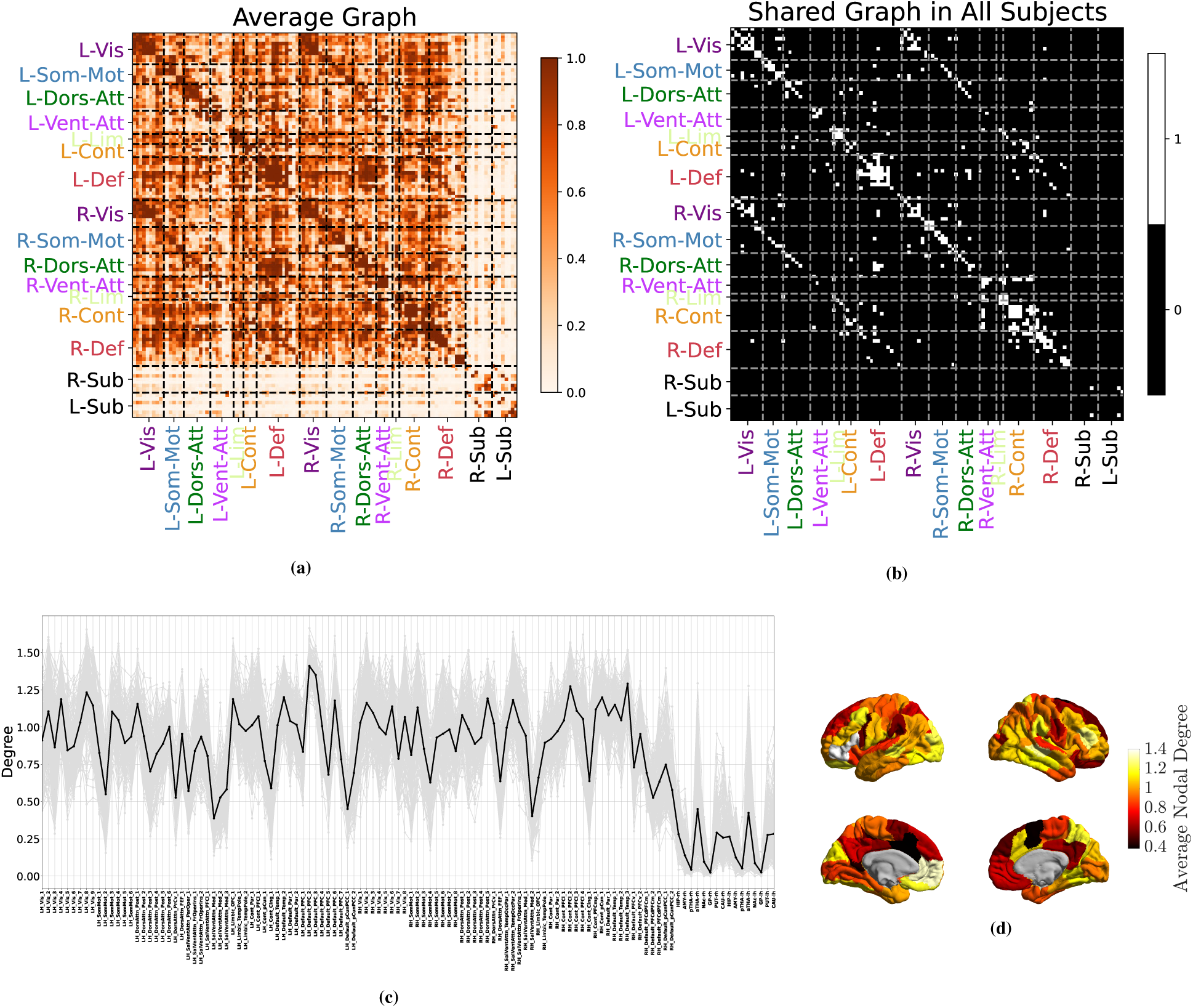
Learned causal graphs are highly consistent across subjects. **(a)** Average causal graph, computed as the mean of all the causal graphs of 200 subjects. The color of the (*i, j*) entry in this matrix shows how likely it is for node *i* to cause node *j*. A clear distinction can be seen in the causal links among cortical regions (colored labels) and subcortical ones (black labels). **(b)** The shared causal graph, containing only the edges that are present in all 200 subjects. This graph is dominated by diagonal elements (self- and within-subnetwork causation) and links among symmetrically located regions across the two hemispheres. **(c)** Distribution of nodal degree, computed separately for each node in the causal graph and each subject. Gray colors correspond to different subjects and the black line shows the average nodal degrees across subjects. **(d)** Average cortical nodal degree (black line in (c)) shown over the brain cortical surface.

Nodal centralities also show strong consistency among the subjects. Figure 4c shows the nodal degrees for all subjects (gray lines) as well as its mean across subjects (black line, also depicted in Figure 4d). Statistically significant differences exist between the degree distributions of many pairs of nodes (about 90% of the pairs have *p <* 0.001, two-sided Wilcoxon signed-rank test, computed between nodal degrees of each pair of parcels), while significant correlations exist between nodal degrees for all pairs of subjects (all pairs have 0.56 *≤ r ≤* 0.96, *p <* 10*^−^*^10^, Pearson Correlation Coefficient, computed based on the nodal degrees of each pair of subjects separately). Note that the correlations of degrees across subjects are also remarkably higher than what would be implied by the correlations of the graphs themselves (Supplementary Figure 43e). Similar consistency for in-degree, out-degree, betweenness, and eigenvector centralities can be observed among subjects (Supplementary Figures 45-48). Consistently across subjects, medial ventral attention regions, cingulate cortices, and lateral primary sensorimotor areas show particularly low nodal degrees across both hemispheres, whereas bilateral default mode areas, particularly the left ventromedial prefrontal cortex, show notably strong nodal degrees. Bilateral anterior thalami are particularly causally connected compared to other subcortical regions, even though subcortical areas have significantly lower degrees than cortical areas in general, with bilateral posterior thalami, nuclei accumbens, and globus pallidi showing the least causal connections across the whole brain at rest.

Causal graphs are also sparser and more consistent across subjects compared to functional connectivity. A major motivation for building causal connectomes is the removal of spurious connections in functional connectivity (FC) profiles that reflect mere correlation but no causation. For causal graphs learned by CaLLTiF, we indeed observed significantly lower edge density compared to FC graphs (see Methods for details on the computation of FC graphs) (Supplementary Figures 43a,43b, no overlap existing between the support of the two distributions). In fact, FC graphs included approximately 95% of CaLLTiF’s discovered causal edges (Supplementary Figure 43c), while only about half of all functional connectivity edges are also causal (Supplementary Figure 43d). Interestingly, among the approximately 5% of causal edges that were not in the FC graphs, the majority came from non-zero lags. This is remarkable, given that causal edges from non-zero lags are significantly fewer in general (cf. Figure 6a), but are fundamentally not discoverable by FC which only measures contemporaneous co-fluctuations. Moreover, causal connectomes are significantly more consistent across subjects compared to FC connectomes (Supplementary Figure 43e, Cohen’d *>* 2, *p <* 0.001, one-sided Wilcoxon signed-rank test), further reinforcing the expectation that causal edges are “pruned” and more reliable compared to functional edges.

#### Net resting-state causal effect flows from attention and default mode to sensorimotor networks

One of the main advantages of directed causal connectomes over undirected functional and structural connectomes is the former’s ability to show the directed flow of causal effect between brain regions. In graphs learned by CaLLTiF, nodal causal flows (outflow minus inflow, see Methods) are also highly consistent across subjects (Figure 5c,5d), even though the two notions of centrality are generally dissociated across parcels (Figure 5b and Supplementary Figure 53). On average across all subjects, we observed particularly high causal flows (source-ness) in several regions of bilateral medial ventral attention networks, specific dorsal attention areas (ventral precentral, ventral frontal cortices, and frontal eye fields), and bilateral hippocampi, even though subcortical areas are much less connected to the rest of the network in general. In contrast, bilateral visual areas show the strongest negative causal flow (sink-ness) across all subjects. There is also notable variability among parcels within a subnetwork, such as the notable bilateral contrast between the strongly positive and weakly negative causal flows of frontal and posterior parts of the dorsal attention network, respectively.

**Figure 5:**
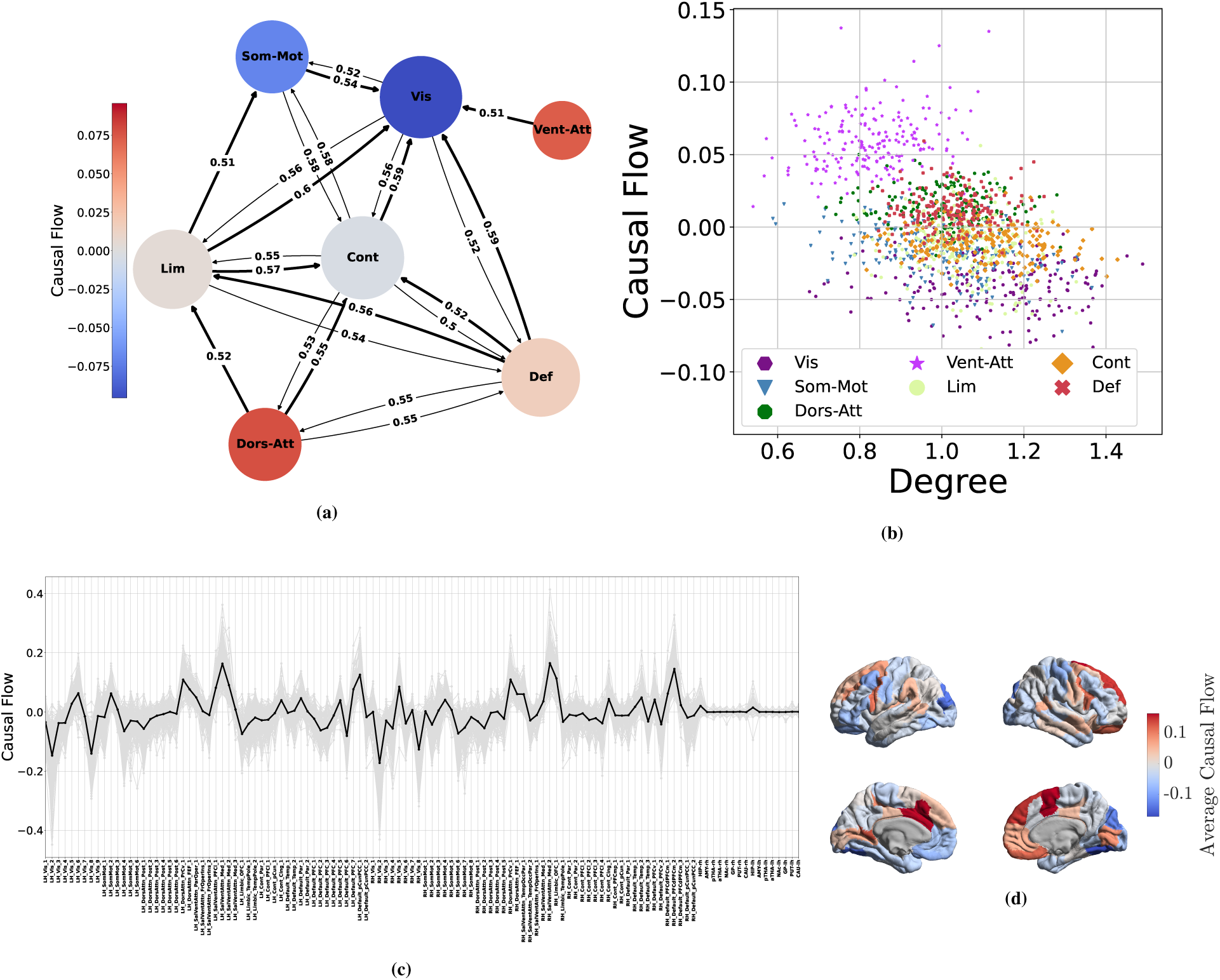
Patterns of causal flow across subjects. **(a)** The average subnetwork graph, computed as the mean of subnetwork graphs of all the subjects. In the subnetwork graph of each subject, the weight of an edge from subnetwork *i* to *j* is the number of nodes in subnetwork *i* that connect to nodes in subnetwork *j*, normalized by the number of all possible edges between these subnetworks. Edges with weights less than 0.5 are removed for better visualizations. To further ease the visual inspection of causal flows, if two networks are bidirectionally coupled we have shown the stronger edge with a thicker line (see Supplementary Figures 51, 52 for the weight matrix before thresholding and the corresponding nodal degrees and causal flows.) **(b)** The joint distributions of causal flow and degree for each “node” of the subnetwork graphs across all subjects. **(c)** Distribution of nodal causal flow, computed separately for each node in the causal graph and each subject. Gray colors correspond to different subjects and the black line shows the average nodal degrees across subjects. **(d)** Average cortical nodal causal flows (black line in (c)) shown over the brain cortical surface.

To better assess the overall net causal effects between different functional networks, we computed an average “subnetwork graph” in which each node represents a functional cortical network and edges denote thresholded average directed connectivity from one functional network to another (see Methods for detailed computations). The result is illustrated in Figure 5a. Ventral attention and visual networks are clearly the strongest source and sink of causal flow, respectively. The dorsal attention and somatomotor networks are also a clear (though weaker) source and sink, respectively. The default mode network (DMN) is also a net source of causal flow, even though its outflows and inflows are nearly balanced. Similarly, the control and limbic networks have near-zero causal flows (near-balanced inflow and outflow). Several directed paths, however, can be seen from both attention and default mode networks to sensorimotor networks through the limbic and control networks. Therefore, in summary, causal graphs learned by CaLLTiF show the strongest net resting state causal effect to flow from the ventral and dorsal attention as well as the default mode networks, through control and limbic networks, towards sensorimotor networks. The DMN, control, and limbic networks have large average degrees (Supplementary Figure 52) and near-balanced causal flows, making them hubs that largely distribute the flow of causal effect in the resting-state causal connectome (see the Discussion section for a more detailed analysis of this network).

#### Casual graphs are strongly dominated by contemporaneous and lag-1 connections

Given that the final causal graph returned by CaLLTiF is a union over subgraphs at different lags (cf. Methods), we can go back and ask how much causal effects in each lag have contributed to the final graph. Figure 6a shows the percentage of edges in the final graph that exist *only* in one lag (including lag 0, or contemporaneous edges). Increasing the lag order resulted in significantly sparser single-lag subgraphs, which contributed less to the end result. In particular, approximately 70% of the end graphs came from lag 0 alone, a pattern that appears consistently across all subjects (Supplementary Figure 57). Even further, such contemporaneous edges are substantially stronger than edges from lags 1-3 (Figure 6b). This further confirms that the contemporaneous effects are particularly important for fMRI, where most neural dynamics occur at timescales shorter than 1 TR (typically shorter than 1-2 seconds). This is even the case in HCP data, with TR = 0.72s which is among the shortest TRs currently available in fMRI research. That being said, all lags had a non-zero (and significant by construction) contribution to the end graph in all subjects. Even lag 3 had a median of approximately 0.2% unique contributions to the final graph across subjects. We also found very small intersections among lags. This not only highlights the importance of considering multiple lags rather than just the first one or two but also demonstrates that it is incorrect to assume that if one region causes another, that causation will appear continuously across all lags. In summary, we found contemporaneous effects dominant in the final causal graphs of CaLLTiF, even though all lags had significantly non-zero and mostly unique contributions.

**Figure 6:**
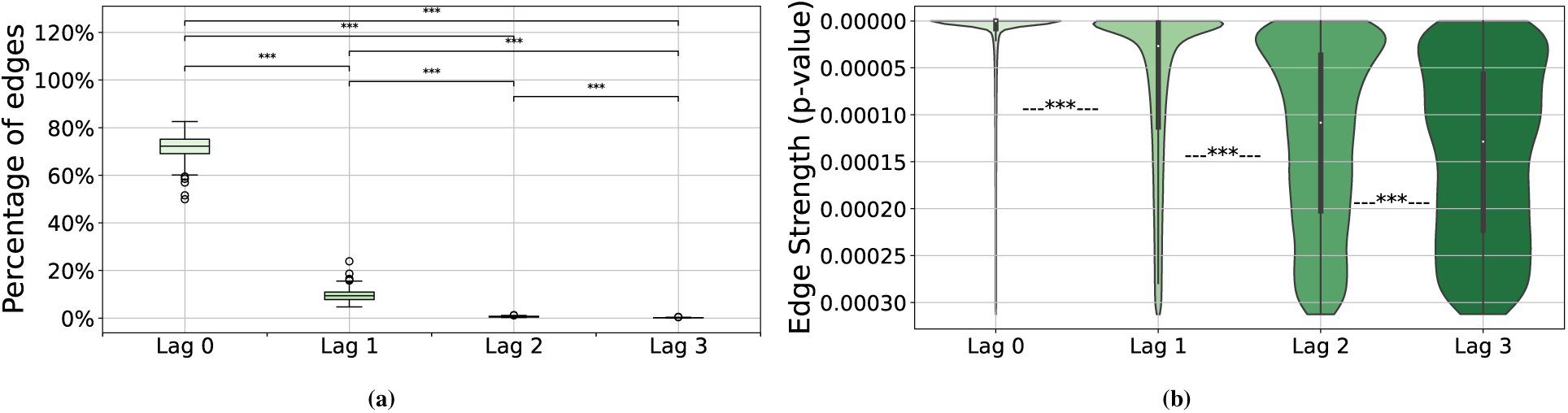
The contributions of each lag to the final causal graph in CaLLTiF. **(a)** For each lag, the box plot shows the distribution of the percentage of edges that come only from that lag across subjects. In all boxplots, the center line represents the median, the box spans the interquartile range (IQR), the whiskers extend up to 1.5 times the IQR from the box limits, and individual points beyond the whiskers indicate outliers.**(b)** The strength (statistical significance) of edges across lags. For each edge within the subgraph of each lag, we define its strength as the p-value of the partial correlation test that was used to conclude the presence of that edge (see Methods) even though all edges have a statistically significant p-value by definition, edges in larger lags are significantly closer to the threshold for significance than those in smaller lags. *^∗∗∗^* = *p <* 0.001, one-sided Wilcoxon rank-sum test.

#### Causal connections are modulated by pairwise Euclidean distance

As one would expect from a network learned over a set of nodes embedded in physical space, the causal graphs learned by CaLLTiF are modulated in a number of ways by the Euclidean distance between pairs of nodes. First, we found degree similarity (correlation coefficient between nodal degrees of two parcels over all subjects) to decay statistically significantly, though weakly in effect size, with parcel distance (Pearson *r* = *−*0.12, *p* = 10*^−^*^43^, 95% confidence interval (*−*0.14*, −*0.1)) as shown in Figure 7a (See Supplementary Figures 54 and 55 for separate maps of degree similarities and pairwise nodal distances). This relationship is stronger among intra-hemispheric parcels (Pearson *r* = *−*0.27*, p* = 10*^−^*^82^, 95% confidence interval (*−*0.29*, −*0.24)) where connections are denser and shorter-distance, compared to inter-hemispheric parcels (Pearson *r* = *−*0.09*, p* = 10*^−^*^5^, 95% confidence interval (*−*0.13*, −*0.05)). Thus, in summary, nodes that are physically closer to each other also have more similar causal connections to the rest of the network, particularly if they belong to the same hemisphere.

**Figure 7:**
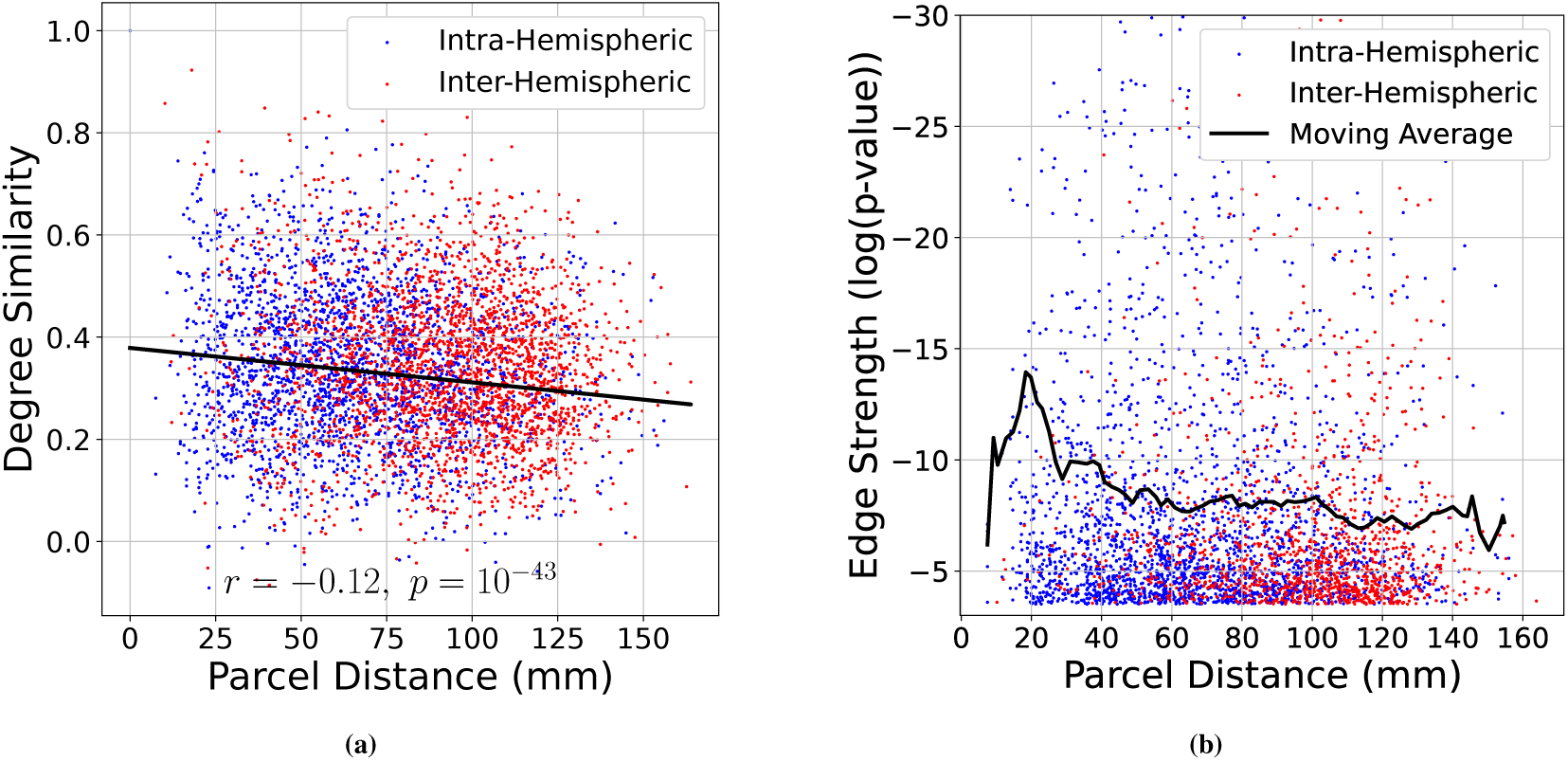
Effect of Euclidean distance on edge attributes. **(a)**Degree similarity (correlation coefficient between nodal degrees of two parcels over all subjects) as a function of the Euclidean distance between the parcels. Parcel pairs in the same hemisphere (intra-hemispheric) and parcels in two different hemispheres (inter-hemispheric) are shown in blue and red, respectively. Degree similarity decays statistically significantly with parcel distance (Pearson *r* = *−*0.12, *p* = 10*^−^*^43^, 95% confidence interval (*−*0.14*, −*0.1)), much more so among intra-hemispheric parcels (Pearson *r* = *−*0.27, *p* = 10*^−^*^82^, 95% confidence interval (*−*0.29*, −*0.24)) than inter-hemispheric ones (Pearson *r* = *−*0.09, *p* = 10*^−^*^5^, 95% confidence interval (*−*0.13*, −*0.05)). **(b)** Edge strength (as in Figure 6b) as a function of Euclidean distance between edge endpoints (note the inverted vertical axis). The solid line shows the corresponding moving average of log(*p*) with 10mm window size and 8mm window overlap. The upper limit of the vertical axis is limited to *−*30 for better visualization.

The strength of CaLLTiF edges is also modulated by the Euclidean distance between edge endpoints, even though we observed that there are approximately as many long-distance edges as short ones (See Supplementary Figures 56). We define the strength of each edge in the final graph (union over lags) as the *minimum* p-value of respective partial correlation tests across all lags (cf. Methods). As seen from Figure 7b, the mean strength of causal edges (black solid line) initially increases with the Euclidean length of the edge until about 20mm and then decays with Euclidean edge length thereafter.

Finally, we found no major differences between the Euclidean distances of edges contributed by different lags. Given that causal effects take time to spread along axonal fibers throughout the brain, one might expect physically-closer pairs of nodes to be connected by lower-lag edges and more distant pairs of nodes to be connected by larger-lag edges. However, as seen in Supplementary Figure 58, this is not quite the case. Given the slow sampling of fMRI, even the most distant regions can causally affect each other in time scales shorter than 1 TR. Thus, the observation that the physical distance of pairs of nodes was not related to edge lag should not be taken as an indication that such relationships would – or would not – be observed when sampling with higher temporal precision.

#### Degree, but not casual flow, shows significant laterality and gender differences

We observed that nodal degrees were statistically significantly higher in the right hemisphere (Figure 8a, Cohen’s d = 0.07 and *p* = 10*^−^*^48^, one-sided Wilcoxon signed-rank test), even though no such laterality was found in nodal causal flows (Figure 8b, Cohen’s d = 0.02 and *p* = 0.23, one-sided Wilcoxon signed-rank test). To understand which subnetworks might be playing a stronger role in the hemispheric asymmetry observed in the distribution of nodal degrees, Figure 8c shows the mean degrees of corresponding pairs of regions in the left and right hemispheres, color-coded by functional subnetworks (cf. Supplementary Figures 49 for separate plots per subnetwork). The ventral attention, dorsal attention, and executive control networks show clearly larger causal degrees in the right hemisphere, whereas the limbic network and DMN have larger causal degrees in the left hemisphere. A similar plot for causal flows (Figure 8d, Supplementary Figure 50) shows a lot more symmetry, except for the limbic network which shows exceptionally higher causal flows (i.e., source-ness) in the right compared to the left hemispheres. The DMN also shows some asymmetry in its causal flow, where right DMN nodes are mostly sources of causal flow whereas left DMN causal flows are more evenly distributed around zero. Thus, in summary, various functional subnetworks show laterality in degree distributions, culminating in a net increase in right vs. left nodal degrees. Causal flows, however, are mostly symmetric, except for the limbic network which shows a strong flow from the right to the left hemisphere.

**Figure 8:**
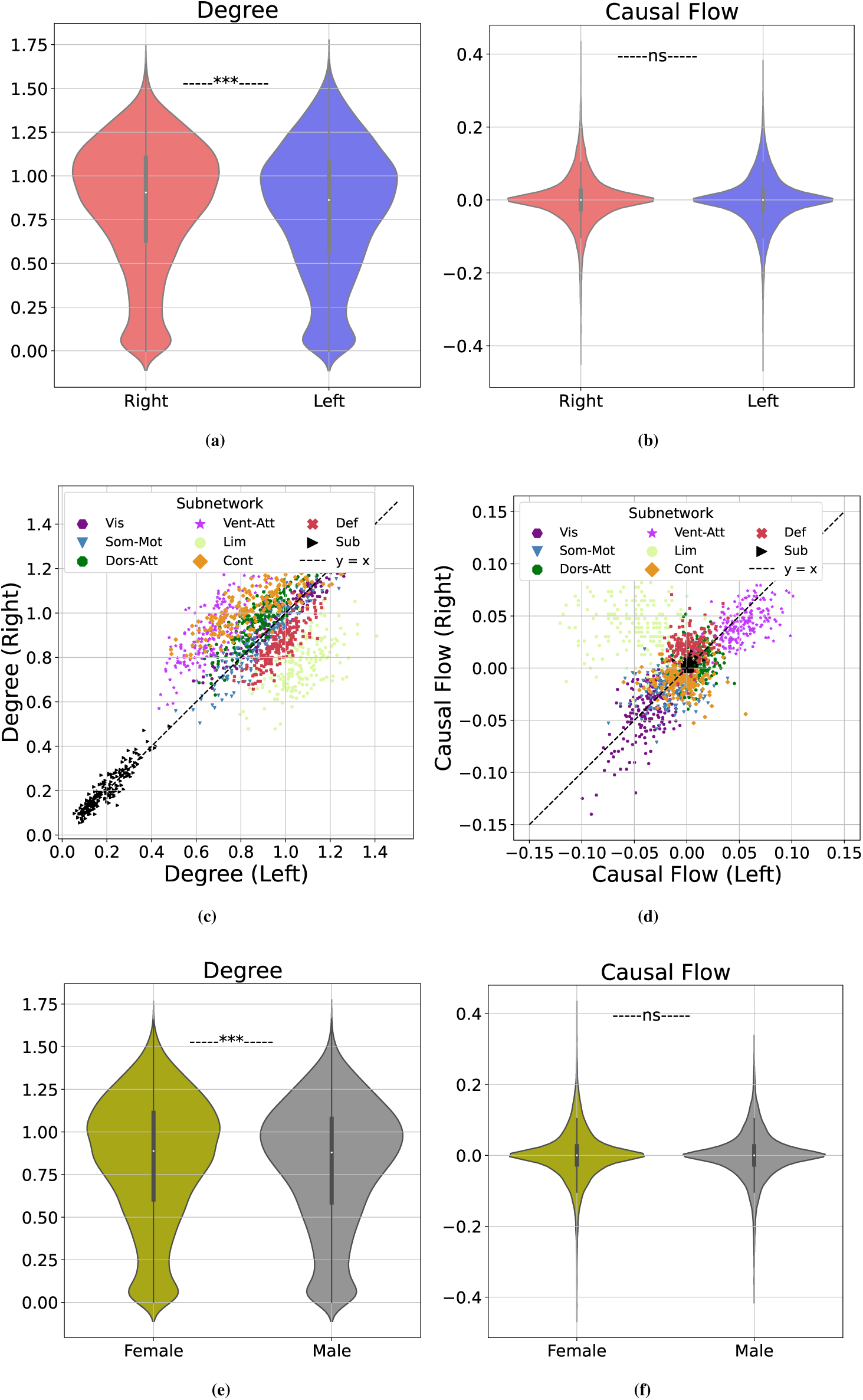
Hemispheric laterality and gender differences in causal connectomes. **(a)** Distributions of nodal degree in the right and left hemispheres, combined across all subjects. **(b)** Similar to **(a)** but for causal flows. **(c)** Nodal degrees, averaged across subjects and color-coded by functional subnetwork, for pairs of corresponding parcels within the right and left hemispheres. To properly pair nodes across the two hemispheres, degrees of all the parcels with the same label in the Schaefer/Tian atlas were averaged and then paired. Deviations from the dashed y = x line indicate laterality. **(d)** Similar to **(c)** but for causal flows. **(e)** Distributions of nodal degrees in female and male subjects, combined across all brain regions. **(f)** Similar to **(e)**, but for causal flows.

Similarly, degree, but not casual flow, shows a small but statistically significant difference between men and women. In causal connectomes learned by CaLLTiF, we found nodal degrees to be statistically significantly higher in women compared to men (Figure 8e, Cohen’s d *>* 0.05, *p <* 10*^−^*^5^, one-sided Wilcoxon rank-sum test). Nodal causal flows, on the other hand, were statistically indistinguishable between men and women (Figure 8f, Cohen’s d = *−*3.77 *×* 10*^−^*^18^, *p* = 0.81, one-sided Wilcoxon rank-sum test). These result demonstrate that nodal degrees in causal connectomes are generally more heterogeneous and sensitive while causal flows tend to be more homogeneous and stereotyped across individuals and hemispheres. Further research is needed to pinpoint the root causes of these differences (and lacks thereof) as well as potential implications of them in health and disease.

## DISCUSSION

In this study, we investigated the problem of whole-brain causal discovery from fMRI. We first comprehensively compared existing causal discovery techniques suitable for whole-brain fMRI by examining both theoretical properties and numerical outcomes on simulated fMRI. To address the limitations of existing algorithms, we proposed CaLLTiF which improves upon the state of the art in several directions including learning contemporaneous edges and cycles, type I error control, and scalability. A core aspect of CaLLTiF is its treatment of contemporaneous effects. Our results with the HCP data (Figures 6a and 6b) confirmed the importance of being able to reveal such “contemporaneous” effects, where these effects accounted for the majority and strongest of network edges. Further, the distributions of edges with different Euclidean distances at each lag (Supplementary Figure 58) demonstrates how broadly neural signals can propagate across the brain in one TR interval, even with the relatively fast sampling (TR = 0.72s) in the HCP dataset.

Furthermore, in interpreting CaLLTiF’s outputs, it is important to note its by-design conservative method of correction for temporal multiple comparisons. In the Macaque_Full simulated dataset where the ground truth is known, we found Alpha Level = 0.01 to maximize the F1 score, while CaLLTiF’s correction for temporal multiple comparisons would have suggested 0.01*/*32 = 0.0003 (cf. Equation (4)) and thus would have obtained sparser graphs. Similarly, we obtained causal graphs from human fMRI that are about 30-55% dense across all subjects and 40-60% dense among cortical nodes (Supplementary Figure 43a). These are generally consistent with, but sparser than, the near-66% cortical density found using tract-tracing results in non-human primates (Markov et al., 2014). In other words, graphs returned by CaLLTiF are likely to have higher precision but lower recall than what would maximize the F1 score. This conservativeness is by design and desirable *in the absence of a ground-truth causal graph*, ensuring that we have strong confidence in all discovered edges (at least 99% confidence in every detected edge in our experiments with human fMRI). Nevertheless, CaLLTiF’s level of conservativeness can also be tuned as needed by tuning its pre-correction significance threshold (*q* in Equation (4)).

An unexpected finding of our study was the higher accuracy of causal discovery when conditioning pairwise independence tests (see Equation (1)) on all other nodes in the network, as done in CaLLTiF, compared to using a more restricted parent set found by PCMCI (cf. Supplementary Figure 16). The approach taken by PCMCI increases statistical power (cf. the trend of optimal ‘Alpha Level’ values in Supplementary Figure 16b), but can significantly increase type I error in the presence of contemporaneous effects. Even further, we found that even using the (lagged) ground-truth parent sets for each node leads to a lower F1 score compared to using complete conditioning sets (Supplementary Figure 39-41). This is likely because CaLLTiF’s conditioning on the *past* of all variables serves as a proxy for the missing contemporaneous parents that should have been conditioned on. On the other hand, one may wonder if this issue could have been better resolved by conditioning on contemporaneous variables themselves. However, this can result in spurious statistical dependence if conditioning on all contemporaneous variables (consider, e.g., testing *X_i_* ⊥̸⊥ *X_j_*|*X_k_* with the ground-truth causal graph *X_i_→ X_k_ → X_j_*). For a detailed discussion on this see Supplementary Note 3.

### Causal connectivity during resting state

When applying CaLLTiF to resting state human fMRI, we found the strongest causal effect to flow from attention to sensorimotor networks. The strongest sources and sinks were the ventral attention and visual networks, followed by dorsal attention and somatomotor networks, respectively. Despite the lack of “ground-truth” connectivity as in simulated data, we can still evaluate these findings based on their agreement with prior findings on the roles of resting state networks. The dorsal attention network (involving regions in the intraparietal and superior frontal cortex) is commonly believed to handle top-down selection processes and is only modulated by stimulus detection, while the ventral attention network (including areas such as the temporoparietal and inferior frontal cortex) specializes in detecting behaviorally relevant stimuli, particularly those that are salient or unexpected, and directing attention to them (Corbetta & Shulman, 2002). These networks exhibit internally correlated activity patterns (particularly during the task) (Corbetta, Patel, & Shulman, 2008) and their flexible interaction facilitates dynamic attentional control aligned with both top-down goals and bottom-up sensory inputs (Vossel, Geng, & Fink, 2014). Nonetheless, the ventral attention network is more strongly involved in the “reorientation” of attention, namely, interrupting one thought process and orienting attention towards something salient, while the dorsal attention network is more strongly implicated in focused and guided attentional tasks such as visual search under high short-term memory load (Vossel et al., 2014).

Arguably, resting state activity is more aligned with the former (salience-based reorientation) than the latter (goal-driven focused attention). Despite a lack of sensory salience, attention is frequently reoriented during periods of rest by endogenously-salient thoughts and memories. Intermittent periods of focused attention can also arise, particularly given the long durations of each resting-state scan (*∼* 15min). Our findings thus confirm and corroborate the existing hypotheses about the roles of attention networks and how they jointly but unevenly drive brain dynamics during rest. Furthermore, due to the lack of meaningful sensory (particularly visual) input during a resting state scan, sensory areas receive more top-down influence from higher-order cortices than they provide bottom-up information to them. As such, the sink-ness of sensorimotor areas in graphs learned by CaLLTiF is arguably more consistent with the nature of resting state activity compared to a contrasting, sensory-driven flow found, e.g., in (Rawls et al., 2022). Finally, we found the DMN to be both a hub and a weak source of causal flow, which is consistent with its well-known role in resting state dynamics (Andrews-Hanna, Smallwood, & Spreng, 2014; Greicius, Krasnow, Reiss, & Menon, 2003; Raichle et al., 2001).

Resting-state causal graphs learned by CaLLTiF are also notably aligned with the literature in terms of the laterality of different functional subnetworks (Figure 8c and Supplementary Figure 49). Several studies have found the ventral attention system to be predominantly lateralized to the right hemisphere (Corbetta et al., 2008; Corbetta & Shulman, 2002; Mengotti, Käsbauer, Fink, & Vossel, 2020; Vossel et al., 2014). Similarly, the degree distribution of ventral attention nodes in graphs learned by CaLLTiF is strongly right-lateralized. We found the dorsal attention network to also be right-lateralized, but not as strongly as the ventral network. Similarly, The dorsal attention network is found by prior studies to be organized mostly bilaterally, except for specific regions (Intraparietal sulcus and frontal eye field) in the right dorsal network that show stronger involvement in the attentional control of both hemispheres compared to their left counterparts (Mengotti et al., 2020). Also similar is alignment in the lateralization of the default mode network, where both CaLLTiF and several past studies have found it to be left lateralized (Agcaoglu, Miller, Mayer, Hugdahl, & Calhoun, 2015; Banks et al., 2018; Nielsen, Zielinski, Ferguson, Lainhart, & Anderson, 2013; Swanson et al., 2011). Sensorimotor cortices, on the other hand, were found to be highly symmetric and not particularly lateralized to either hemisphere in causal graphs learned by CaLLTiF, a finding that is also consistent with the generally symmetric involvement of primary sensory and motor areas in contralateral processing (Agcaoglu et al., 2015). Finally, research on the laterality of the control and limbic networks is ongoing and, to the best of our knowledge, inconclusive (see, e.g., (Morton, 2020)). In graphs learned by CaLLTiF, however, we observe strong left lateralization of the limbic and right lateralization of the control networks, respectively. Thus, in summary, we observe clear laterality in all but sensorimotor cortical networks which either corroborate the existing literature or clarify previously inconclusive observations.

### Hyperparameter selection and sensitivity

CaLLTiF has two hyperparameters, *τ*_max_ and *α*_level_. Due to CaLLTiF’s type-I error control over lags (see Methods), these parameters are interdependent, as described by Equation (4). This makes *τ*_max_ the only effectively ‘free’ hyper-parameter, which can be systematically selected by analyzing how much each additional lag contributes to the final graph (as in Figure 6a). Furthermore, the end results of CaLLTiF are remarkably robust to variations in *τ*_max_. Supplementary Figure 59 illustrates the sensitivity of CaLLTiF by comparing graphs generated from real HCP data with *τ*_max_ = 1, 2, and 4 against those with *τ*_max_ = 3 analyzed in the main text. For comparison, we also show the percentage of changes in resulting graphs if we fix the value of *α*_level_ (i.e., ignore Equation (4)) so that, e.g., increasing *τ*_max_ from 3 to 4 only adds lag-4 edges without affecting other lags. As we can see, at *τ*_max_ = 2 and 4 (33% change in *τ*_max_), the resulting graphs change less than 6% for fixed type I error and less than 3% for fixed *α*_level_. Similarly, when changing *τ*_max_ to 1 (67% change in *τ*_max_), the resulting graphs change by less than 13% and 6% in the two conditions, respectively.

### Limitations

The present study has a number of limitations. From a biological perspective, synaptic causation happens at the level of neuronal activity, from which fMRI is a noisy readout. This lack of access to the true underlying neural activity limits the accuracy of any causal discovery method working with fMRI, and attempting to extract the underlying neural activity from fMRI data is often futile (Supplementary Figure 42, also cf. (Nozari et al., 2023)). The low temporal resolution of fMRI, even with the TR value of 720ms in the HCP data, also limits the precision of causal discovery. As we saw from Supplementary Figure 58, edges of all lengths are observed even at lag 0. This indicates the possibility that some of the edges discovered by CaLLTiF may be polysynaptic paths but resemble a direct monosynaptic connection at low temporal resolution. Finally, similar to most constraint-based methods, the causal graphs returned by CaLLTiF are not tied to a generative dynamical model (as is the case with VARLiNGAM, DYNOTEARS, DCM, etc). If such generative models are needed, VAR models based on CaLLTiF’s extended time-lagged graph constitute a natural choice, but further research is needed to compare the dynamic predictive accuracy of such models against potential alternatives (Ljung, 1999).

### Conclusions

Overall, this study demonstrates the interplay between the theoretical challenges of causal discovery and the practical limitations of fMRI, and the design of an algorithmic solution that can bridge this gap. This work motivates several follow-up studies, including the application of the proposed CaLLTiF method to task fMRI and comparing its outcomes against structural connectivity.

## MATERIAL AND METHODS

### Simulated fMRI Data

When comparing different causal discovery algorithms or different hyperparameters of the same algorithm, we used several benchmarks of simulated fMRI data with known ground truth connectivity from (Sanchez-Romero et al., 2019). In general, this dataset included two groups of networks, one consisting of 9 simple small-scale synthetic graphs and one consisting of two graphs extracted from the macaque connectome. From the latter group, we only used the smallest (Macaque_SmallDegree) and the largest (Macaque_Full).

The details of generating BOLD signals from each graph are detailed in (Sanchez-Romero et al., 2019). In brief, the same simulation procedure was used for simple and macaque-based graphs, where the authors used the model proposed in (Smith et al., 2011) which is itself based on the DCM architecture of (K. Friston et al., 2003). Underlying neural dynamics are simulated using the linear differential equation *dz/dt* = *σAz* + *Cu*, where *A* denotes the ground-truth connectivity. To simulate resting-state data, the *u* input was modeled using a Poisson process for each of the regions (*C* = *I*). The neuronal signals *z* were then passed through the Balloon-Windkessel model (Buxton, Wong, & Frank, 1998; Smith et al., 2011) to obtain simulated BOLD data.

### Resting-State fMRI from the Human Connectome Project

For the real fMRI analysis, we used ICA-FIX resting-state data from the Human Connectome Project S1200 release (Barch, 2017; Burgess et al., 2016; Essen et al., 2013). Resting-state fMRI images were collected with the following parameters: TR = 720 ms, TE = 33.1 ms, flip angle = 52 deg, FOV = 208x108 mm, matrix = 104x90, slice thickness = 2.0 mm, number of slices = 72 (2.0 mm isotropic), multi-factor band = 8, and echo spacing = 0.58 ms. Brains were normalized to fslr32k via the MSM-AII registration and the global signal was removed. We removed subjects from further analysis if any of their four resting state scans had excessively large head motion, defined by having frames greater than 0.2 mm frame-wise displacement or a derivative root mean square (DVARS) above 75. Also, subjects listed in (Elam, 2020) under “3T Functional Preprocessing Error of all 3T RL fMRI runs in 25 Subjects” or “Subjects without Field Maps for Structural scans” were removed. Among the remaining 700 subjects, the 200 with the smallest head motion (DVARS) were selected for analysis. For all subjects, we parcellated the brain into 100 cortical regions (Schaefer 100x7 atlas (Schaefer et al., 2018)) and 16 subcortical ones (Melbourne Scale I atlas (Tian, Margulies, Breakspear, & Zalesky, 2020)). The Human Connectome Project experiments were carried out by the WU-Minn consortium and its adherence to ethical standards was approved by the by the Internal Review Board of the respective institutions. Explicit informed consent was acquired from all participants involved in the study (Essen et al., 2013).

### Causal discovery methods

One aim of causal inference is to construct a causal graph based on observational data. The relationship between a probability distribution and its depiction as a graph plays a significant role in this process. Nevertheless, it is not always feasible to deduce a causal graph solely from observational data. Further assumptions are therefore required. Here, we briefly summarize the main assumptions and principles underlying the list of causal discovery methods studied in this work (cf. Table 1).

#### PCMCI

PCMCI was proposed in (Runge et al., 2019) as a constraint-based causal discovery method designed to work with time-series data. The algorithm is composed of two main steps. In the first step, the algorithm selects relevant variables using a variant of the undirected graph discovery part of the PC algorithm (Spirtes & Glymour, 1991). This step removes irrelevant variables for conditioning and therefore increases statistical power. In the second step, the algorithm uses the momentary conditional independence (MCI) test, which measures the independence of two variables conditioned on the set of their parents identified in step 1. The MCI test helps to reduce the false positive rate, even when the data is highly correlated. PCMCI assumes that the data is stationary, has time-lagged dependencies, and has causal sufficiency. Even when the stationarity assumption is violated, PCMCI was shown to perform better than Lasso regression or the PC algorithm (Runge et al., 2019). However, PCMCI is considered not suitable for highly predictable (almost deterministic) systems with little new information at each time step (Runge et al., 2019). The Python implementation of PCMCI is available in the Tigramite package at https://github.com/jakobrunge/tigramite.

As noted earlier, PCMCI only returns *◦−◦* edges among contemporaneous variables. While this allows PCMCI to relax the common DAG assumption and allow for cycles, it results in a mixed summary graph, where multiple types of edges (*→*, *→*, and/or *◦−◦*) can exist between two nodes. In contrast, we require all algorithms to output a directed graph. Therefore, when reporting F1 scores for PCMCI, we only include directed edges coming from lagged relationships and exclude the contemporaneous *◦−◦* edges. The only exception is what we call ‘Mixed PCMCI’ (See Supplementary Figures 28-30), where the contemporaneous *◦−◦* edges are also included in the computation of *adjacency* F1 scores.

#### PCMCI^+^

PCMCI^+^ is an extension of the PCMCI method which incorporates directed contemporaneous links in addition to the lagged ones (Runge, 2020). The approach revolves around two key concepts. First, it divides the undirected graph edge removal phase into separate lagged and contemporaneous conditioning phases, thereby reducing the number of conditional independence tests required. Second, it incorporates the idea of momentary conditional independence (MCI) tests from PCMCI (Runge et al., 2019) specifically in the contemporaneous conditioning phase. PCMCI^+^ also outputs a time-series graph with different types of contemporaneous edges, including directed edges (*→* and *→*), unoriented edges (*◦−◦*), and conflicting edges (*× − ×*). Consistent with our requirement of a regular digraph at the end, we disregarded the unoriented and conflicting edges and retained only the directed ones. Similar to most other causal discovery algorithms, PCMCI^+^ does not permit cycles in the contemporaneous links, which could potentially account for its relatively underwhelming performance over fMRI data. The Python implementation of PCMCI+ is also available in the Tigramite package https://github.com/jakobrunge/tigramite.

#### VARLiNGAM

VARLiNGAM is a causal discovery method that combines non-Gaussian instantaneous models with autoregressive models. This method, proposed in (Hyvärinen et al., 2010), builds on the fact that in the absence of unobserved confounders, linear non-Gaussian models can be identified without prior knowledge of the network structure. VARLiNGAM is capable of estimating both contemporaneous and lagged causal effects in models that belong to the class of structural vector autoregressive (SVAR) models and provides ways to assess the significance of the estimated causal relations. These models are a combination of structural equation models (SEM) and vector autoregressive (VAR) models. In addition, VARLiNGAM emphasizes the importance of considering contemporaneous influences, as neglecting them can lead to misleading interpretations of causal effects. Nevertheless, VARLiNGAM does not permit cycles in the contemporaneous links either, which could potentially account for its relatively poor performance over brain fMRI data with many feedback loops. The VARLiNGAM method is available from https://github.com/cdt15/lingam and a tutorial can be found at https://lingam.readthedocs.io/en/latest/tutorial/var.html.

#### DYNOTEARS

Dynamic NOTEARS (DYNOTEARS) method, proposed in (Pamfil et al., 2020), is a score-based method designed to discover causal relationships in dynamic data. It simultaneously estimates relationships between variables within a time slice and across different time slices by minimizing a penalized loss function while ensuring that the resulting directed graph is acyclic (including acyclicity of contemporaneous connections). The goal is to identify the best set of conditional dependencies that are consistent with the observed data. DYNOTEARS builds on the original NOTEARS method proposed in (Zheng, Aragam, Ravikumar, & Xing, 2018), which uses algebraic properties to characterize acyclicity in directed graphs for static data. Python implementations are available from the CausalNex library (https://github.com/quantumblacklabs/causalnex) as well as https://github.com/ckassaad/causal_discovery_for_time_series.

#### DGlearn

DGlearn is a score-based method for discovering causal relationships from observational data. Importantly, it is one of few algorithms that can learn cyclic structures from cross-sectional data. The method, introduced in (Ghassami et al., 2020), is based on a novel characterization of equivalence for potentially cyclic linear Gaussian directed graphical models. Two structures are considered equivalent if they can generate the same set of data distributions. DGlearn utilizes a greedy graph modification algorithm to return a graph within the equivalence class of the original data-generating structure. The Python implementation of DGlearn is available at https://github.com/syanga/dglearn.

#### FASK

The Fast Adjacency Skewness (FASK) method, proposed in (Sanchez-Romero et al., 2019), is a hybrid method for causal discovery from cross-sectional data, combining constraint-based and noise-based elements. It leverages (and needs) non-Gaussianity in the data and allows for cycles. This algorithm is composed of two main steps. The first step, called FAS-Stable, outputs an undirected graph *G*_0_ by iteratively performing conditional independence tests under the increasing size of the conditioning set and using the Bayesian information criterion (BIC) to compare the conditioning sets. In the second step, assuming i.i.d. non-Gaussian data, each of the *X − Y* adjacencies in *G*_0_ are oriented as a 2-cycle (⇆) if the difference between *corr*(*X, Y*) and *corr*(*X, Y |X >* 0), and *corr*(*X, Y*) and *corr*(*X, Y |Y >* 0), are both significantly nonzero, and as a unidirectional edge otherwise. The pseudo-code for FASK can be found in Supporting Information A of (Sanchez-Romero et al., 2019) and Java source code for it is available at http://github.com/cmu-phil/tetrad.

#### MVGC

In (Granger, 1969), Granger introduced a statistical version of Hume’s regularity theory, stating that *X_p_* Granger-causes *X_q_*, if past values of *X_p_* provide unique, statistically significant information about future values of *X_q_* (Assaad et al., 2022). While this allows for optimal forecasting of an effect and has been extended to multivariate systems (Barnett & Seth, 2014), MVGC cannot account for contemporaneous effects and the presence of unobserved confounders can result in spurious edges. Python implementation of MVGC is available at https://github.com/ckassaad/causal_discovery_for_time_series.

#### NTS-NOTEARS

NTS-NOTEARS is a nonlinear causal discovery method designed for time-series data (Sun, Liu, Poupart, & Schulte, 2021). It employs 1-D convolutional neural networks to capture various types of relationships, including linear, nonlinear, lagged, and contemporaneous connections among variables. The method ensures that the resulting causal structure forms a directed acyclic graph. It builds upon the NOTEARS approach (Zheng et al., 2018), and is similarly based on continuous optimization. Similar to other algorithms above, it assumes the presence of no hidden confounding factors and stationarity of the data-generating process. In our analysis, we compare NTS-NOTEARS as a state-of-the-art nonlinear method against the aforementioned linear algorithms in synthetic fMRI (cf. Supplementary Figure 38). A Python implementation of NTS-NOTEARS is available at https://github.com/xiangyu-sun-789/NTS-NOTEARS

#### CaLLTiF (proposed method)

The proposed CaLLTiF method builds upon PCMCI (Runge et al., 2019) but, instead of using a PC-type approach in the first step to estimate the set of parents for lagged variables, it starts from a complete conditioning set including all lagged variables. This choice dramatically decreases computational cost, but surprisingly, it is also optimal, as shown in Supplementary Figure 16, because as mentioned in the discussion section, the approach of PCMCI discards contemporaneous effects. Using a complete conditioning set, CaLLTiF then performs Momentary Conditional Independence (MCI) partial correlation tests between all pairs of variables. Specifically, for any pair *X_i_*(*t − τ*)*, X_j_*(*t*) with *i, j ∈* 1*, . . ., N* and time delays *τ ∈* 0, 1*, . . ., τ*_max_, a causal link is established (*X_i_*(*t − τ*) *→ X_j_*(*t*) if *τ >* 0 and *X_i_*(*t*)*◦−◦X_j_*(*t*) if *τ* = 0), if and only if:

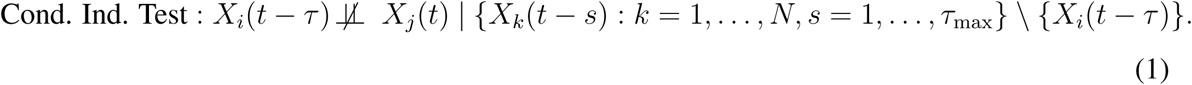

Note that, despite being complete, the conditioning sets only include variables from *prior* time lags. As noted earlier, to test a conditional independence of the form *X ⊥̸⊥ Y |Z*, we compute the partial correlation coefficient *ρ*(*X, Y |Z*) between *X* and *Y* conditioned on the set of variables in *Z* and the corresponding p-value for the null hypothesis that *ρ*(*X, Y |Z*) = 0. An edge is placed between *X_j_*(*t*) and *X_i_*(*t − τ*) if this p-value is less than the hyperparameter ‘Alpha Level’. The value of this threshold was selected optimally in simulated fMRI and using temporal correction for multiple comparisons (see below) in real data. Finally, for contemporaneous pairs (*τ* = 0), each *◦−◦* edge is replaced with ⇆ if there are no other edges between those two variables at other lags, and is replaced with a directed edge or a ⇆ based on the lagged direction(s) otherwise. For a more detailed summary of the partial correlation-based edge discovery in CaLLTiF, see Supplementary Note 2. A pseudocode of CaLLTiF is shown in Algorithm 1.

##### Algorithm 1

Causal discovery for Large-scale Low-resolution Time-series with Feedback (CaLLTiF)

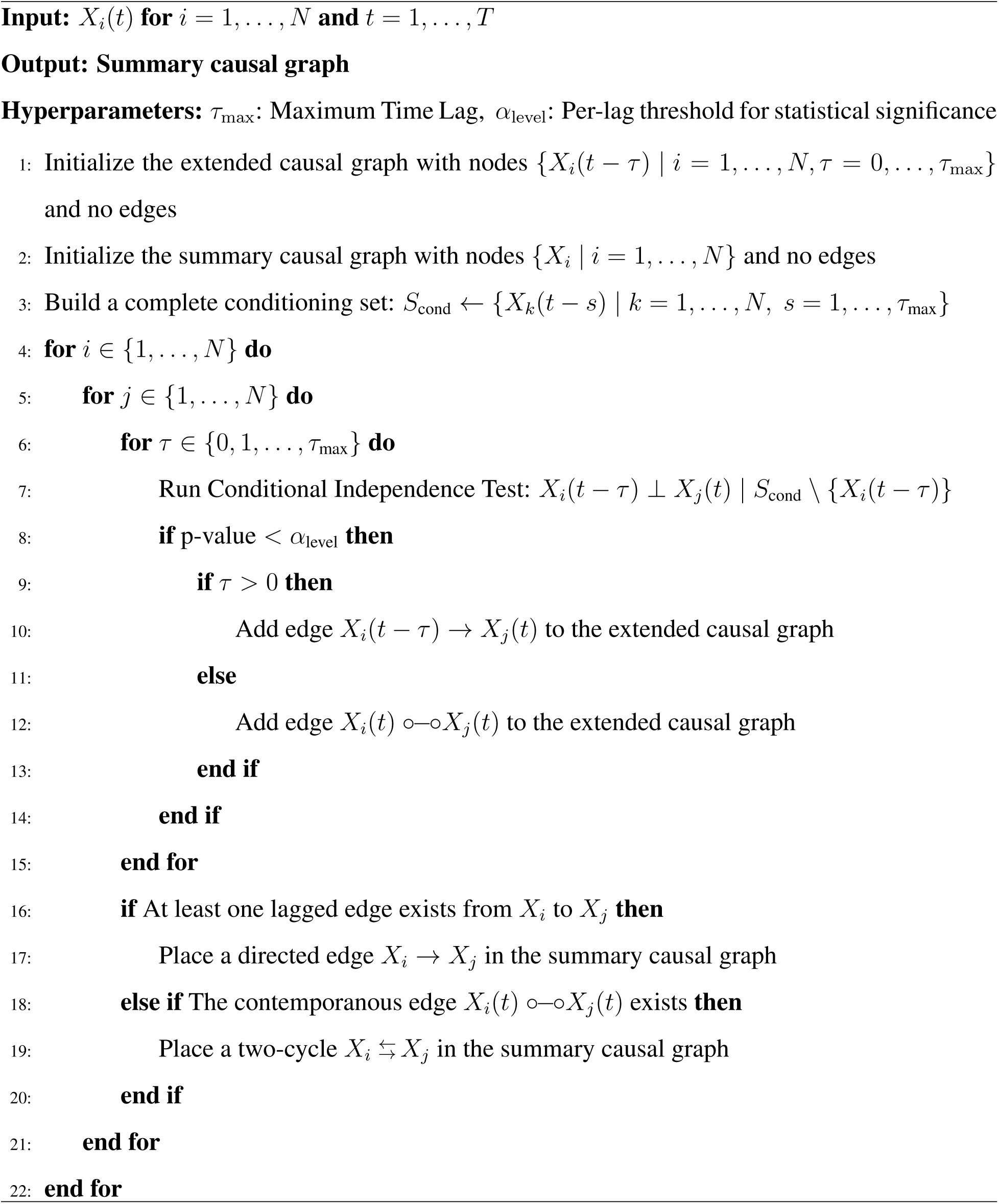

Finally, it is imperative to acknowledge the possibility that some of the directed edges detected by our methodology do not possess a strictly causal connotation. As previously indicated, the orientation method relies on the widely accepted premise that bidirectional connections hold notably greater prevalence than unidirectional links. Thus, we believe that the presented approach shall yield a proximate representation of the true causal graph, concurrently accommodating cyclic structures and circumventing computational barriers.

### Construction of summary causal graphs from causal graphs over lagged variables

Causal discovery algorithms designed for time series data often return a causal graph among the lagged variables

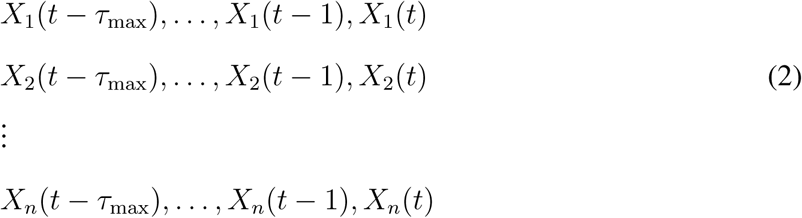

For algorithms other than CaLLTiF, from this we extract a final *summary* graph among the variables *X*_1_*, . . ., X_n_* by placing an edge from *X_i_* to *X_j_* if there exists any *τ ≥* 0 for which there is an edge from *X_i_*(*t − τ*) to *X_j_*(*t*). This is equivalent to an OR operation among binary edges (as opposed, e.g., to a majority vote) and must be taken into account when interpreting the obtained summary graphs. The process is similar in CaLLTiF except that the direction(s) of contemporaneous *◦−◦* edges are first resolved using lagged edges before executing the OR across lags (cf. Algorithm 1).

### Correction for multiple comparisons across lags in CaLLTiF

As noted above, CaLLTiF places an edge from *X_i_* to *X_j_* in its summary graph if there exists at least one *τ ≥* 0 for which there is an edge from *X_i_*(*t − τ*) to *X_j_*(*t*). Therefore, the decision to place an edge from *X_i_* to *X_j_* depends on the outcomes of *τ*_max_ + 1 statistical tests, and to maintain a desired bound on the probability of type I error for each edge in the *summary* graph, we need to account for multiple comparisons across lags.

Formally, for each edge *X_i_ → X_j_* in the final graph, the null hypothesis (i.e., lack of a direct causal effect from *X_i_* to *X_j_*) can be formulated as

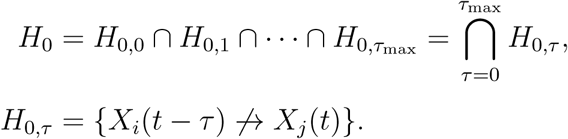

Let *p_τ_* denote the p-value of the partial correlation test between *X_i_*(*t − τ*) and *X_j_*(*t*) and *α*_level_ denote the significance threshold for each partial correlation test. Then, the probability of type I error is

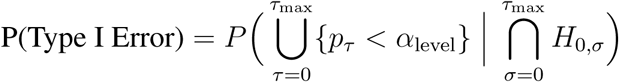

Note that this is different from the family-wise error rate (FWER, bounded by the Bonferroni method and its extensions) or the false discovery rate (FDR). In particular, this is different from FWER in that only one decision is made and the probability of error is computed for that single decision only. So, for instance, if in reality any subset (even one) of *{H*_0_*_,τ_ }* is false and the algorithm rejects any subset (even all) of *{H*_0_*_,τ_ }*, there is no type I error, as an edge exists from *X_i_* to *X_j_* both in the data-generating process and in the final summary graph.

The type I error can then be bounded as

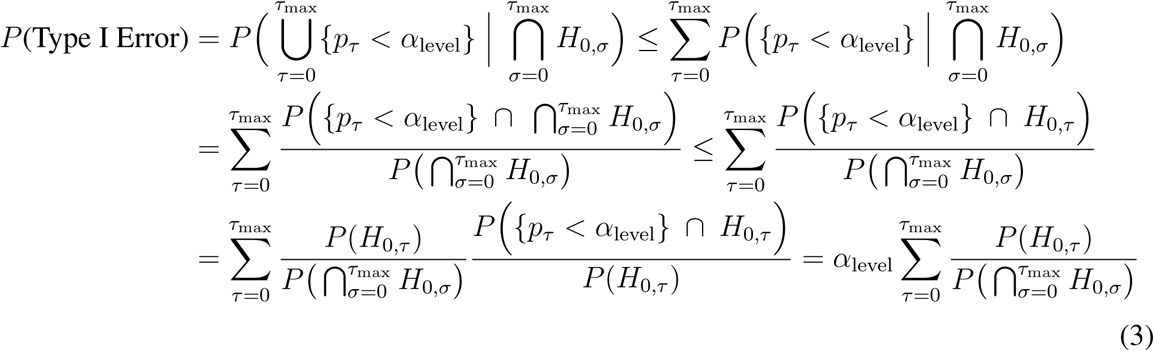

The last expression has no dependence on the data and depends only on the prior distribution we consider on graphs. Assuming a uniform prior, *P* (*H*_0_*_,τ_*) = 1*/*2. Further,

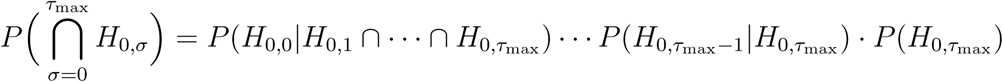

We assume a prior where knowledge of the lack of an edge from *X_i_* to *X_j_* at one lag either increases the probability of lack of an edge between them at other lags or, at least, does not decrease it (independence across lags). Then,

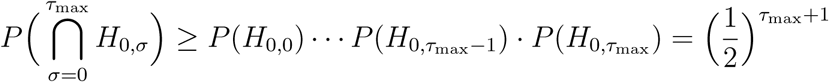

Putting everything together, we get

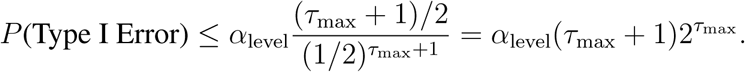

Note, for analogy, that the correction factor (*τ*_max_ + 1)2*^τ^*^max^ takes place of the factor (*τ*_max_ + 1) in a corresponding Bonferroni correction. To have *P* (Type I Error) less than a prescribed threshold *α*, we choose

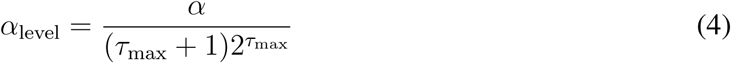

In our experiments with the HCP data, we have *τ*_max_ = 3 and *α* = 0.01, giving a per-lag significance threshold of *α*_level_ *≃* 0.0003. This is notably smaller than the Alpha Level values that maximized F1 scores in simulated Macaque_Full data (0.1 for adjacency F1 score and 0.01 for F1 score), and is due to the conservative nature of this correction for temporal multiple comparisons.

### Computing Functional Graphs

In order to calculate the functional graphs for each subject, we consolidated the data from the four sessions of each subject in the HCP and computed the pairwise correlations among all pairs of parcels. To form a binary functional graph, we placed an edge between any two parcels displaying a statistically significant correlation coefficient (*p <* 0.01, t-test for Pearson correlation coefficient).

### Hyperparameter Selection

All the methods we described in Table 1 have at least one main hyperparameter that significantly affects the end result, particularly in terms of edge density. These include ‘PC Alpha’ and ‘Alpha Level’ for PCMCI, ‘PC Alpha’ for PCMCI^+^, ‘Alpha’ for VARLINGAM, DYNOTEARS, and MVGC, and FASK, and ‘BIC Coefficient’ for DGlearn. These hyperparameters were swept over (simultaneously for PCMCI) using the simulated data and selected such that the F1 score with the ground truth graph is maximized in each case. This process was repeated for all algorithms and all experiments (simple graphs, Macaque_SmallDegree, Macaque_Full). Performance metrics such as Recall, Precision, and F1 scores of each method for a range of their hyperparameters are shown in Supplementary Figures 6-12 for the simulated Simple Network graphs, in Supplementary Figures 20-24 for the simulated Macaque_SmallDegree data, and in Supplementary Figures 31-38 for the simulated Macaque_Full data.

Time-series algorithms (PCMCI, PCMCI^+^, VARLiNGAM, DYNOTEARS) also have a hyperparameter controlling the number of lags used for causal discovery. Based on our prior work (Nozari et al., 2023), we set this value to 3 for the HCP data (TR = 0.72s), and confirmed its sufficiency based on the contributions of higher-order lags (Figure 6a). For the simulated data, (TR = 1.2s), we used a maximum lag of 2 to match its slower sampling.

### Computing F1 Scores, Degrees, and Causal Flows

In our experiments using simulated fMRI data, access to ground truth graphs allows for evaluating the performance of causal discovery methods. In this work, we evaluate causal discovery algorithms as binary classifiers deciding the presence or lack of *n*^2^ edges among *n* nodes. This allows us to evaluate algorithms using standard classification metrics such as precision, recall, and F1 score (Davis & Goadrich, 2006; Fawcett, 2006; Powers, 2020; Sokolova & Lapalme, 2009; Tharwat, 2020). Given that the F1 score provides a balanced trade-off between precision and recall, we use it as our measure of accuracy. We define two separate metrics, (full) F1 score and adjacency F1 score. For the former, each of the *n*^2^ edges (including any self-loops due to dampening autocorrelation for each node) in the graph is considered as one test sample for classification. In the latter, the ground-truth and learned graphs are first transformed into an undirected graph, placing an edge between two nodes if a directed edge existed in at least one direction. The resulting 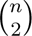 possible edges are then treated as test samples for classification and computation of adjacency F1 score.

To determine the degree and causal flow of a node *i* in a *binary* directed graph, its in-degree (number of edges pointing toward node *i*) and out-degree (number of edges originating from node *i*) are first computed and normalized by the total number of nodes in the graph. The degree of node *i* is then computed as the sum of the out-degree and in-degree, while the causal flow is obtained by subtracting the in-degree from the out-degree. The same process is followed for weighted graphs except that the calculation of in-degree and out-degree involves a weighted mean. Mathematically,

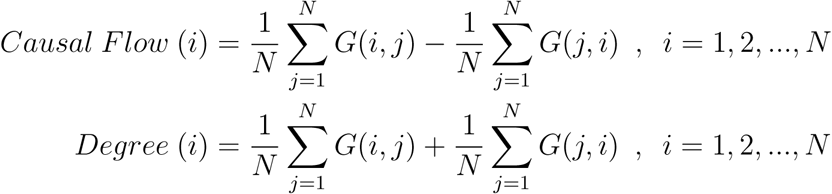

where *G* denotes the graph’s (binary or weighted) adjacency matrix.

### Computing Subnetwork Graphs from Parcel-Level Graphs

Subnetwork graphs were computed by aggregating parcel-level binary graphs into graphs between 16 subnetworks. These subnetworks consist of the standard 7 resting-state subnetworks (Yeo et al., 2011) plus one subcortical subnetwork, separately for the left and right hemispheres. A subnetwork-level graph is then computed for each subject, whereby the weight of an edge from subnetwork *i* to *j* is the number of nodes in subnetwork *i* that connect to nodes in subnetwork *j*, normalized by the number of all possible edges between these subnetworks. The results are then averaged over the subject, as depicted in Supplementary Figure 51.

### Computing

All the computations reported in this study were performed on a Lenovo P620 workstation with AMD 3970X 32-Core processor, Nvidia GeForce RTX 2080 GPU, and 512GB of RAM.

## Supporting information

Supplementary Material for "Whole-Brain Causal Discovery Using fMRI"

## SUPPORTING INFORMATION

All the fMRI data used in this work is publicly available. The simulated fMRI benchmarks can be downloaded from https://github.com/cabal-cmu/Feedback-Discovery and the human fMRI data can be accessed via the HCP S1200 Release at https://www.humanconnectome.org/study/hcp-young-adult/document/1200-subjects-data-release. The Python code for this study is publicly available at https://github.com/nozarilab/2023Arab_CaLLTiF.

## AUTHOR CONTRIBUTIONS

EN and AG designed and supervised the study; FA performed the research; HJ and MAKP assisted in the analyses of human fMRI data; FA and EN drafted and all authors edited the manuscript.

## ACKNOWLEDGMENTS

The research conducted in this study was partially supported by NSF award no. 2239654 (to EN), the Canadian Institute for Advanced Research (fellowship awarded to MAKP), and the Air Force Office of Scientific Research under award no. FA9550-20-1-0106 (to MAKP).

## TECHNICAL TERMS

Causal Discovery: The process of identifying causal relationships between variables from observational data, namely, determining how changing the value of each variable causally influences others.
Contemporaneous Causal Effect: A causal relationship occurring within the same observation interval when the underlying causal processes are faster than the rate of sampling.
Causal Flow: The net difference between outgoing and incoming edges in a causal graph, indicating whether a node is a source (positive causal flow) or a sink (negative causal flow).
Partial Correlation: A statistical measure assessing the direct linear association between two variables after regressing out the effects of a set of conditioned variables.
Conditional Independence Test: A statistical test that evaluates whether two variables are independent after conditioning on the influence of one or more other variables. Partial correlation is a common method for testing conditional independence in linear models.
Type I Error: In the context of binary decisions, the incorrect rejection of a true null hypothesis, a.k.a. a false positive.
Correction for Multiple Comparisons: The process of adjusting statistical tests to reduce the risk of Type I errors when multiple hypotheses are tested simultaneously.
F1 Score: A balanced performance metric for binary classification, combining precision and recall into a single value that penalizes both false positives and false negatives.
Subnetwork-Level Graph: A low-dimensional graph in which nodes represent functionally-clustered subnetworks of parcels and edges represent average connectivity between parcels in two subnetworks.
Hemispheric Laterality: The asymmetry in the distribution of a variable between the two brain hemispheres.

