## Supplementary Material for "Whole-Brain Causal Discovery Using fMRI" for "Whole-Brain Causal Discovery Using fMRI"

### Supplementary Notes

**1 Contemporaneous Causal Effects** **2**

**2 Distinctions Between CaLLTiF and MVGC** **4**

**3 On the Rationale for Only Conditioning on Past Values in CaLLTiF** **6**

**4 Computational Complexity of CaLLTiF** **7**

### 1 Contemporaneous Causal Effects

Contemporaneous causal connections play an important role in CaLLTiF and are one of its main advantages over several alternative algorithms such as MVGC and DCM. In this Supplementary Note, we define what we mean by a contemporaneous edge, explain why they arise as a result of temporal under-sampling, and provide further details on their processing and importance in CaLLTiF.

Unlike what the name may suggest, a contemporaneous edge does *not* mean or imply an instantaneous causal effect. In most physical systems, and certainly in the brain, causation takes time. However, ground-truth causation can occur at a timescale significantly faster than the sampling timescale of observations. Spike generation, axonal conduction, and synaptic transmission which constitute the backbone of causal flow in the brain, for example, take on the order of $10^1$-$10^2$ms while fMRI sampling using state-of-the-art scanners takes on the order of $10^3$ms. This mismatch between timescales effectively generates hidden variables, namely, the values of all of $X_1(t), \ldots, X_n(t)$ in between sampling times–e.g., $X_1(0.5)$, $X_2(1.2)$, etc.

This effect is illustrated in an example in Supplementary Figure 1. For the sake of simplicity, consider only 3 nodes and a ratio of 2-to-1 between the observation timescale and the ground-truth causation timescale. The variables $X_i(t-1/2), i = 1, 2, 3$ are all latent variables that can cause the observed variables $X_i(t), i = 1, 2, 3$ but, due to being latent, cannot be conditioned on. For instance, $X_1(t)$ and $X_2(t)$ are both caused by $X_2(t-1/2)$, generating a statistically significant correlation between them that cannot be resolved by conditioning on $X_2(t-1/2)$. Theoretically, this latent-confounded statistical dependence is what we define as a *contemporaneous edge* in CaLLTiF, which is the same as what is often shown by a bidirectional edge in an acyclic directed mixed graph (ADMG) (Richardson et al., 2023).

The meaning of a lagged edge in CaLLTiF should also be understood in the same context where slow sampling leads to latent nodes in between sampling times. Assuming that no direct (mono-synaptic) causation can take nearly as long as one TR (720ms in HCP and often longer in most fMRI recordings), a lagged edge in CaLLTiF corresponds to a directed *path* in the ground-truth causal graph in which all the intermediate nodes are latent. This is also the same as what is marked by a directed edge between measured variables in an ADMG. As an example, in Supplementary Figure 1, the lagged edge from $X_2(t-1)$ to $X_1(t)$ in the middle panel captures the directed path $X_2(t-1) \to X_2(t-1/2) \to X_1(t)$ in the left panel. Similarly, all other lagged edges in the middle panel correspond to directed, latent-mediate paths through $X_1(t-1/2)$, $X_2(t-1/2)$, or $X_3(t-1/2)$.

In its last step, CaLLTiF incorporates a heuristic step for summarization across lags, as shown in the right panel of Supplementary Figure 1. When no contemporaneous edge exists between a pair of variables $X_i$ and $X_j$, this step is simply a disjunction (OR) across the lagged edges. However, when a contemporaneous edge exists, two scenarios may happen: (i) one or more lagged edges also exist, or (ii) no lagged edges exist between that pair of variables. In (i), we use the direction of the lagged edges to disambiguate the (intrinsically undirected) contemporaneous edge and then summarize across all lagged and contemporaneous

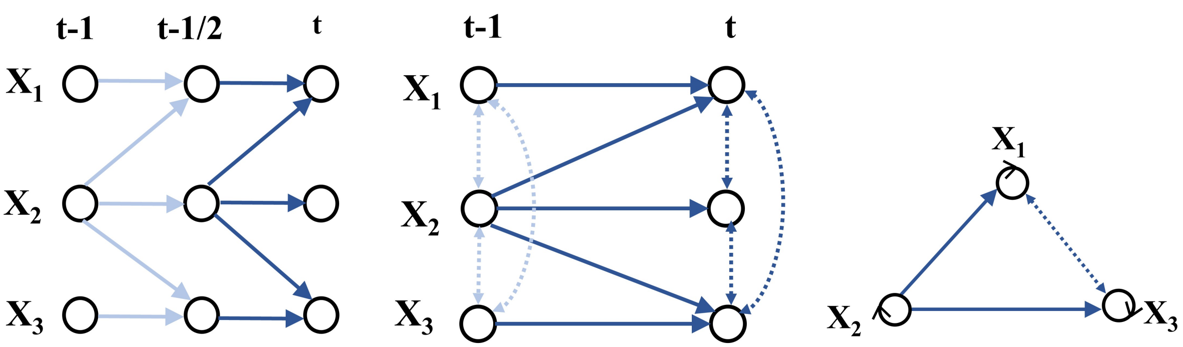

**Supplementary Figure 1: Example showing how contemporaneous causal links are generated in CaLLTiF and how the final summary graph is constructed based on that.** (a) Ground truth window causal graph in the true causal timescale (b) Corresponding window causal graph estimated by CaLLTiF (c) Corresponding summary causal graph outputted by CaLLTiF

edges using disjunction. While this can be seen as effectively ignoring the contemporaneous edge in building the summary graph, it can be very helpful in avoiding spurious two-cycles when only a uni-directional effect exists. This is the case, for example, between $X_2$ and $X_1$ (as well as $X_2$ and $X_3$) in Supplementary Figure 1. The lagged edge from $X_2$ to $X_1$ (and $X_3$) disambiguates the direction of the contemporaneous effect between them, leading to a correct unidirectional edge in the summary graph. In fact, it is straightforward to show that when all nodes have persistent auto-causation (the horizontal chains in the left panel of Supplementary Figure 1), the lack of any lagged edge from a node $X_i$ to any other node $X_j$ necessarily implies the lack of any direct causal effect from $X_i$ to $X_j$. In other words, if lagged edges are only present in one direction, not putting an edge in the opposite direction in the summary graph is always a correct decision. Finally, if no such lagged edges are present to disambiguate the direction of the contemporaneous effect between a pair of variables (case (ii)), then a two-cycle is placed between the two variables in the summary graph which is the most likely prior for a pair of nodes in brain networks (Tigges et al., 1973; Felleman and Van Essen, 1991; Markov et al., 2014).

A note is here warranted on why we need contemporaneous edges at all. Theoretically, if both a contemporaneous edge and one or more lagged edges exist between a pair of variables (case (i)), then the contemporaneous edge is (and should be) effectively ignored, as described above. On the other hand, if only a contemporaneous edge exists between two nodes $X_i$ and $X_j$ (case (ii)), it can be shown under the same assumption of persistent auto-causation above that the contemporaneous edge can only be due to a latent common cause $X_k(t-\tau)$ where $k \neq i, j$ and $t-1 < \tau < t$. In other words, the contemporaneous edge is spurious in this case. An example of this can be seen between $X_1$ and $X_3$ in Supplementary Figure 1. As such, in both cases, the contemporaneous effect seems either ineffective or wrong. This is true, however, only in theory when one has sufficient data, computational power, and knowledge of the exact parametric forms of the causal relationships between all variables to detect any and all lagged effects, no matter how weak or noisy they may be.

In reality, however, noise is constantly injected at every time point and statistical correlations between variables at time $t-1$ and those at time $t$ decay exponentially with the number of unobserved intermediate steps between them. As a result, with the typical number of data samples and timescale differences in fMRI, causal relationships between a vast majority of pairs of variables can only be detected contemporaneously. This can be clearly seen from Figure 6a in the main text, where about %70 of detectable effects are contemporaneous only. This is the case even with the "fast" sampling time of 0.72s in HCP data. In most other state-of-the-art fMRI recordings, sampling times are between 1-3s, making contemporaneous effects almost all that remain detectable.

### 2 Distinctions Between CaLLTiF and MVGC

Similar to MVGC, CaLLTiF detects the presence of an edge using multivariate linear autoregressive models and statistical hypothesis testing to determine statistical significance. In this Supplementary Note, we discuss their differences in further detail and explain why CaLLTiF can achieve better accuracy in large networks, such as the simulated Macaque_Full and HCP datasets we examined in this work. For ease of reference and uniformity of notation, we first revisit the underlying statistical tests used by each algorithm.

#### 2.1 Statistical test for determining the presence of an edge in CaLLTiF

**Computing the test statistic (partial correlation coefficient).** To determine the presence of an edge between $X_i(t-\tau)$ and $X_j(t)$ in CaLLTiF, we first regress out the effects of all potential common causes and mediators from both variables. This is done by constructing two linear regression models, one that predicts $X_j(t)$ and another that predicts $X_i(t-\tau)$ from all potential common causes and mediators.

$$X_i(t-\tau) = \sum_{\substack{k\in\{1,\ldots,N\} \\ s\in\{1,\ldots,\tau_{\max}\} \\ (k,s)\neq(i,\tau)}} \alpha_{k,s} X_k(t-s) + E_{X_i}(t), \qquad t = \tau_{\max}, \ldots, T-1 \tag{S1a}$$

$$X_j(t) = \sum_{\substack{k\in\{1,\ldots,N\} \\ s\in\{1,\ldots,\tau_{\max}\} \\ (k,s)\neq(i,\tau)}} \beta_{k,s} X_k(t-s) + E_{X_j}(t), \qquad t = \tau_{\max}, \ldots, T-1 \tag{S1b}$$

The regression coefficients $\{\alpha_{k,s}\}$ and $\{\beta_{k,s}\}$ are fit using standard least squares, the residuals of each model are calculated,

$$\hat{E}_{X_i}(t) = X_i(t-\tau) - \sum_{\substack{k\in\{1,\ldots,N\} \\ s\in\{1,\ldots,\tau_{\max}\} \\ (k,s)\neq(i,\tau)}} \hat{\alpha}_{k,s} X_k(t-s), \qquad t = \tau_{\max}, \ldots, T-1 \tag{S2a}$$

$$\hat{E}_{X_j}(t) = X_j(t) - \sum_{\substack{k\in\{1,\ldots,N\} \\ s\in\{1,\ldots,\tau_{\max}\} \\ (k,s)\neq(i,\tau)}} \hat{\beta}_{k,s} X_k(t-s), \qquad t = \tau_{\max}, \ldots, T-1 \tag{S2b}$$

and the Pearson correlation coefficient is calculated between $\hat{E}_{X_i}(t)$ and $\hat{E}_{X_j}(t)$. Let $r_{i,j,\tau}$ denote this correlation coefficient, a.k.a. the partial correlation coefficient between $X_i(t-\tau)$ and $X_j(t)$ conditioned on $\{X_k(t-s) : (k,s) \neq (i,\tau)\}$.

**Hypothesis testing (t-test).** We place an edge between $X_i(t-\tau)$ and $X_j(t)$ if the partial correlation coefficient $r_{i,j,\tau}$ is statistically significantly different from 0. This is done using the standard t-test for Pearson correlation coefficient, namely, with the null hypothesis $H_0 : r_{i,j,\tau} = 0$ and the t statistic

$$t = \begin{cases} \frac{r_{i,j,\tau}}{\sqrt{1-r_{i,j,\tau}^2}} \sqrt{T - N\tau_{\max} - 2} & \text{if} \quad \tau = 0 \\ \frac{r_{i,j,\tau}}{\sqrt{1-r_{i,j,\tau}^2}} \sqrt{T - N\tau_{\max} - 1} & \text{if} \quad \tau > 0 \end{cases}$$

where $T$ is the number of observations, $N$ is the number of variables, and $\tau_{\max}$ is the number of time lags.

#### 2.2 Statistical test for determining the presence of an edge in MVGC

**Computing the test statistic (conditional Granger causality index, CGCI).** To determine the presence of an edge from $X_i$ to $X_j$, MVGC also fits two linear regression models (note the lack of time indexing, as MVGC learns the summary graph directly instead of learning an extended window graph and summarizing it as done in CaLLTiF). A "full model" predicts $X_j(t)$ using the past history of all the variables, while a "restricted model" predicts $X_j(t)$ using the past history of all other variables but $X_i$:

$$\text{Full model:} \qquad X_j(t) = \sum_{\substack{k\in\{1,\dots,N\}\\ s\in\{1,\dots,\tau_{\max}\}}} \alpha_{k,s} X_k(t-s) + E_{X_j}(t), \qquad t = \tau_{\max},\dots,T-1 \tag{S3a}$$

$$\text{Restricted model:} \qquad X_j(t) = \sum_{\substack{k\in\{1,\dots,N\}\setminus\{i\}\\ s\in\{1,\dots,\tau_{\max}\}}} \beta_{k,s} X_k(t-s) + E_{X_j\setminus X_i}(t), \qquad t = \tau_{\max},\dots,T-1 \tag{S3b}$$

Estimated residuals $\hat{E}_{X_j}$ and $\hat{E}_{X_j\setminus X_i}$ are then computed similar to Eq. (S2) and the CGCI from $X_i$ to $X_j$ is defined as:

$$CGCI_{X_i\to X_j} = \ln\left(\frac{\hat{\sigma}_R^2}{\hat{\sigma}_F^2}\right)$$

where $\hat{\sigma}_F^2$ and $\hat{\sigma}_R^2$ are the variances of $\hat{E}_{X_j}$ and $\hat{E}_{X_j\setminus X_i}$, respectively.

**Hypothesis testing (F-test).** The statistical significance of the CGCI is tested using an F-test with the statistic

$$F = \frac{(\hat{\sigma}_R^2 - \hat{\sigma}_F^2)/\tau_{\max}}{\hat{\sigma}_F^2/(T-\tau_{\max}-N\tau_{\max})} = \frac{T-\tau_{\max}-N\tau_{\max}}{\tau_{\max}}\left(e^{CGCI_{X_i\to X_j}}-1\right)$$

where $T$ is the number of observations and $N\tau_{\max}$ is the number of coefficients in the full model.

#### 2.3 Comparison between independence test in MVGC and CaLLTiF

The strongest similarity between MVGC and CaLLTiF comes from their underlying linear regression models in Eq. (S3) and Eq. (S1). Nevertheless, there are major differences in how these regression models are used and how their statistical significance is assessed in each algorithm. In particular, partial correlation measures the *remaining association* between two variables *after having removed* the effect of all others. On the other hand, CGCI measures the *added predictive power* of the history of one variable in explaining the other, while all other variables are kept as regressors in a multivariate system. In other words, both algorithms are based on how much one variable ($X_i$) can explain variance in another ($X_j$), but CaLLTiF assesses the significance of this relationship *per se* (in an *absolute* sense), while MVGC compares the additional explained variance coming from $X_i$ with how much variance all other variables were unable to explain (in a *relative* sense).

This difference between how regression models are used in MVGC and CaLLTiF can lead to a higher sensitivity to the total number of variables in MVGC. In our experiments both with simulated and real data ($N \sim 100$), we indeed found MVGC to generate overly sparse graphs for standard significance thresholds ($\alpha \leq 0.05$) and unreliable/overly dense graphs for higher thresholds. Specifically, for the Macaque_Full simulated dataset, we swept over the significance threshold of MVGC (like we did for all other algorithms) and found its highest F1 score (as shown in Figure 3 in the main text). This was achieved at $\alpha = 0.5$, i.e., when the algorithm returns a complete graph. In other words, any value of $\alpha$ lower than 0.5 led to such inaccurate removal of edges that the result was worse than the complete graph. Likewise, in our experiments with the HCP data, even though we can no longer compute or optimize the ground-truth accuracy of MVGC over its significance threshold, we ran MVGC at the same significance level of 0.01 used for CaLLTiF. The resulting graphs, however, were completely empty. This is even despite the fact that CaLLTiF actually uses a much smaller significance threshold (0.01/32) for each t-test due to its conservative correction for multiple comparisons across lags. In other words, while CaLLTiF can find hundreds of edges with %99 or higher statistical confidence, MVGC can find none. One can certainly increase the significance threshold and have MVGC detect more edges, but at the risk of increased false positives (type I error) when there is no ground-truth to compare the results against.

### 3 On the Rationale for Only Conditioning on Past Values in CaLLTiF

A key element in CaLLTiF (and constraint-based methods for causal discovery in general) that allows for distinguishing between correlation and causation is conditioning on variables that can potentially be a common cause or mediator. The fork ($X_1 \leftarrow X_2 \rightarrow X_3$) and chain ($X_1 \rightarrow X_2 \rightarrow X_3$) architectures are the simplest network motifs that showcase the effect of a common cause and mediator, respectively. In the fork motif, $X_1$ and $X_3$ are correlated but neither causes the other–their correlation is due to their common cause $X_2$. Assuming linear Gaussian generative mechanisms, it is straightforward to show that the Pearson correlation coefficient between $X_1$ and $X_3$ is statistically significantly nonzero, while their partial correlation coefficient after conditioning on $X_2$ is zero on average (finite samples) and asymptotically (infinite samples). Similarly in the chain motif, conditioning on $X_2$ makes $X_1$ and $X_3$ statistically independent, as desired.

Nevertheless, conditioning is not universally beneficial. Excessive conditioning can also generate spurious correlations (and therefore edges) between variables that are otherwise not correlated. The simplest network motif showcasing this effect is the collider, a.k.a. V-structure ($X_1 \rightarrow X_2 \leftarrow X_3$). In this network, $X_1$ and $X_3$ are independent per se, but they become dependent if one conditions on their common child $X_2$. Intuitively, knowing the value of $X_2$ can make certain combinations of $X_1$ and $X_3$ more or less likely to have occurred than their default, thus coupling their distributions. This is the reason why constraint-based methods for causal discovery, such as the celebrated PC algorithm (Spirtes and Glymour, 1991) and its variants (including PCMCI), spend significant computational resources on selecting the best conditioning set for each pair of variables.

In this work, we showed that conditioning on all past values of all recorded variables is not only computationally vital for scalability but also most accurate in simulated fMRI (Figure 16 in the main text). Nevertheless, given the importance of contemporaneous effects discussed here, one may wonder why conditioning is restricted to lagged variables ($s > 0$ in Eq. (S1)). In the following, we provide empirical evidence for why conditioning on contemporaneous variables can generate more spurious edges than it can prune, particularly in certain parametric regimes.

We generated linear Gaussian time series from the structural equation models (SEMs) of two network architectures: the chain ($X_1(t) \rightarrow X_2(t) \rightarrow X_3(t)$) and the collider ($X_1(t) \rightarrow X_2(t) \leftarrow X_3(t)$). Auto-causation was added for all variables to generate temporal auto-correlations–the strongest source of dynamics in neural data. For the chain network, the SEM takes the form

$$X_1(t) = \alpha X_1(t-1) + e_1(t) \tag{S4a}$$

$$X_2(t) = \alpha X_2(t-1) + \beta X_1(t) + e_2(t) \tag{S4b}$$

$$X_3(t) = \alpha X_3(t-1) + \beta X_2(t) + e_3(t) \tag{S4c}$$

while for the collider, the SEM reads

$$X_1(t) = \alpha X_1(t-1) + e_1(t) \tag{S5a}$$

$$X_2(t) = \alpha X_2(t-1) + \beta X_1(t) + \beta X_3(t) + e_2(t) \tag{S5b}$$

$$X_3(t) = \alpha X_3(t-1) + e_3(t) \tag{S5c}$$

All auto- and cross-causation weights are set uniformly at $\alpha$ and $\beta$, respectively, to simplify parametric analyses.

We then investigated the impact of conditioning solely on past values of $X_2$ compared to conditioning on both its past and present values on the partial correlation between $X_1(t)$ and $X_3(t)$. For the chain, we would ideally like conditioning to *maximally reduce* correlation between $X_1(t)$ and $X_3(t)$ compared to their unconditional (baseline) correlation coefficient. In contrast, for the collider, we ideally want conditioning *not to increase* the correlation between $X_1(t)$ and $X_3(t)$ compared to their unconditional baseline. Therefore, we compare three correlation coefficients for each network motif: unconditional correlation ($R_{X_1(t),X_3(t)}$), partial correlation conditioning only on the past ($R_{X_1(t),X_3(t)|X_2(t-1),...,X_2(t-\tau_{\max})}$), and partial correlation conditioning on past and present ($R_{X_1(t),X_3(t)|X_2(t),X_2(t-1),...,X_2(t-\tau_{\max})}$).

Not surprisingly, $R_{X_1(t),X_3(t)|X_2(t-1),...,X_2(t-\tau_{\max})}$ almost always falls in between the other two, i.e., both the desired and undesired effects of conditioning only on the past are smaller compared to conditioning on past and present. Therefore, to assess whether conditioning only on the past (as done on CaLLTiF) is better than conditioning on the past and present overall, we computed the following metric for each network motif:

$$D = \frac{R_{X_1(t),X_3(t)|X_2(t-1),...,X_2(t-\tau_{\max})} - R_{X_1(t),X_3(t)}}{R_{X_1(t),X_3(t)|X_2(t),X_2(t-1),...,X_2(t-\tau_{\max})} - R_{X_1(t),X_3(t)}} \tag{S6}$$

This is simply a linear mapping of the interval between $R_{X_1(t),X_3(t)}$ and $R_{X_1(t),X_3(t)|X_2(t),X_2(t-1),...,X_2(t-\tau_{\max})}$ to $[0, 1]$, such that

- if $R_{X_1(t),X_3(t)|X_2(t-1),...,X_2(t-\tau_{\max})} = R_{X_1(t),X_3(t)}$ then $D = 0$, and
- if $R_{X_1(t),X_3(t)|X_2(t-1),...,X_2(t-\tau_{\max})} = R_{X_1(t),X_3(t)|X_2(t),X_2(t-1),...,X_2(t-\tau_{\max})}$ then $D = 1$.

Ideally, we would like $D = 1$ for the chain and $D = 0$ for the collider. In practice, conditioning on the past is better whenever $D_{\text{chain}} > D_{\text{collider}}$ while conditioning on past and present is preferred when $D_{\text{chain}} < D_{\text{collider}}$.

Supplementary Figure 2 shows the values of $D_{\text{chain}}$ and $D_{\text{collider}}$ for different values of SEM parameters $\alpha$ and $\beta$. Each line corresponds to one value of $\beta$ (cross-correlation) and it is traversed from left to right as $\alpha$ is increased from 0 to 1. The area above the dashed black line indicates the range of parameters where conditioning only on the past is preferable ($D_{\text{chain}} > D_{\text{collider}}$). This is the case for all values of $\alpha$ if $\beta \lesssim 0.7$ (any autocorrelation strength in weakly- to moderate-connected networks) and for larger values of $\alpha$ if $\beta \gtrsim 0.7$ (strong autocorrelations in strongly-connected networks). This includes the majority of the joint values of $(\alpha, \beta)$, including brain networks for which regional autocorrelations are often notably stronger than correlations between regions (Nozari et al., 2023).

### 4 Computational Complexity of CaLLTiF

Based on the pseudocode shown in Algorithm 1 in the main text, the computational complexity of CaLLTiF is primarily determined by the total number and complexity of the conditional independence tests (line 7 of Algorithm 1). The total number of tests is $N^2(\tau_{\max} + 1)$, i.e., one for every pair of nodes and every time lag between 0 and $\tau_{\max}$. The complexity of each conditional independence test varies depending on its implementation. Here we used linear partial correlations given the linearity of resting state fMRI dynamics (Nozari et al., 2023), which significantly reduces the overall computational complexity of CaLLTiF. For partial correlations, a naive implementation using matrix inversion has a complexity of $O(p^3)$, where $p = N\tau_{\max}$ (contemporaneous edge) or $p = N\tau_{\max} - 1$ (lagged edge) is the size of the conditioning set. Nevertheless, following the original implementation of PCMCI (Runge et al., 2019), for the results shown in this work we used the `numpy.linalg.lstsq` function in Python, which uses an iterative optimization and can hence be significantly more efficient depending on its convergence. Therefore, the overall complexity of CaLLTiF is, in the worst case, $O(N^5\tau_{\max}^4)$. The heuristic steps at the end of the algorithm, such as performing OR operation across lags, statistical significance tests, or edge replacements do not significantly impact the overall complexity.

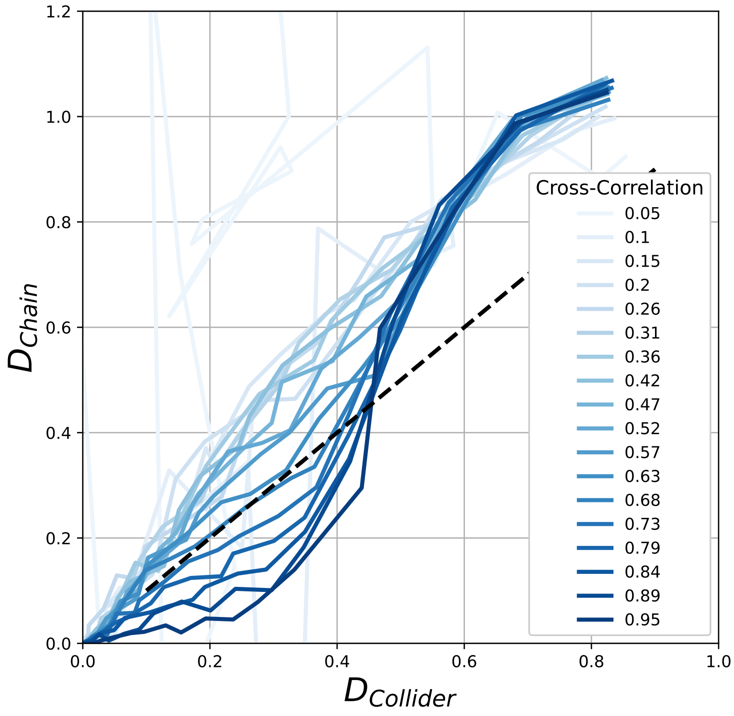

**Supplementary Figure 2: The desired and undesired effects of conditioning only on past variables, as opposed to conditioning on both past and contemporaneous variables in simple network motifs.** Each line shows the values of the metric $D$ defined in Eq. (S6) for two simple network motifs: the chain $(X_1(t) \rightarrow X_2(t) \rightarrow X_3(t))$ for which conditioning is generally helpful by avoiding/weakening a spurious edge between $X_1$ and $X_3$, and the collider $(X_1(t) \rightarrow X_2(t) \leftarrow X_3(t))$ for which conditioning is generally harmful by creating a spurious edge between the two. Simulated time series data is generated from each network using the SEMs in Eq. (S4) and Eq. (S5). Each line corresponds to one value of $\beta$ (cross-correlation) and it is traversed from left to right as $\alpha$ is increased from 0 to 1. The area above the dashed black line indicates the range of parameters where conditioning only on the past is preferable ($D_{\text{chain}} > D_{\text{collider}}$).

### Supplementary Figures for Simulated fMRI from Simple Networks

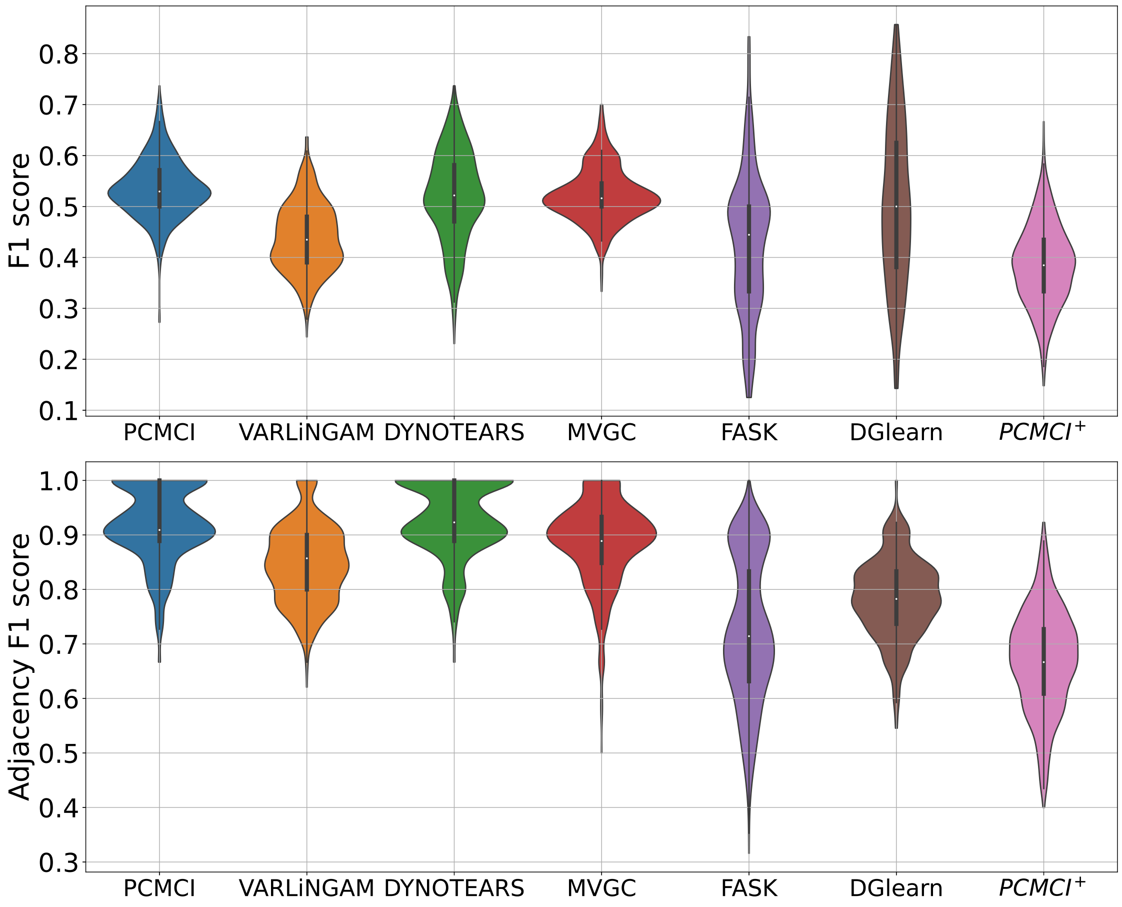

**Supplementary Figure 3: Results of comparing several state-of-the-art causal discovery algorithms over a benchmark of simulated fMRI generated from simple networks with 5-10 nodes each.** F1 score of seven state-of-the-art algorithms (six from Table 1 and MVGC) for correctly identifying the full (directed) graphs and corresponding underlying undirected graphs, respectively. All methods are evaluated using optimized values of their respective hyperparameters (see Methods). The benchmark data includes 60 repetitions of fMRI data from each of the 9 graphs, so each violin plot is based on 540 samples.

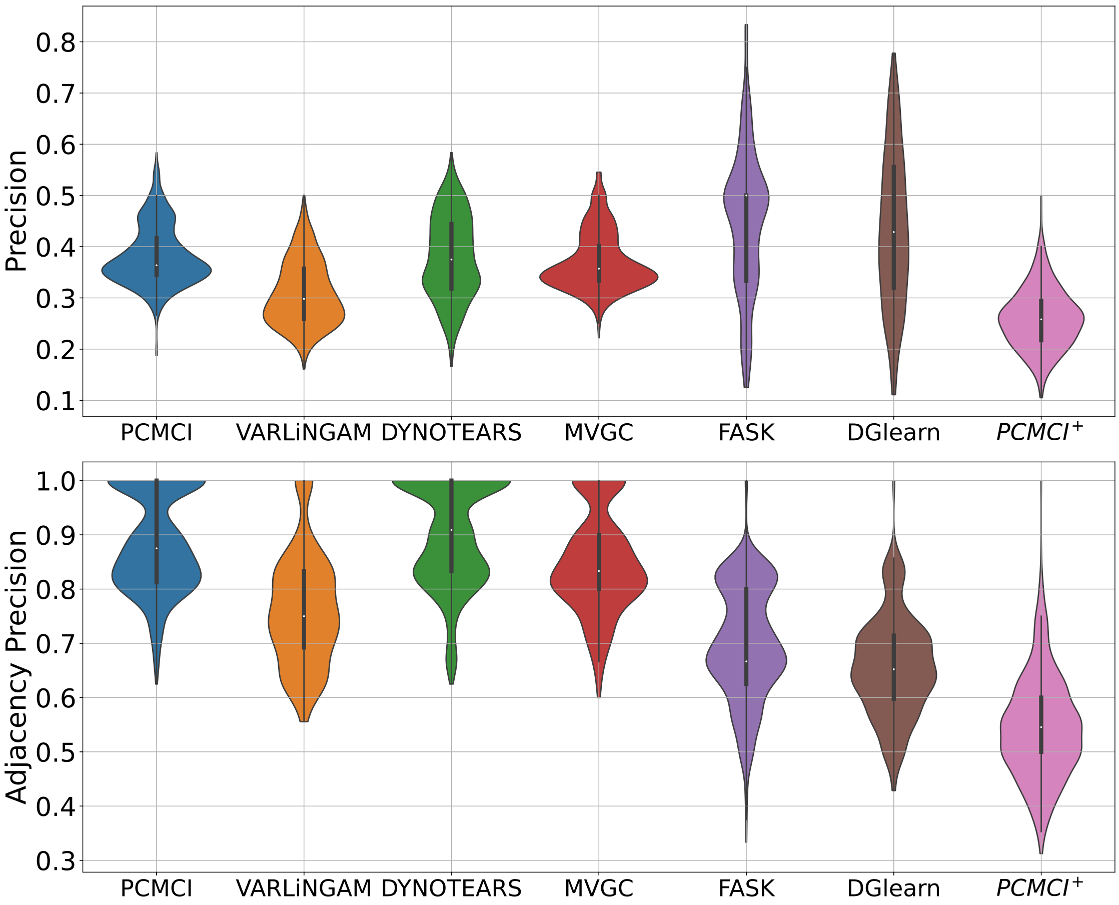

**Supplementary Figure 4: Results of comparing several state-of-the-art causal discovery algorithms over a benchmark of simulated fMRI generated from simple networks with 5-10 nodes each.** Precision of seven state-of-the-art algorithms (six from Table 1 and MVGC) for correctly identifying the full (directed) graphs and corresponding underlying undirected graphs, respectively. All methods are evaluated using optimized values of their respective hyperparameters (see Methods). The benchmark data includes 60 repetitions of fMRI data from each of the 9 graphs, so each violin plot is based on 540 samples.

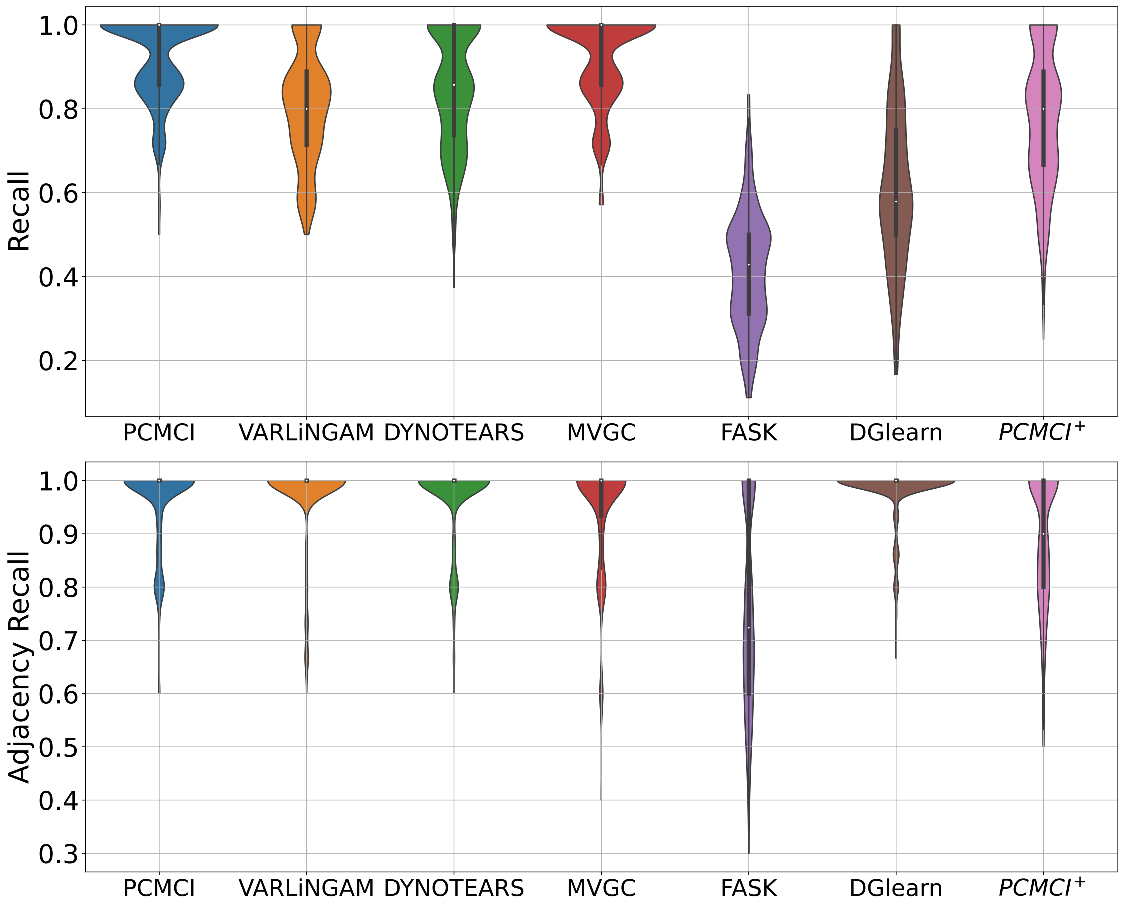

**Supplementary Figure 5: Results of comparing several state-of-the-art causal discovery algorithms over a benchmark of simulated fMRI generated from simple networks with 5-10 nodes each.** Recall of seven state-of-the-art algorithms (six from Table 1 and MVGC) for correctly identifying the full (directed) graphs and corresponding underlying undirected graphs, respectively. All methods are evaluated using optimized values of their respective hyperparameters (see Methods). The benchmark data includes 60 repetitions of fMRI data from each of the 9 graphs, so each violin plot is based on 540 samples.

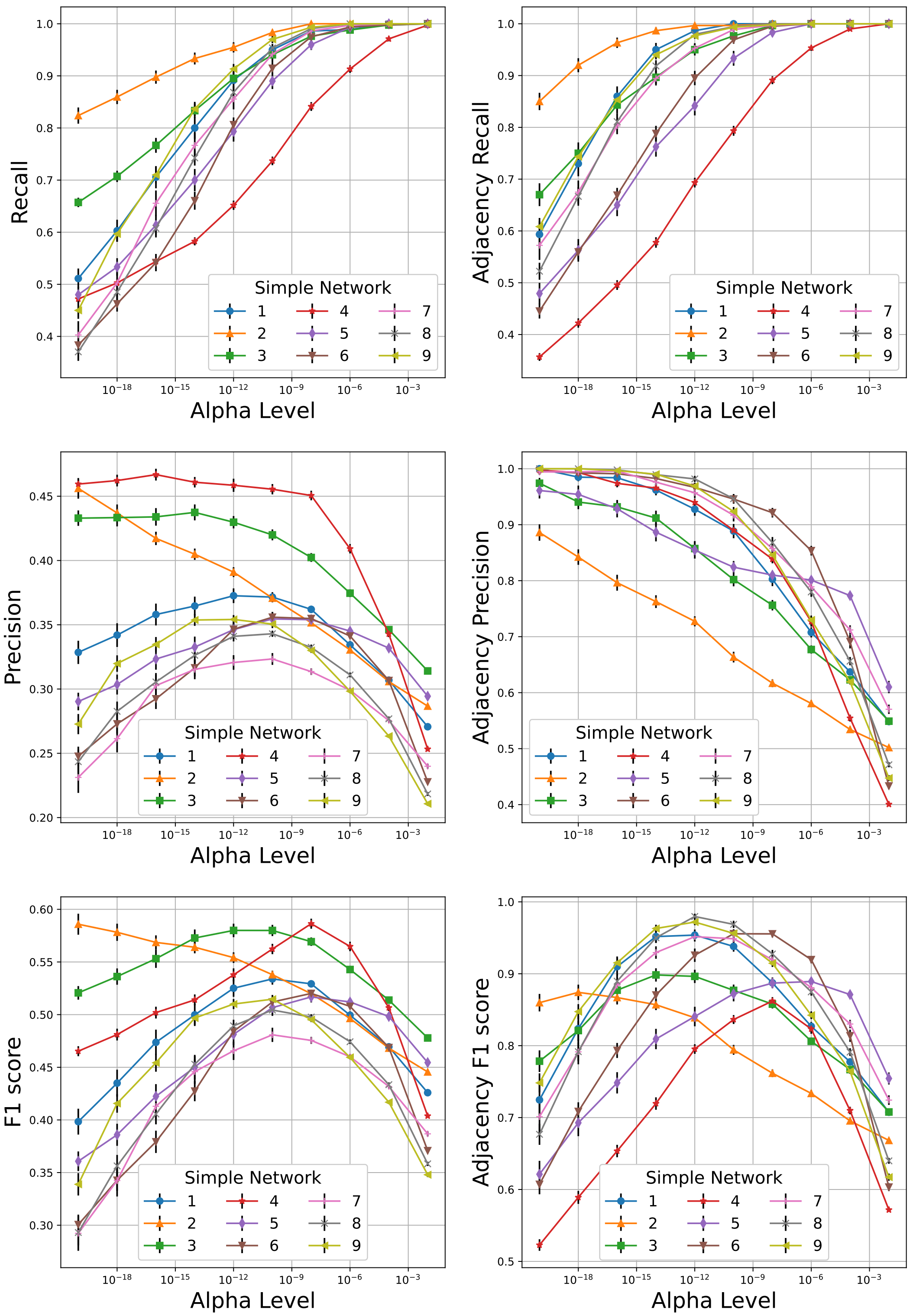

**Supplementary Figure 6: Detailed performance curves of PCMCI over simulated fMRI from simple networks for varying values of its hyperparameter Alpha Level.** In all graphs, the error bars depict the standard error of the mean.

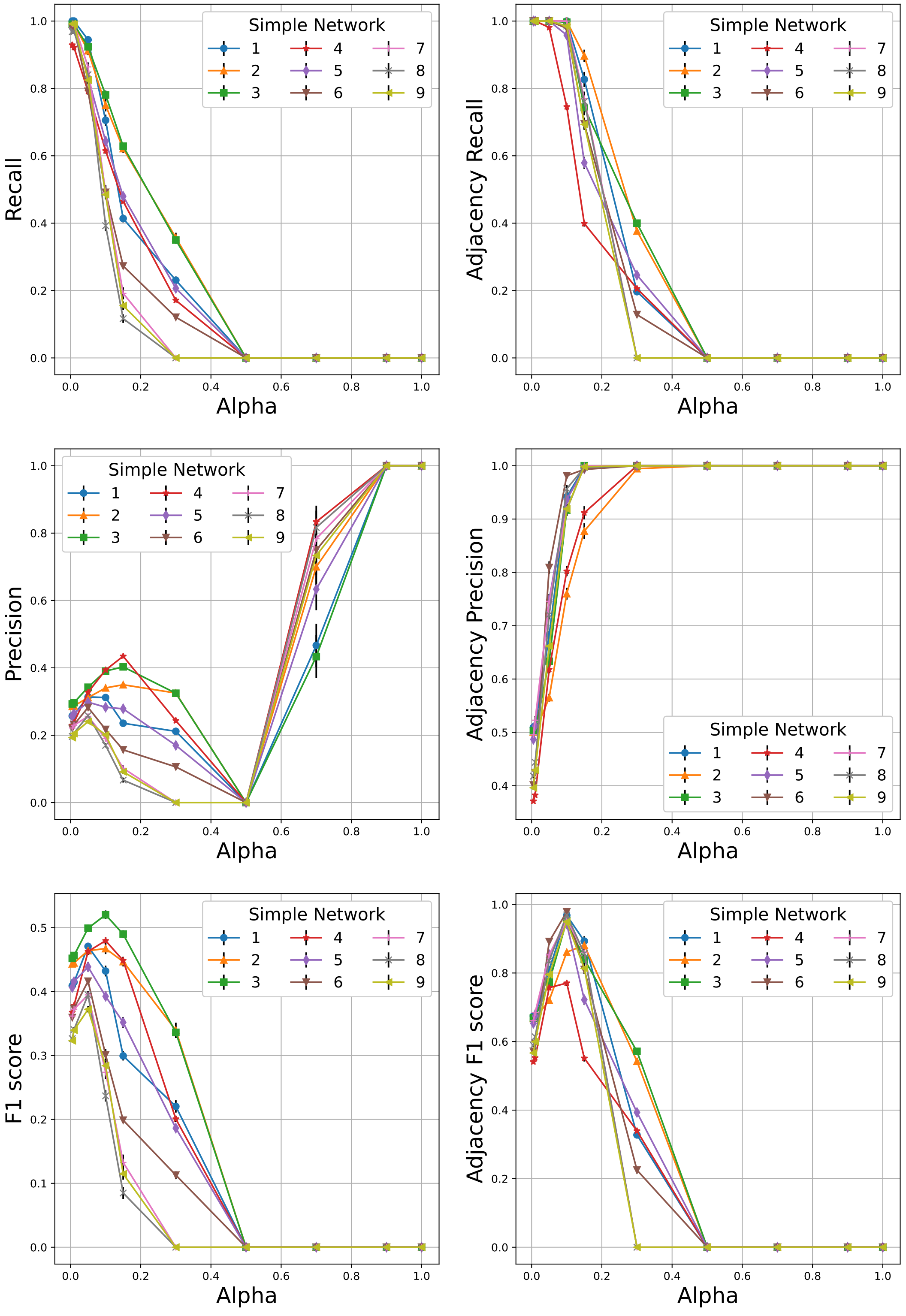

**Supplementary Figure 7: Detailed performance curves of VARLiNGAM over simulated fMRI from simple networks for varying values of its hyperparameter Alpha.** In all graphs, the error bars depict the standard error of the mean.

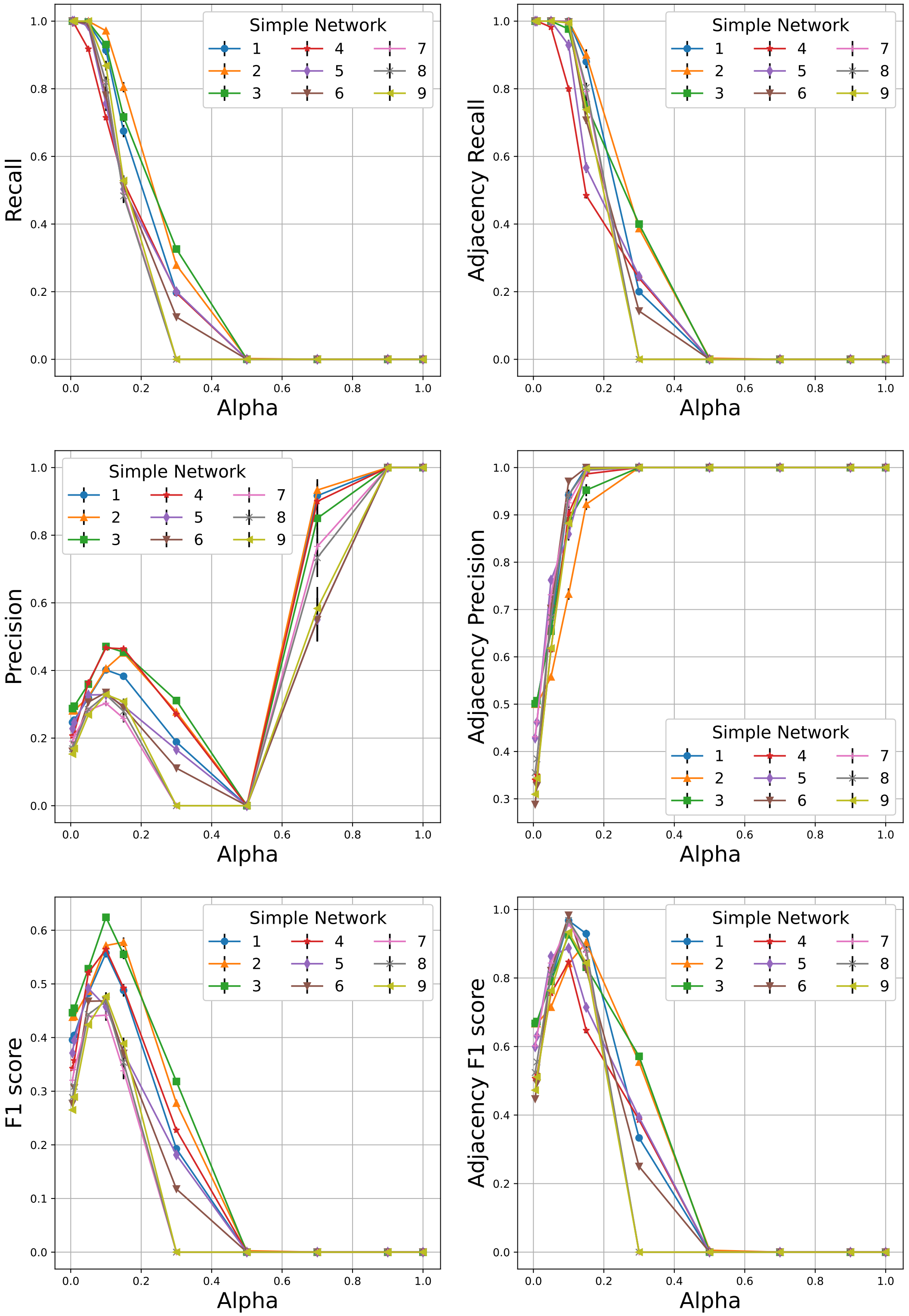

**Supplementary Figure 8: Detailed performance curves of DYNOTEARS over simulated fMRI from simple networks for varying values of its hyperparameter Alpha.** In all graphs, the error bars depict the standard error of the mean.

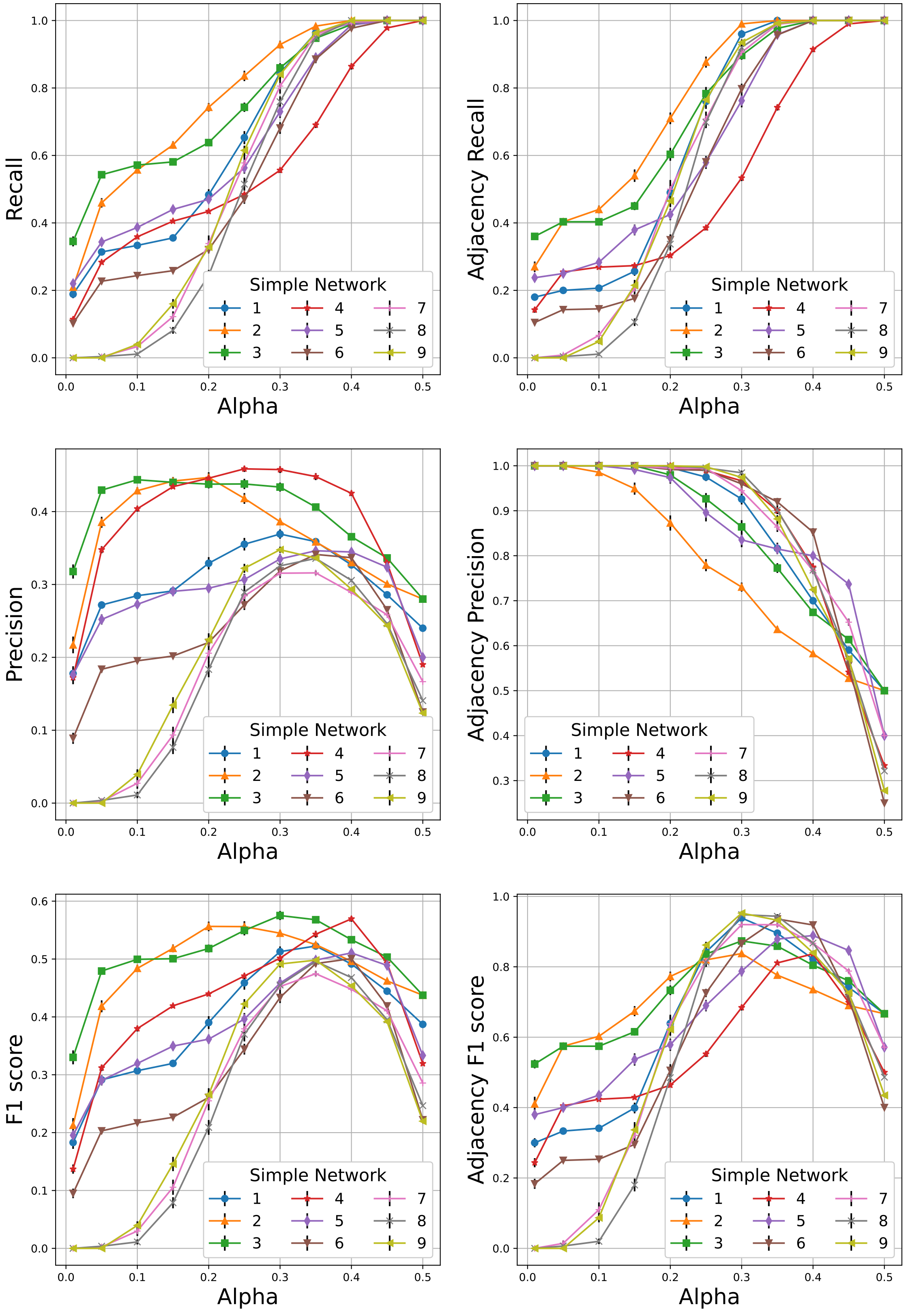

**Supplementary Figure 9: Detailed performance curves of MVGC over simulated fMRI from simple networks for varying values of its hyperparameter Alpha.** In all graphs, the error bars depict the standard error of the mean.

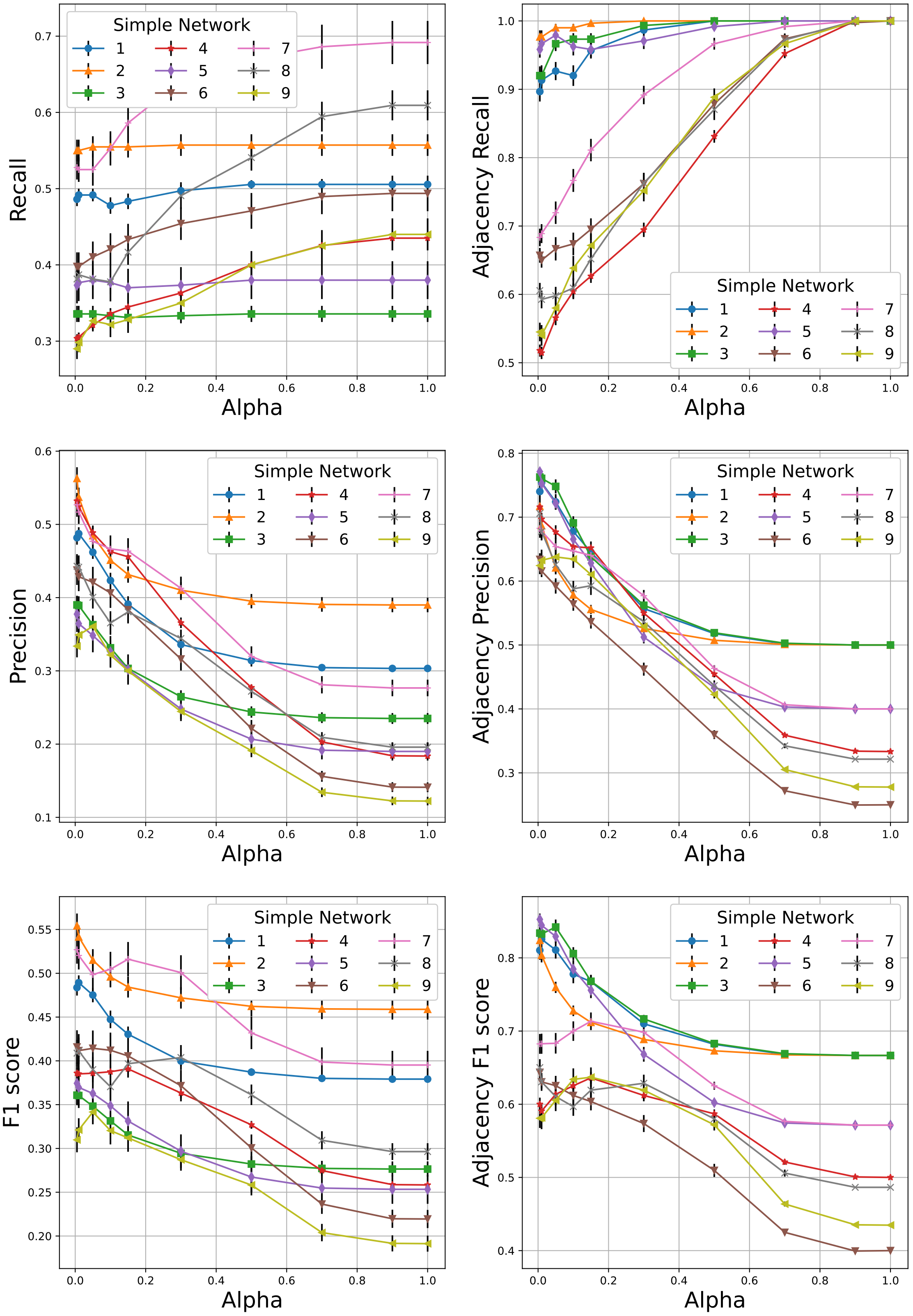

**Supplementary Figure 10: Detailed performance curves of FASK over simulated fMRI from simple networks for varying values of its hyperparameter Alpha.** In all graphs, the error bars depict the standard error of the mean.

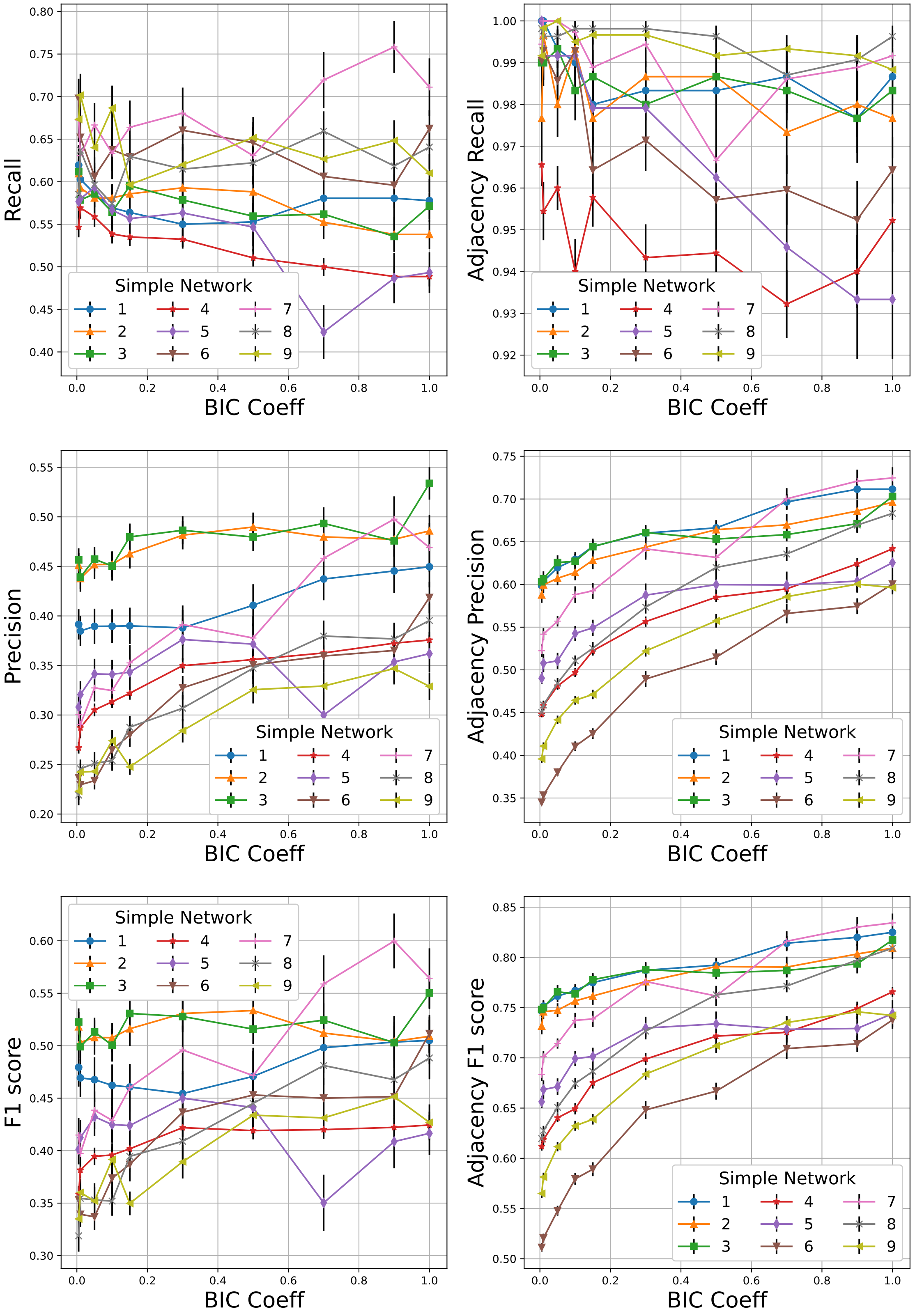

**Supplementary Figure 11: Detailed performance curves of DGlearn over simulated fMRI from simple networks for varying values of its hyperparameter BIC coefficient.** In all graphs, the error bars depict the standard error of the mean.

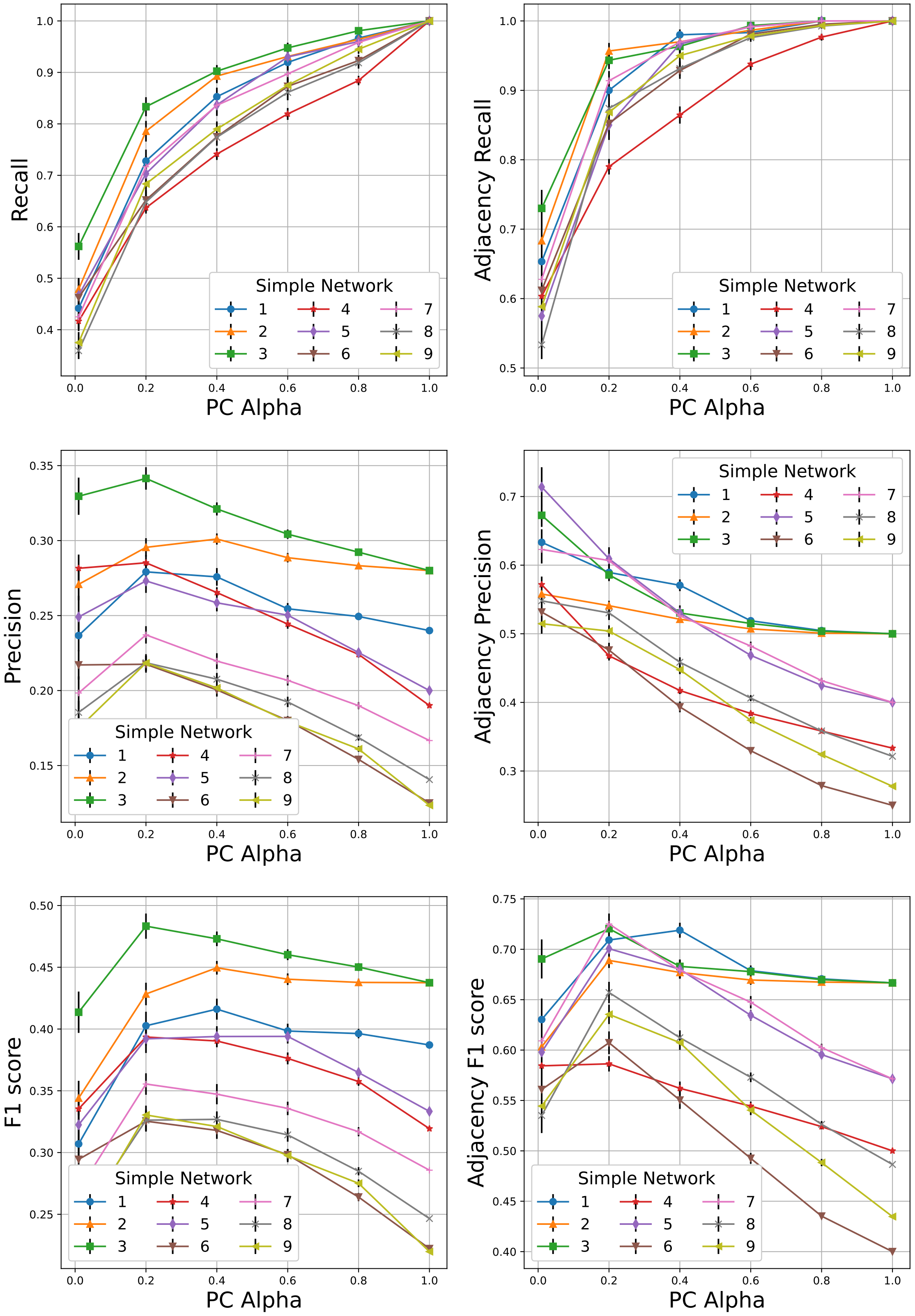

**Supplementary Figure 12: Detailed performance curves of PCMCI$^+$ over simulated fMRI from simple networks for varying values of its hyperparameter PC Alpha.** In all graphs, the error bars depict the standard error of the mean.

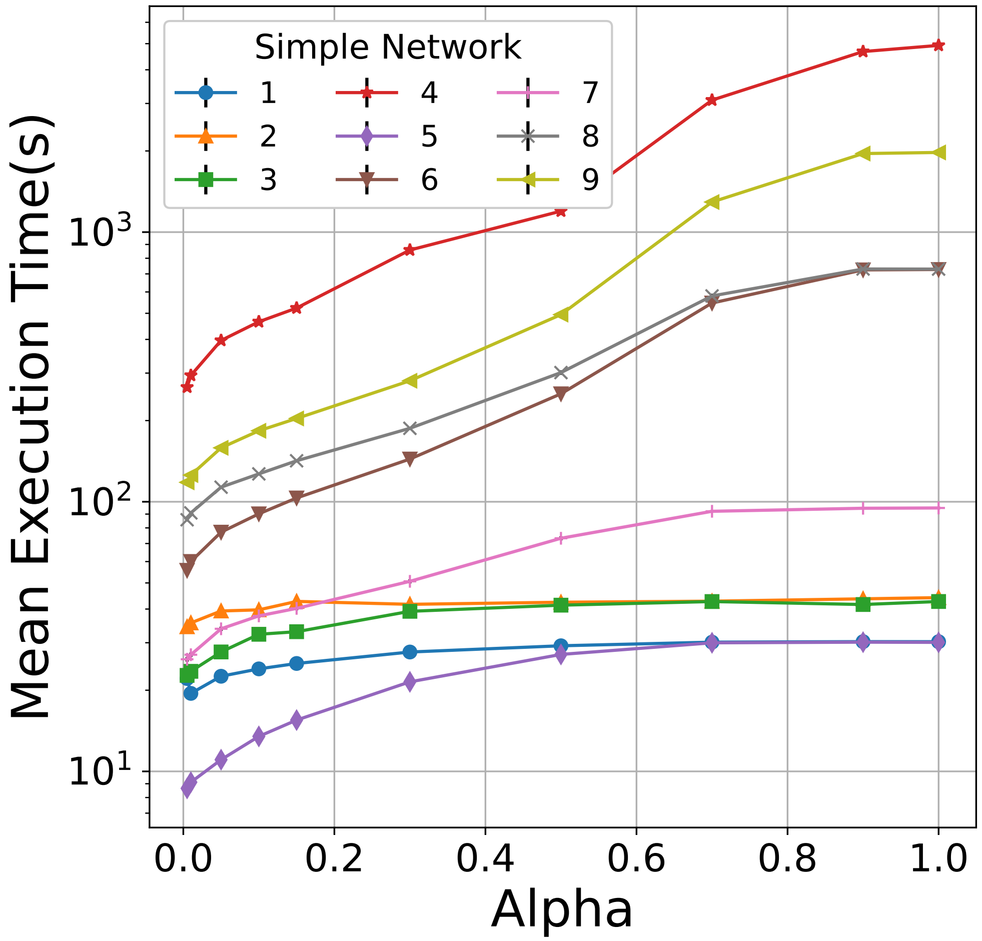

**Supplementary Figure 13: Execution run times of FASK over simulated fMRI from simple networks for varying values of its hyperparameter Alpha.** Error bars depict the standard error of the mean.

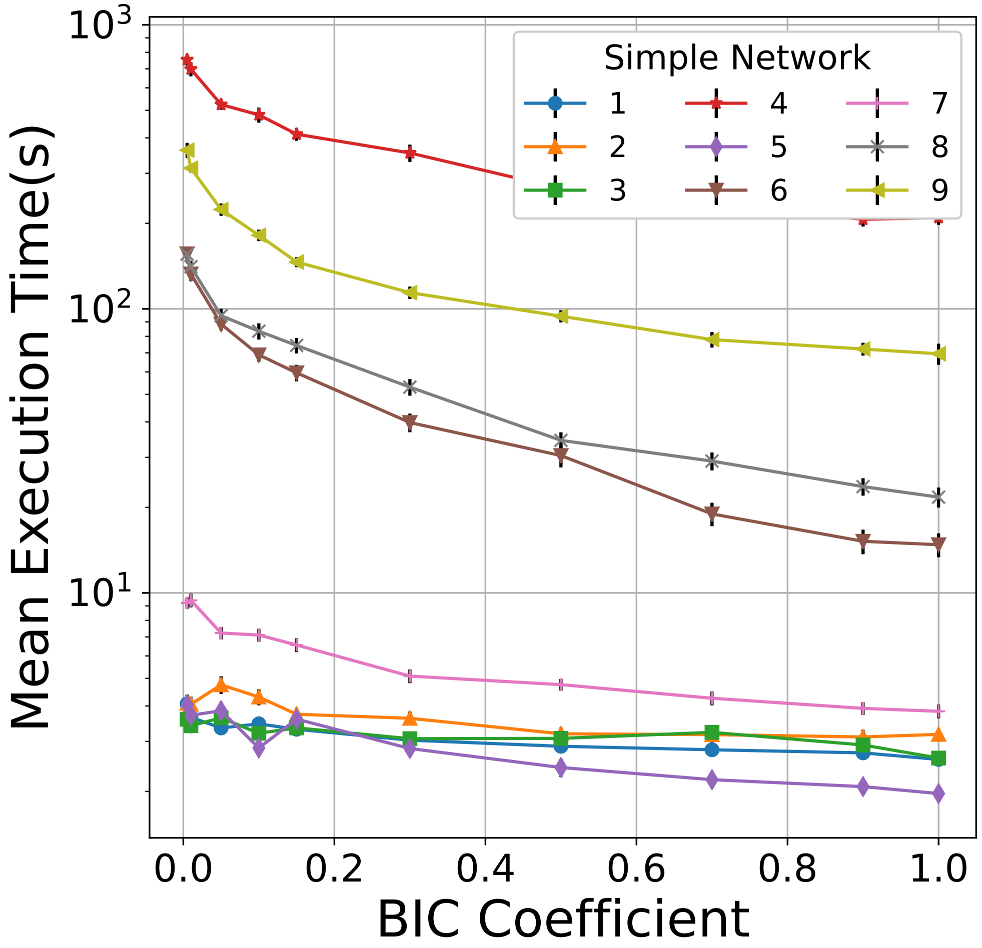

**Supplementary Figure 14: Execution run times of DGlearn over simulated fMRI from simple networks for varying values of its hyperparameter BIC coefficient.** In all graphs, the error bars depict the standard error of the mean.

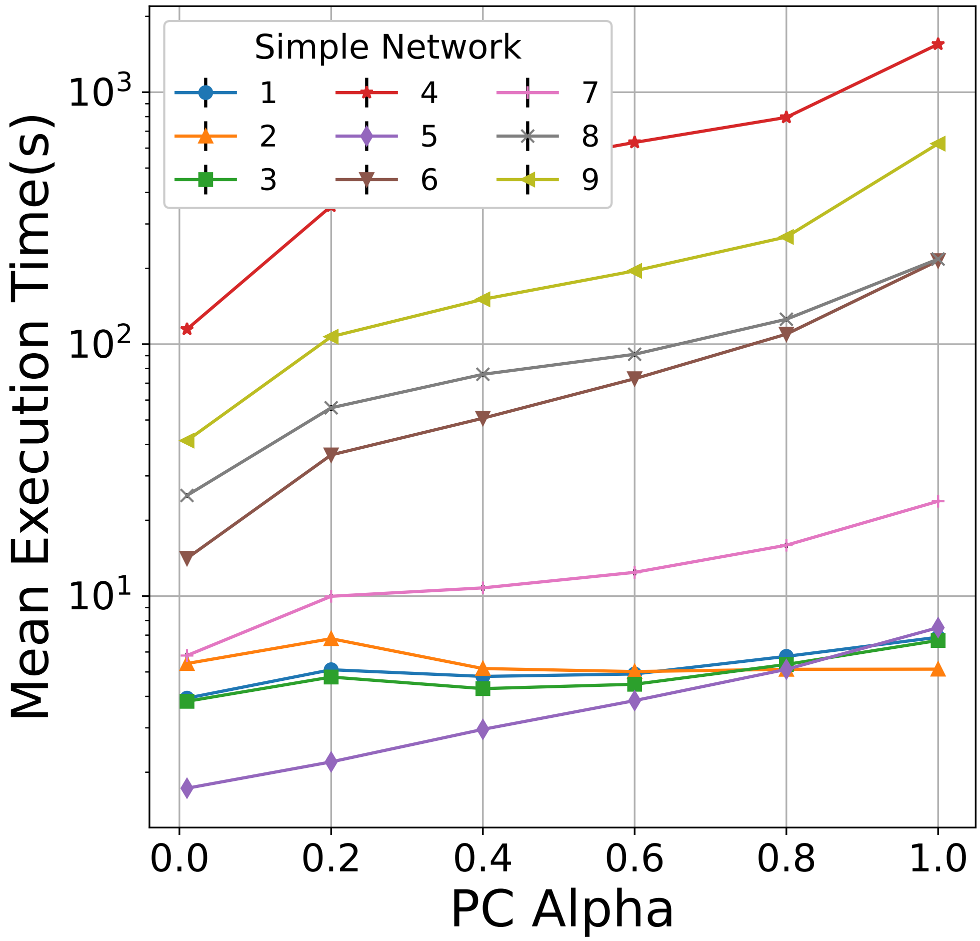

**Supplementary Figure 15: Execution run times of PCMCI$^+$ over simulated fMRI from simple networks for varying values of its hyperparameter PC Alpha.** Error bars depict the standard error of the mean.

### Supplementary Figures for Simulated fMRI from Macaque_SmallDegree Network

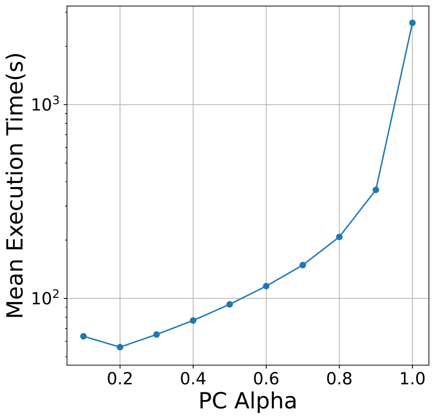

**(a)**

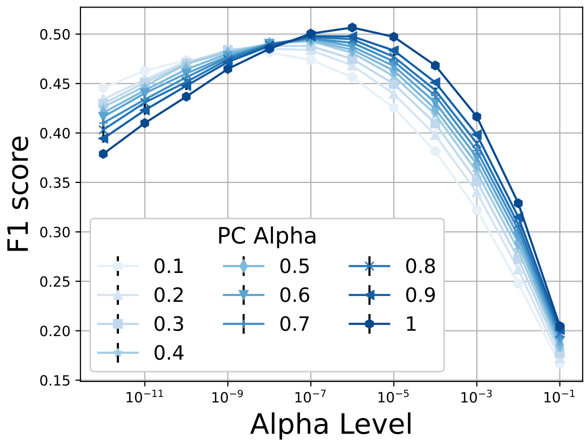

**(b)**

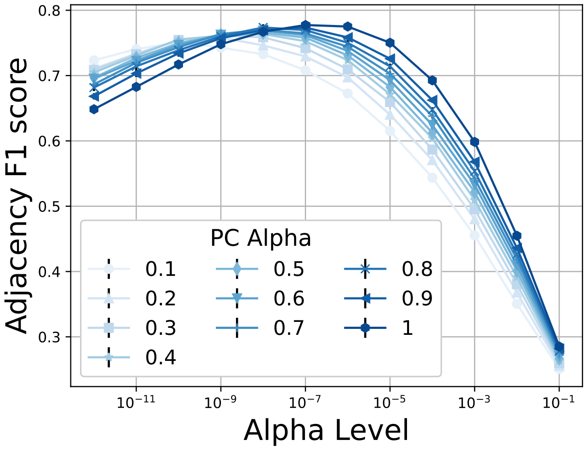

**(c)**

**Supplementary Figure 16: The effect of the 'PC Alpha' hyperparameter of PCMCI on its performance over simulated Macaque_SmallDegree fMRI.** (a) Mean Execution time for different values of PC Alpha, showing a super-exponential growth. (b,c) F1 scores for varying values of PC Alpha and Alpha Level. Alpha Level denotes a second hyperparameter that controls the sparsity of the returned graph (the higher the Alpha Level the denser the resulting graphs). Best F1 scores are achieved for PC Alpha = 1. In all graphs, the error bars depict the standard error of the mean.

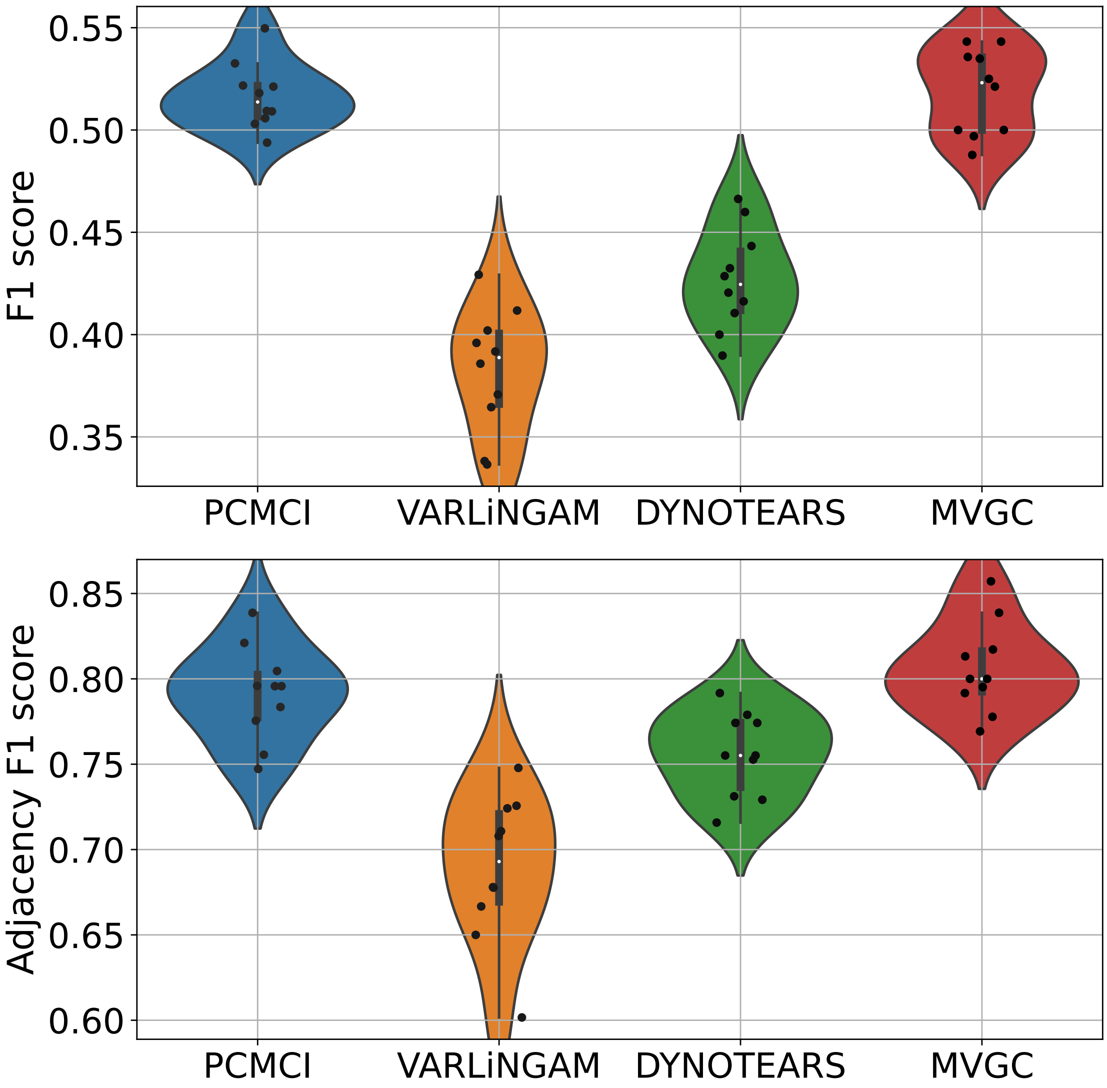

**Supplementary Figure 17: Comparing the scalable subset of algorithms from Figure 1 over simulated fMRI data from the Macaque_SmallDegree benchmark .** F1 score of identifying the full (directed) graph (top). Each distribution is calculated based on 10 repetitions of simulated data from the same underlying graph. Corresponding Adjacency F1 score of identifying the underlying undirected graph of the Macaque_SmallDegree network (bottom).

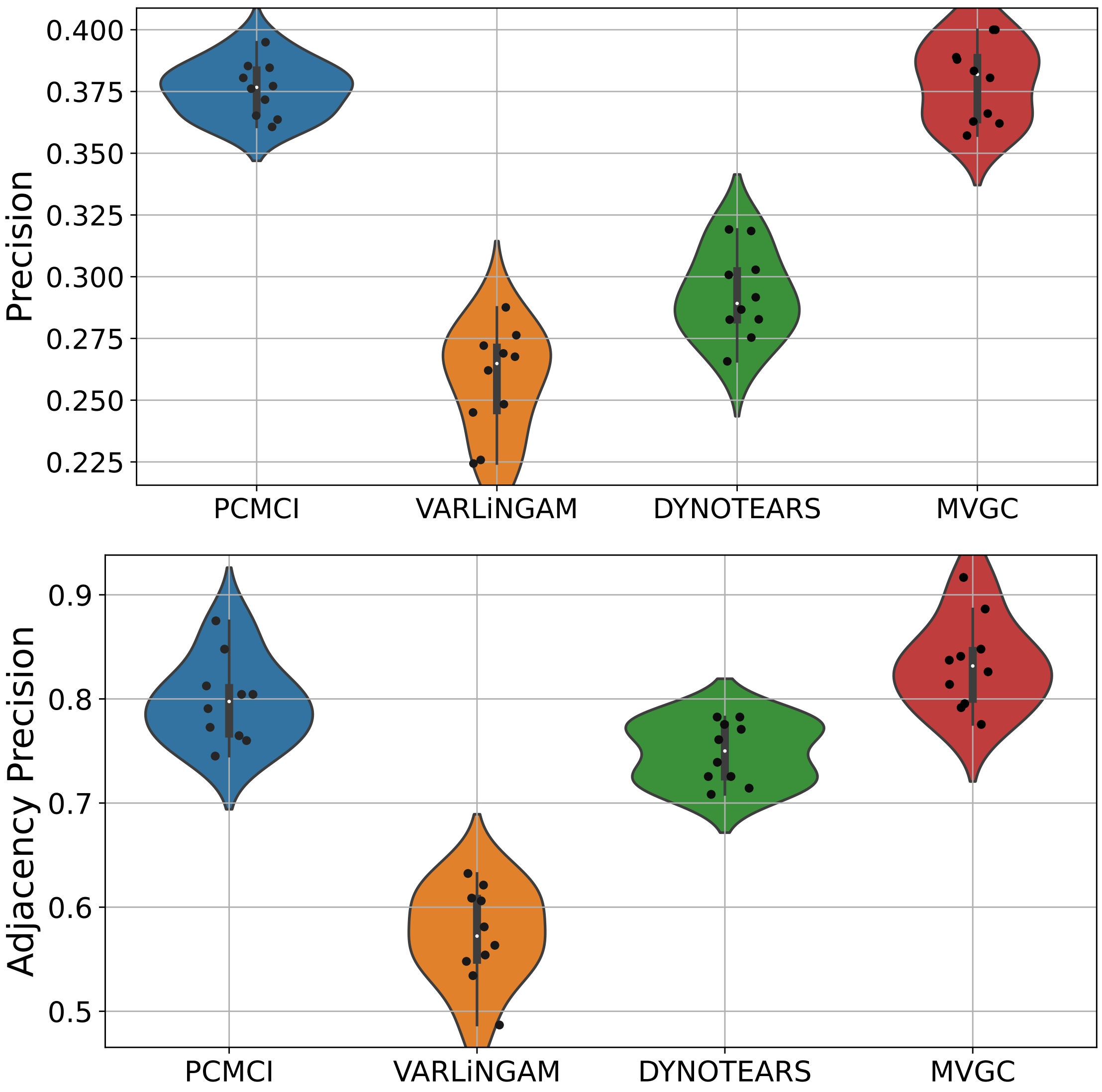

**Supplementary Figure 18: Comparing the scalable subset of algorithms from Figure 1 over simulated fMRI data from the Macaque_SmallDegree benchmark .** Precision of identifying the full (directed) graph (top). Each distribution is calculated based on 10 repetitions of simulated data from the same underlying graph. Corresponding Adjacency precision of identifying the underlying undirected graph of the Macaque_SmallDegree network (bottom).

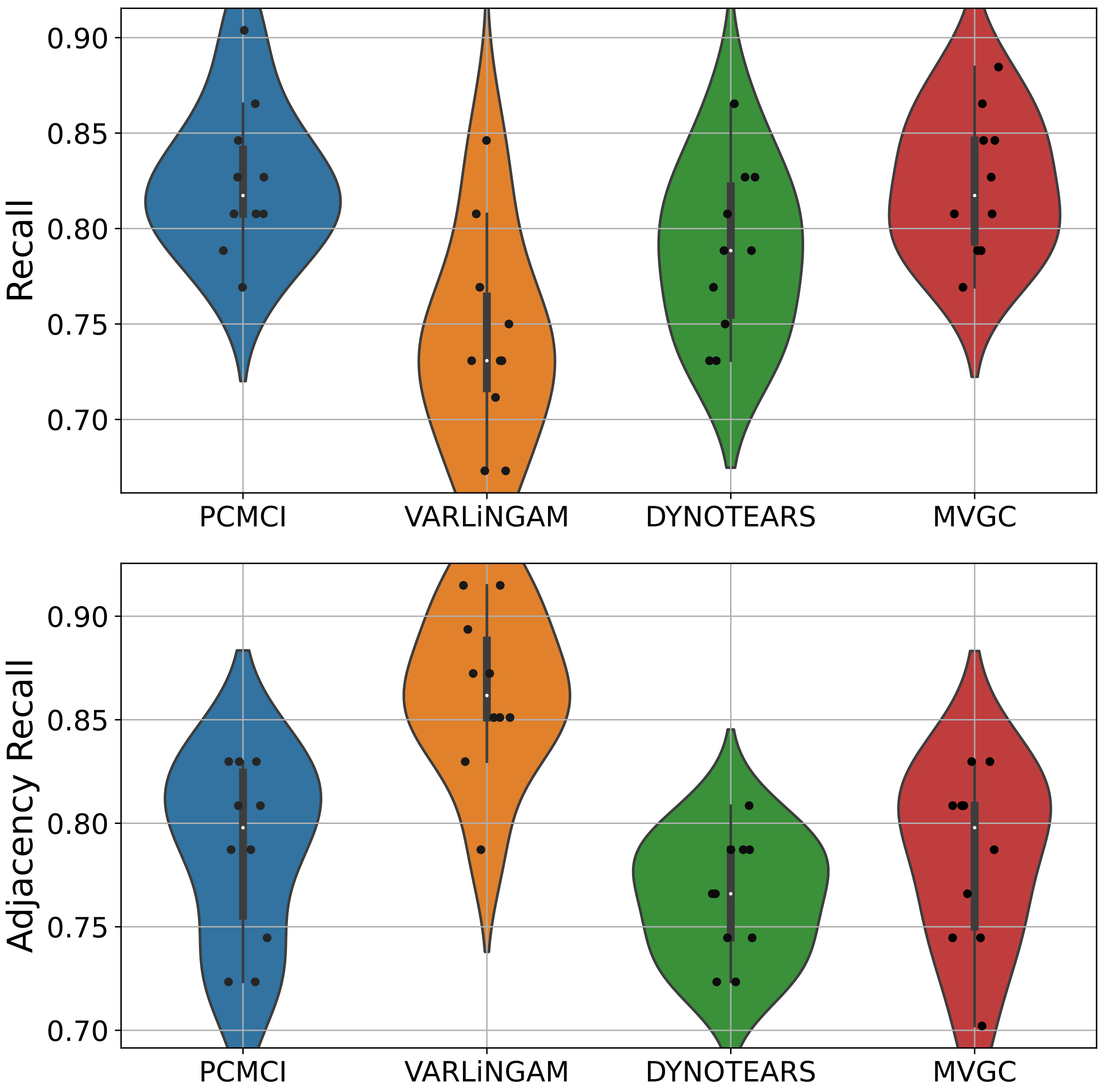

**Supplementary Figure 19: Comparing the scalable subset of algorithms from Figure 1 over simulated fMRI data from the Macaque_SmallDegree benchmark .** Recall of identifying the full (directed) graph (top). Each distribution is calculated based on 10 repetitions of simulated data from the same underlying graph. Corresponding Adjacency recall of identifying the underlying undirected graph of the Macaque_SmallDegree network (bottom).

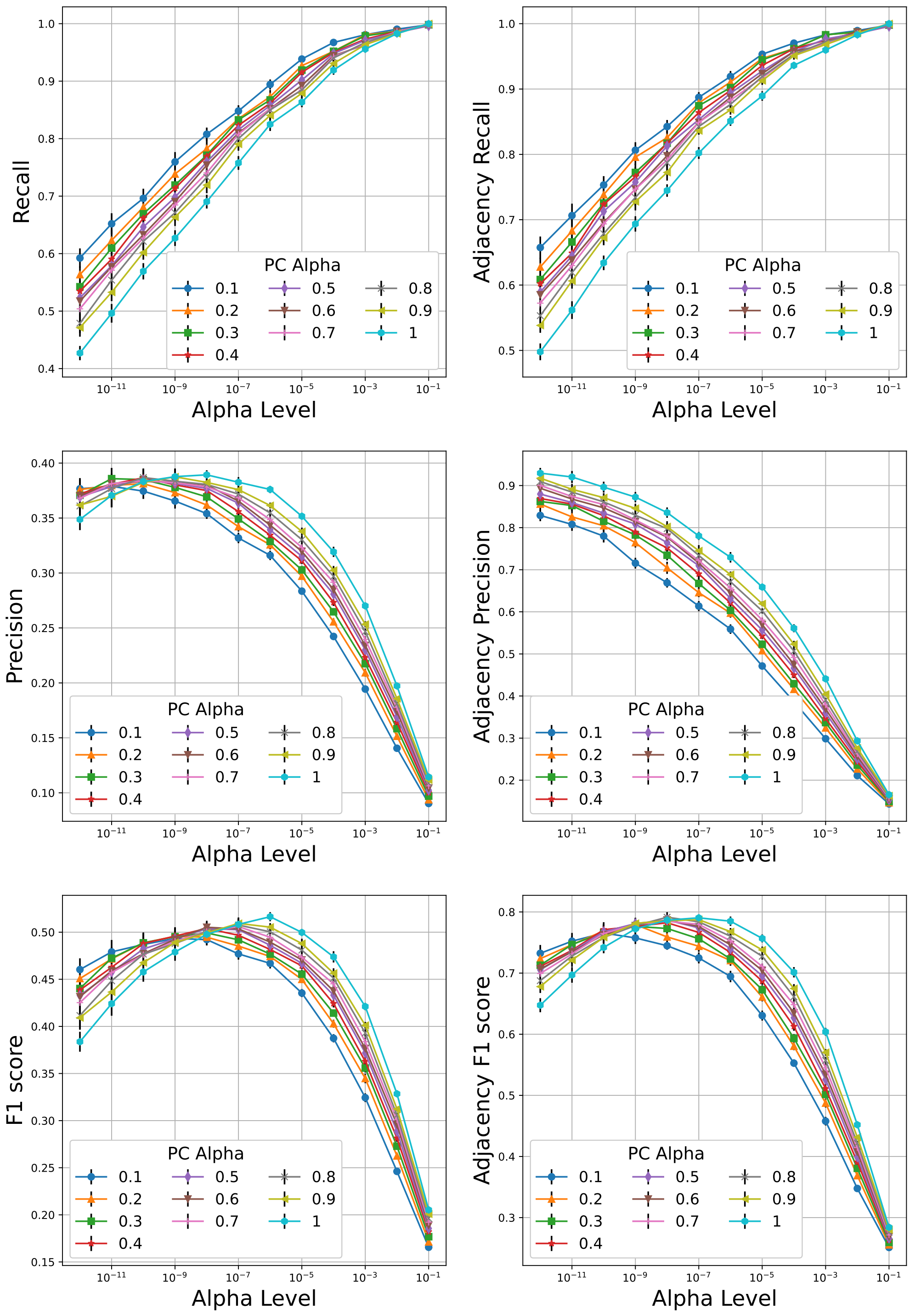

**Supplementary Figure 20: Detailed performance curves of PCMCI over simulated fMRI from Macaque_SmallDegree network for varying values of its hyperparameters PC Alpha and Alpha Level.** In all graphs, the error bars depict the standard error of the mean.

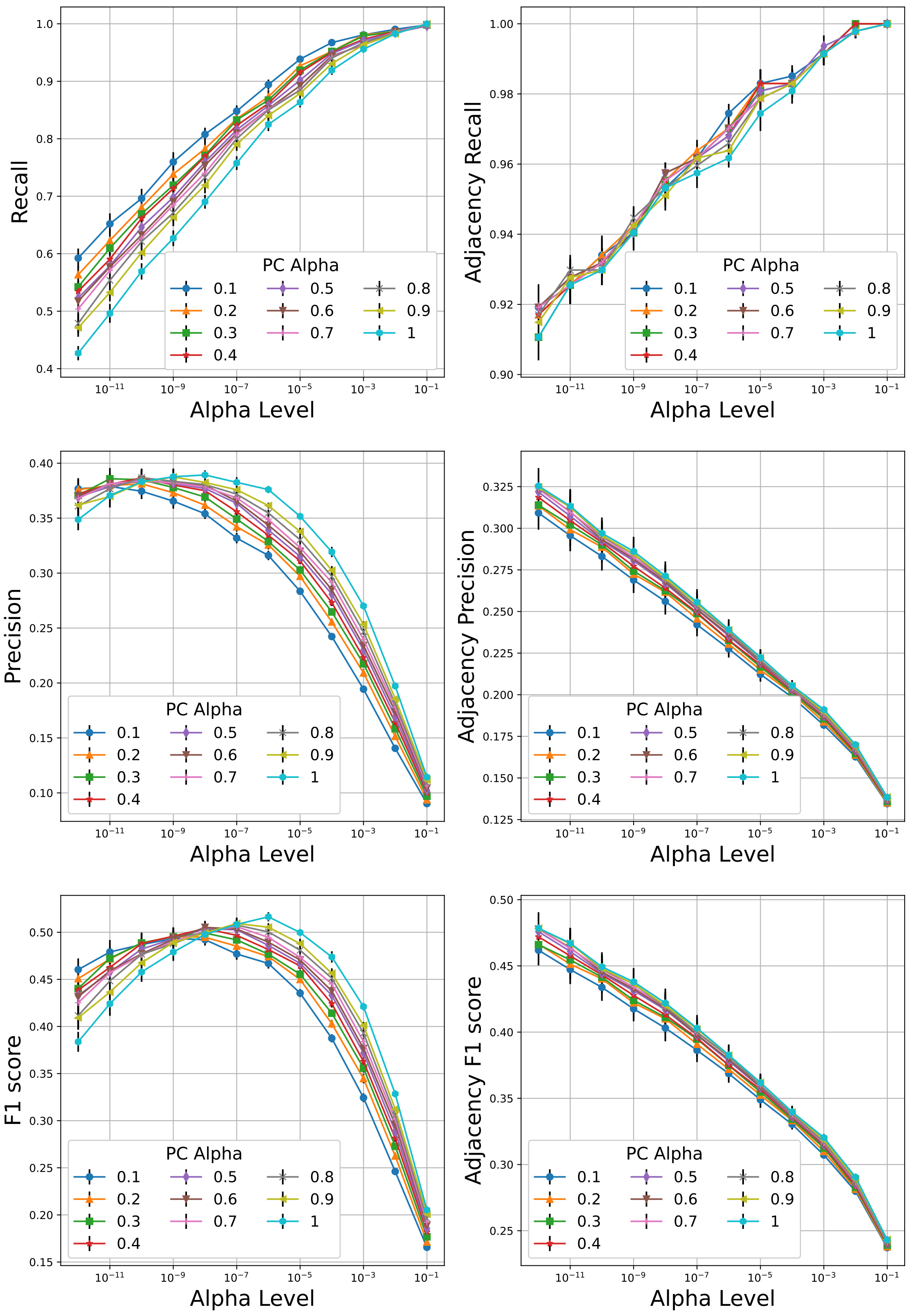

**Supplementary Figure 21: Detailed performance curves of Mixed-PCMCI over simulated fMRI from Macaque_SmallDegree network for varying values of its hyperparameters PC Alpha and Alpha Level.** In all graphs, the error bars depict the standard error of the mean.

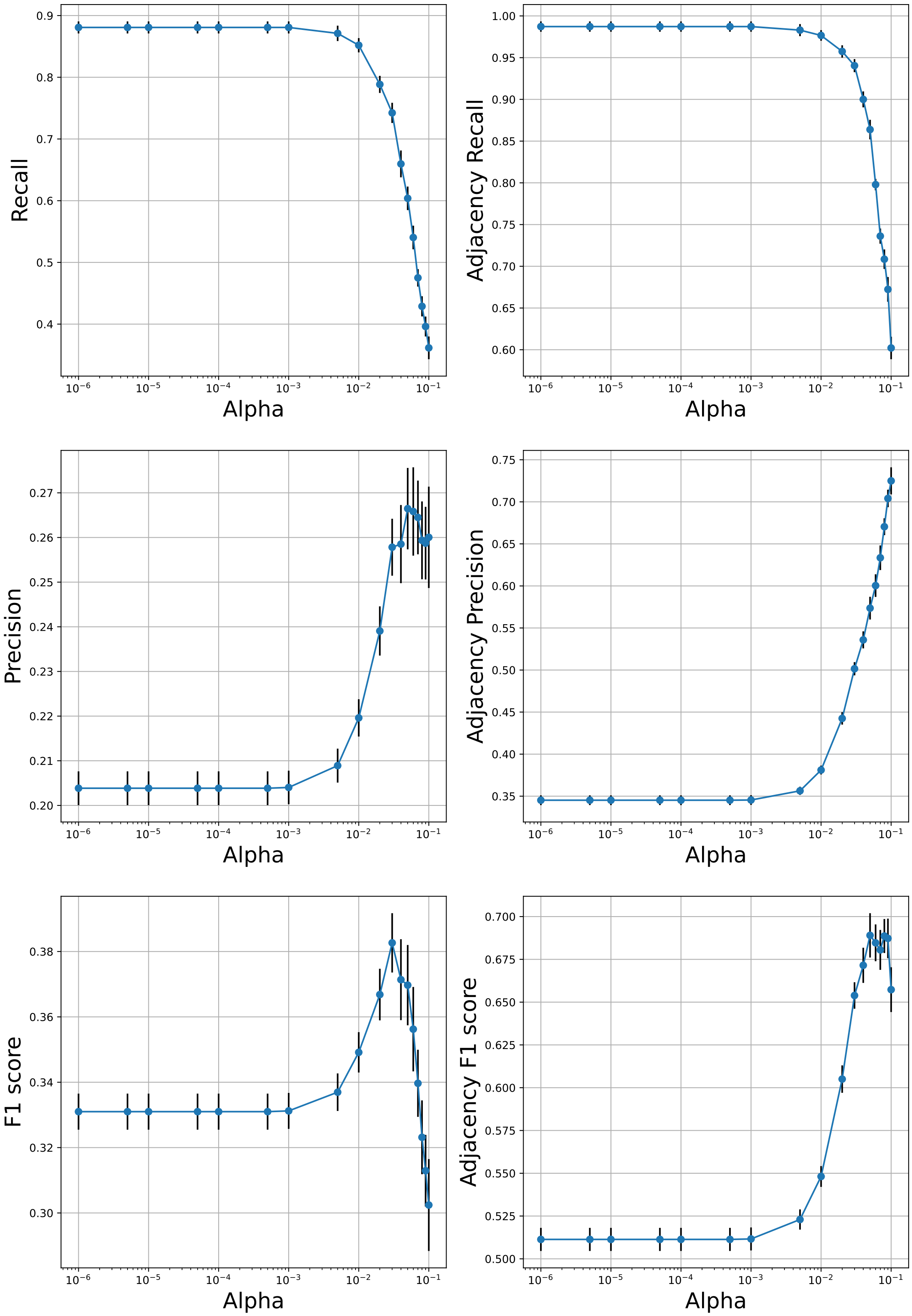

**Supplementary Figure 22: Detailed performance curves of VARLiNGAM over simulated fMRI from Macaque_SmallDegree network for varying values of its hyperparameter Alpha.** In all graphs, the error bars depict the standard error of the mean.

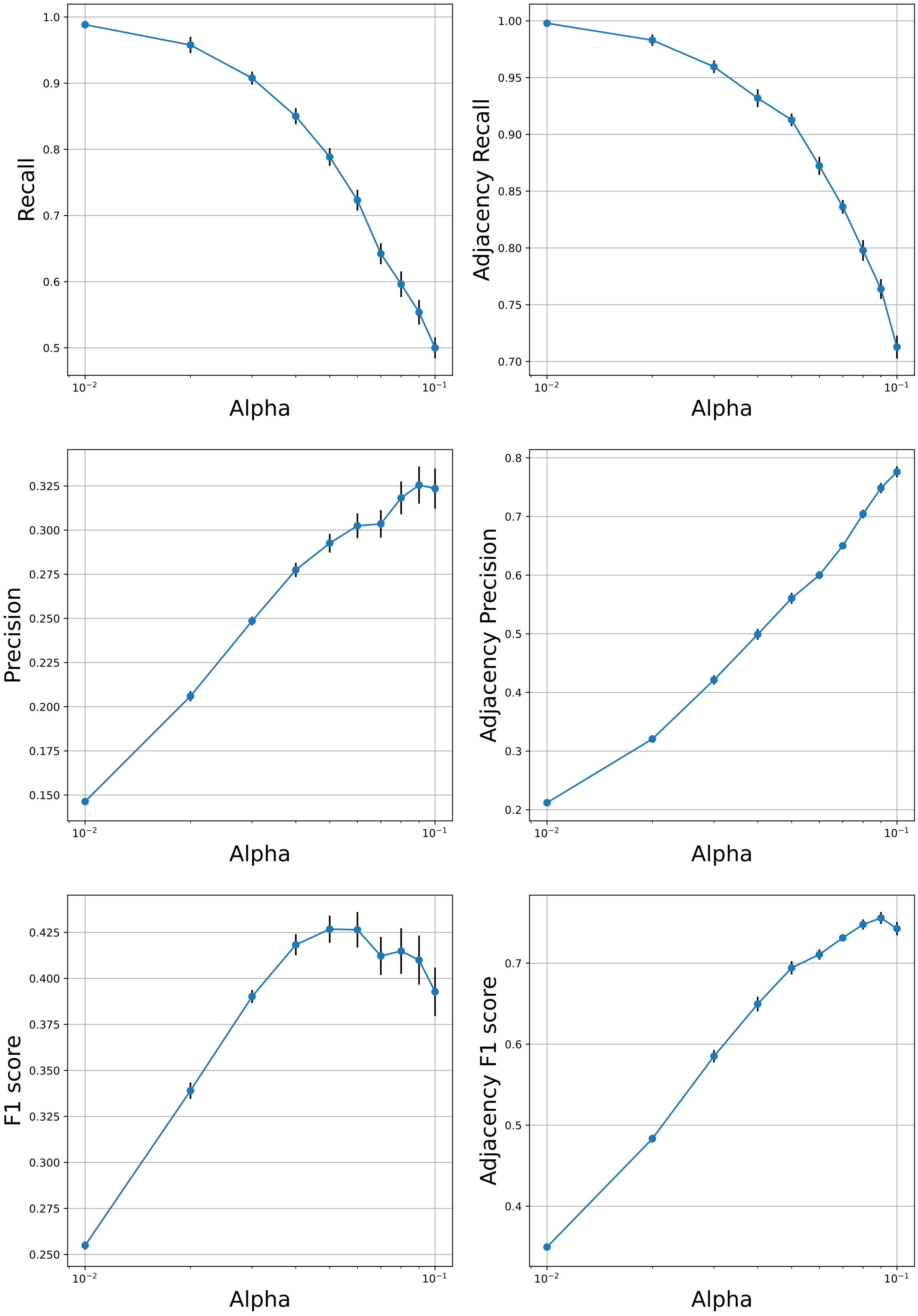

**Supplementary Figure 23: Detailed performance curves of DYNOTEARS over simulated fMRI from Macaque_SmallDegree network for varying values of its hyperparameter Alpha.** In all graphs, the error bars depict the standard error of the mean.

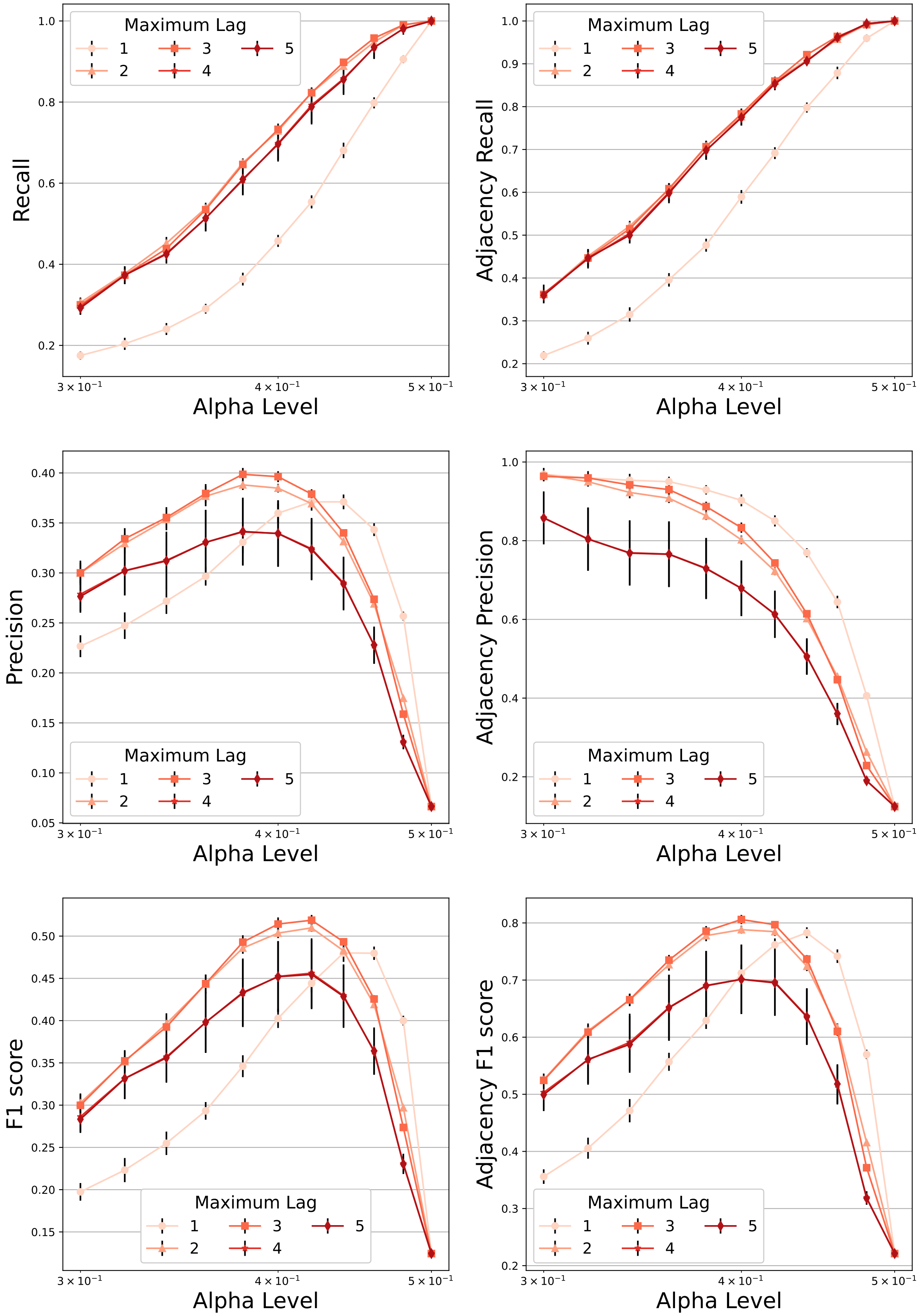

**Supplementary Figure 24: Detailed performance curves of MVGC over simulated fMRI from Macaque_SmallDegree network for varying values of its hyperparameters Alpha and maximum number of lags.** In all graphs, the error bars depict the standard error of the mean.

#### Supplementary Figures for Simulated fMRI from Macaque_Full Network

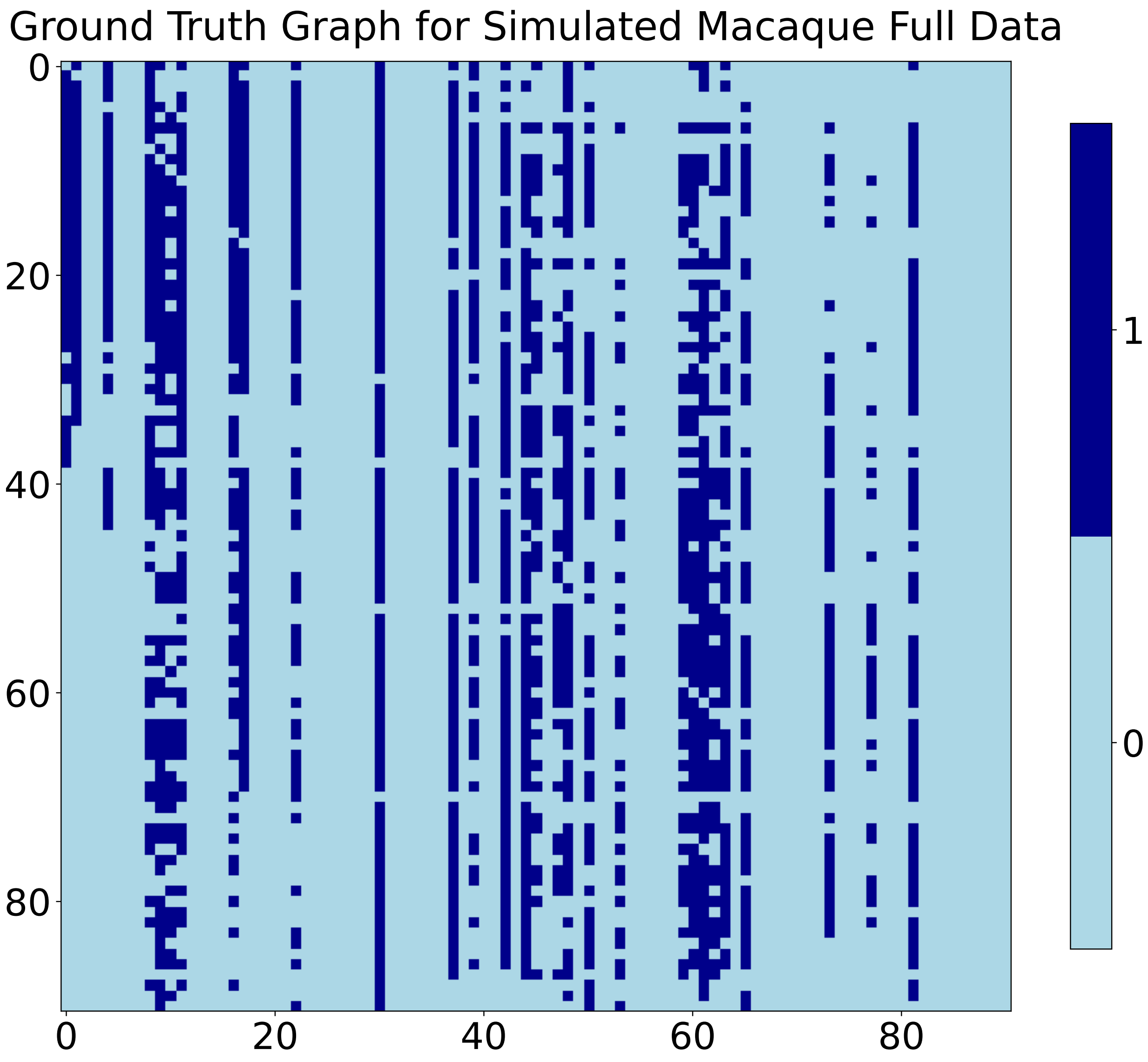

**Supplementary Figure 25: Heat map displaying the ground truth directed matrix for the Macaque_Full simulated dataset.**

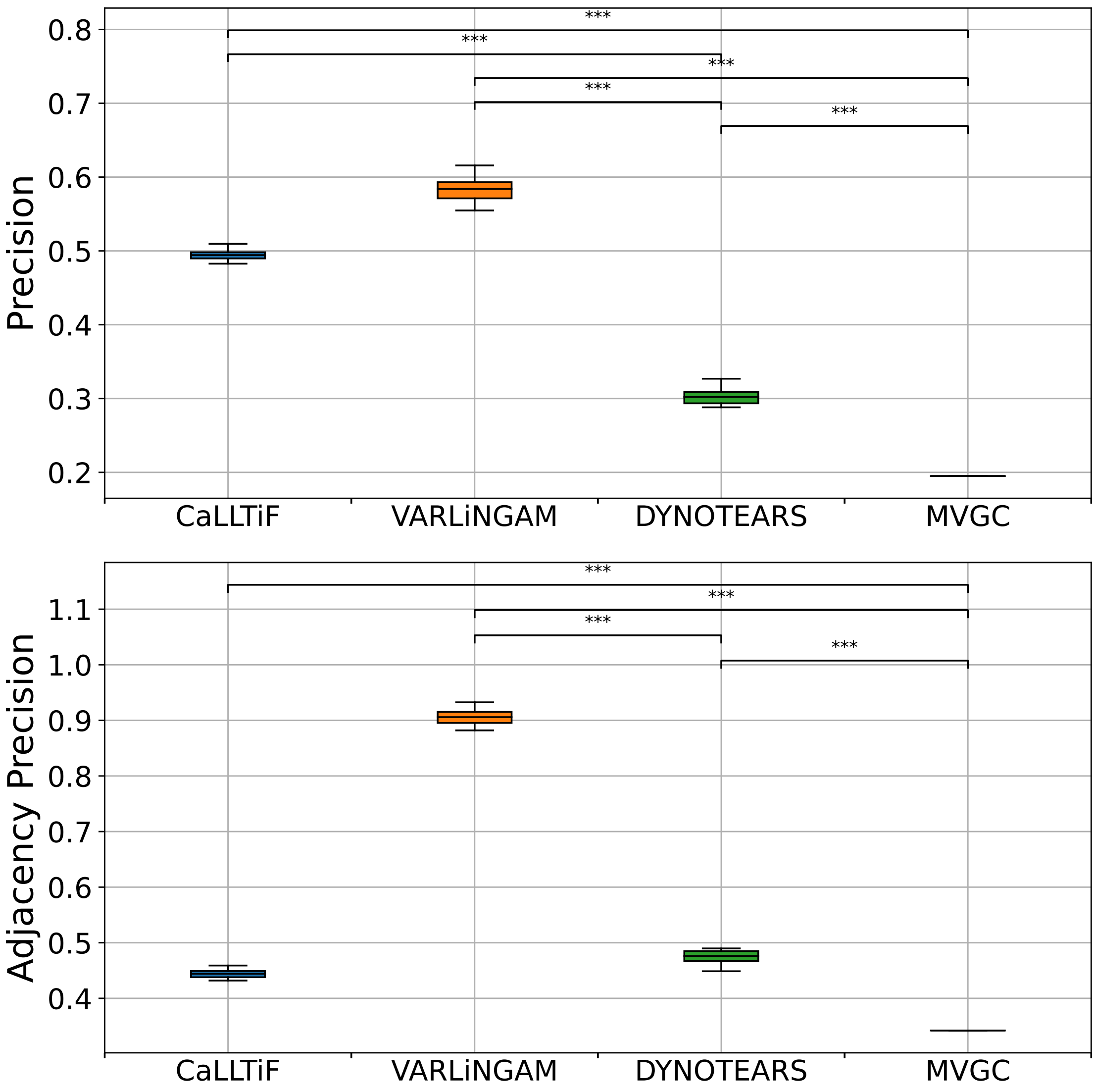

**Supplementary Figure 26: Comparisons between the proposed algorithm (CaLLTiF) and state-of-the-art alternatives over simulated fMRI from the Macaque_Full connectome.** Distributions of Precision and Recall for CaLLTiF and state-of-the-art alternatives. Precision is shown for the values of hyperparameters which give the maximum F1 score. For all repetitions, the best performance of MVGC occurs at $\alpha = 0.5$ which returns a complete graph, hence the point distributions for MVGC. *** denotes $p < 0.001$. All statistical comparisons are performed using a one-sided Wilcoxon signed-rank test. In all boxplots, the center line represents the median, the box spans the interquartile range (IQR), and the whiskers extend up to 1.5 times the IQR from the box limits.

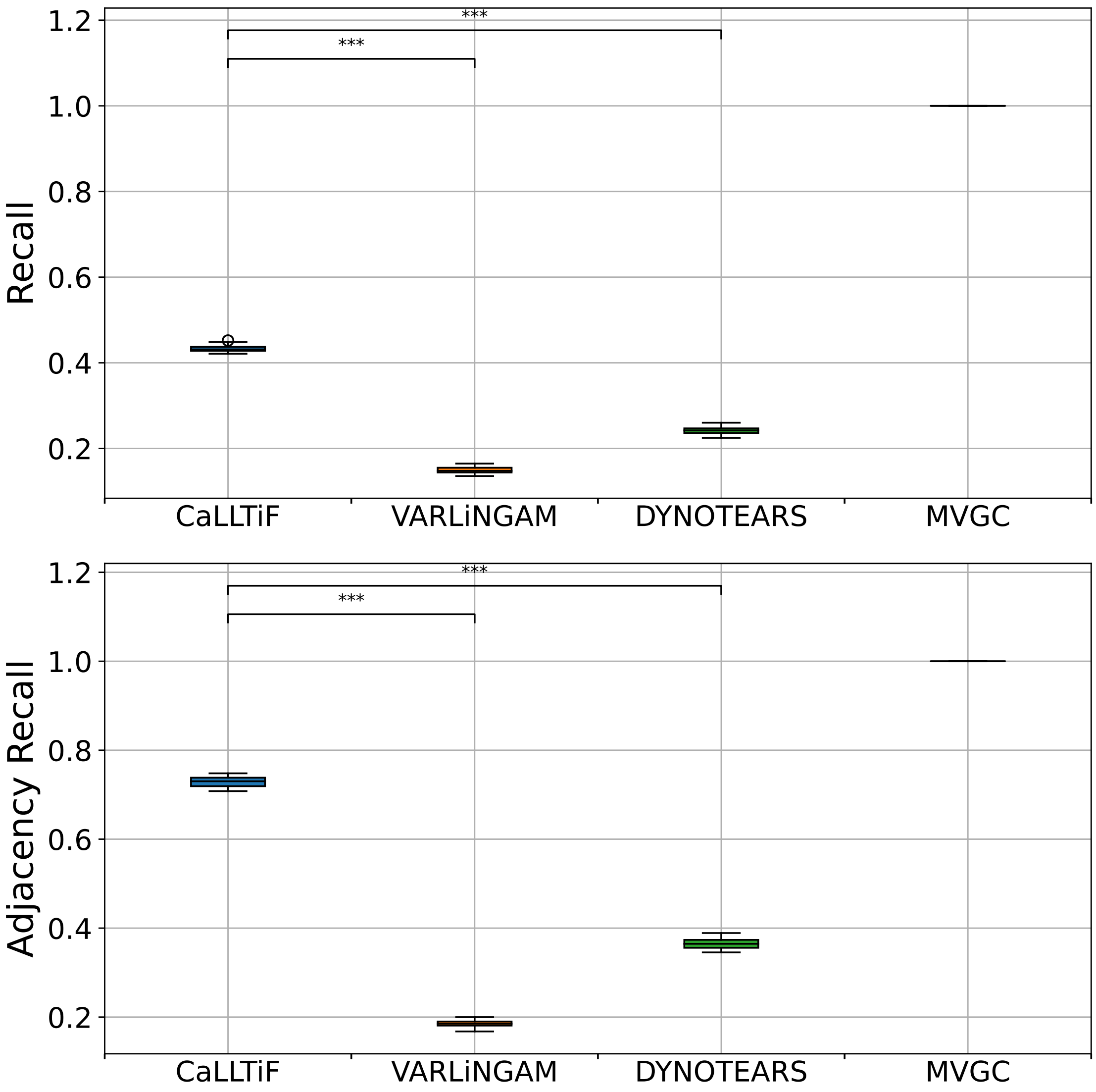

**Supplementary Figure 27: Comparisons between the proposed algorithm (CaLLTiF) and state-of-the-art alternatives over simulated fMRI from the Macaque_Full connectome.** Distributions of Precision and Recall for CaLLTiF and state-of-the-art alternatives. Recall is shown for the values of hyperparameters which give the maximum F1 score. For all repetitions, the best performance of MVGC occurs at $\alpha = 0.5$ which returns a complete graph, hence the point distributions for MVGC. $^{***}$ denotes $p < 0.001$. All statistical comparisons are performed using a one-sided Wilcoxon signed-rank test. In all boxplots, the center line represents the median, the box spans the interquartile range (IQR), and the whiskers extend up to 1.5 times the IQR from the box limits.

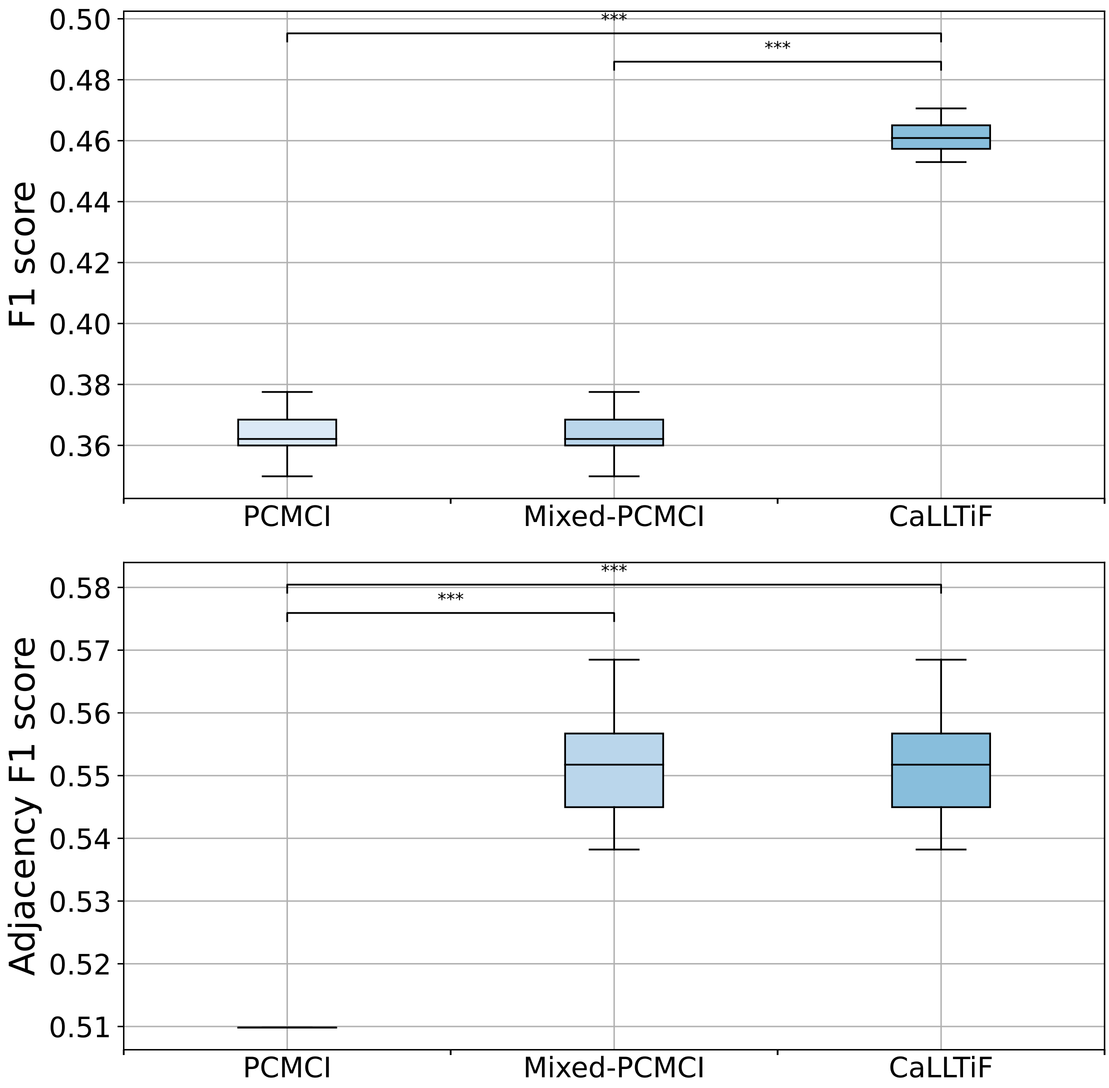

**Supplementary Figure 28: Comparisons between the proposed algorithm (CaLLTiF) and state-of-the-art alternatives over simulated fMRI from the Macaque_Full connectome.** Distributions of F1 score for PCMCI (ignoring the contemporaneous ⊶ connections), Mixed-PCMCI (using the contemporaneous ⊶ connections only for adjacency), and CaLLTiF. F1 scores are shown for the value of hyperparameters which give the maximum F1 score.

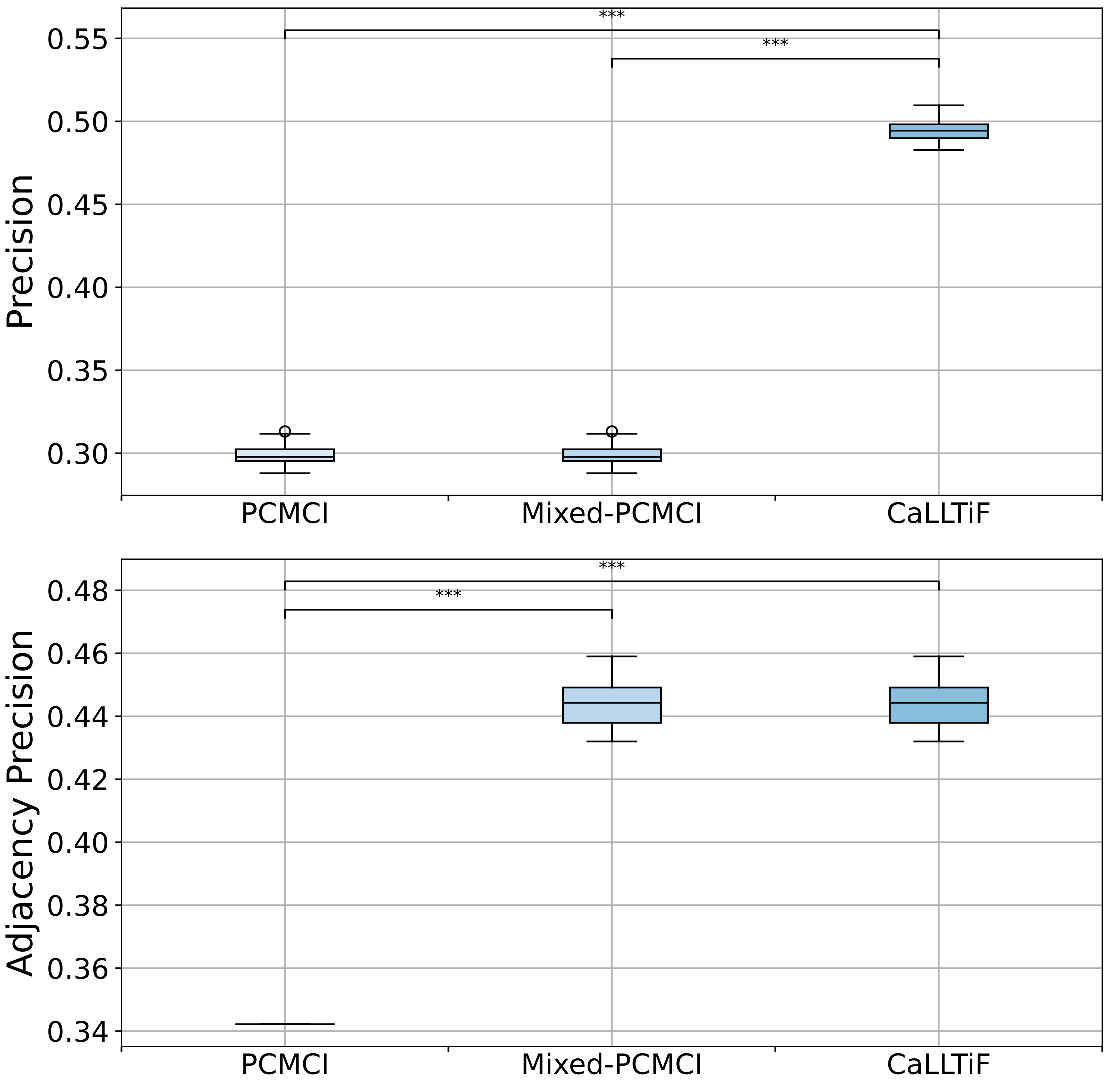

**Supplementary Figure 29: Comparisons between the proposed algorithm (CaLLTiF) and state-of-the-art alternatives over simulated fMRI from the Macaque_Full connectome.** Distributions of precision for PCMCI (ignoring the contemporaneous $\circ\!\!-\!\!\circ$ connections), Mixed-PCMCI (using the contemporaneous $\circ\!\!-\!\!\circ$ connections only for adjacency), and CaLLTiF. Precisions are shown for the value of hyperparameters which give the maximum F1 score.

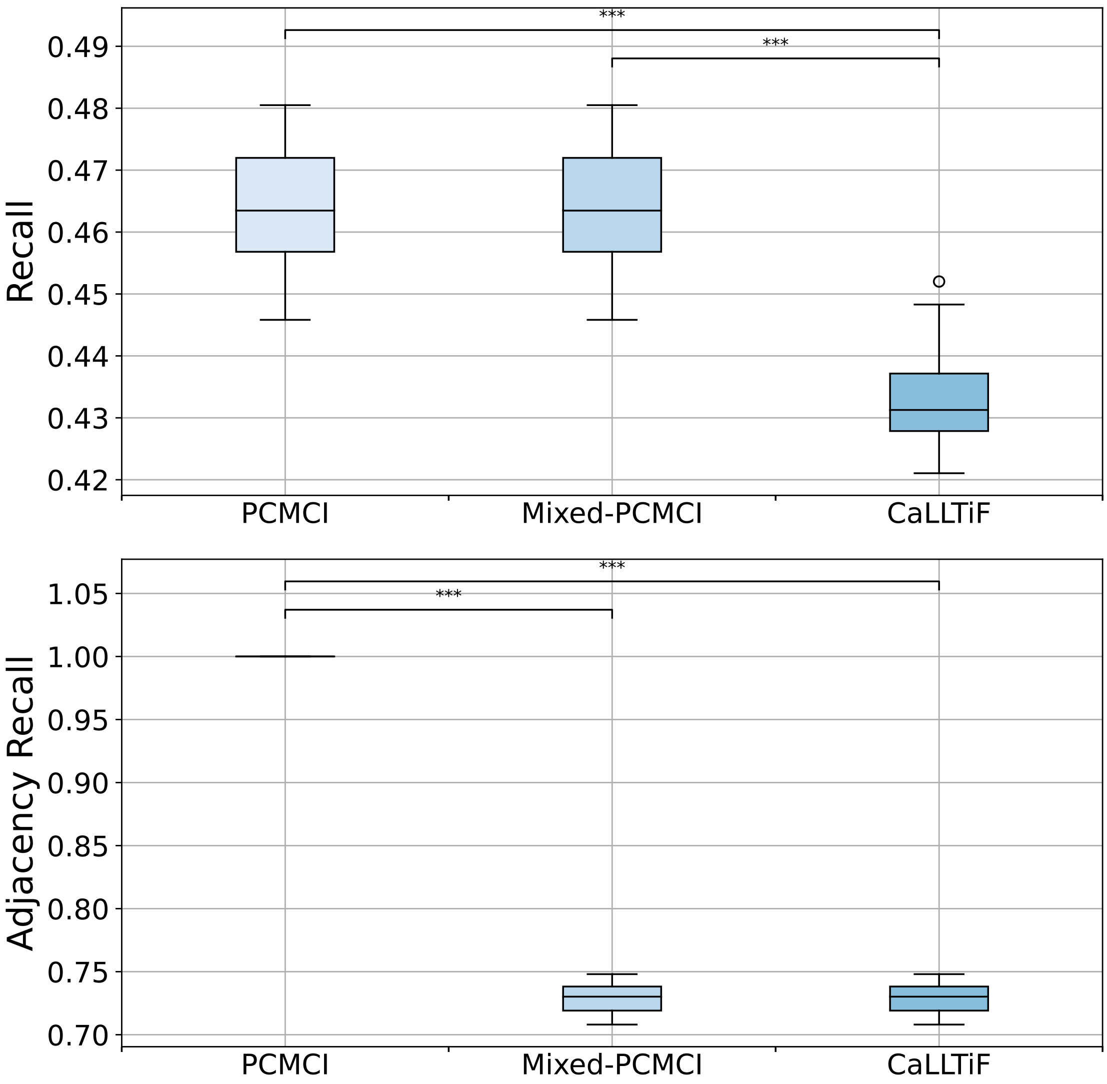

**Supplementary Figure 30: Comparisons between the proposed algorithm (CaLLTiF) and state-of-the-art alternatives over simulated fMRI from the Macaque_Full connectome.** Distributions of recall for PCMCI (ignoring the contemporaneous ⊶ connections), Mixed-PCMCI (using the contemporaneous ⊶ connections only for adjacency), and CaLLTiF. Recalls are shown for the value of hyperparameters which give the maximum F1 score.

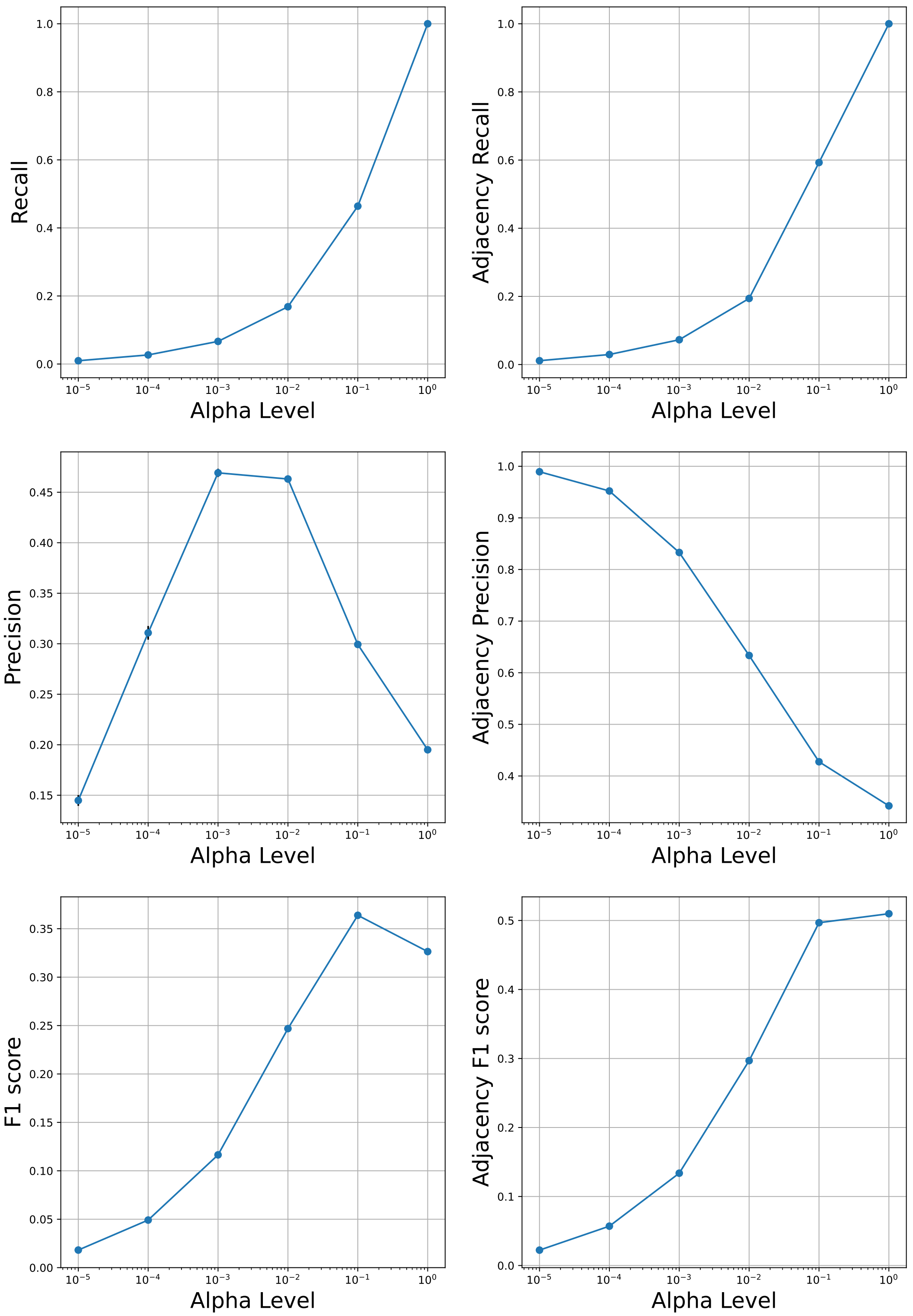

**Supplementary Figure 31: Detailed performance curves of PCMCI over simulated fMRI from Macaque_Full network for varying values of its hyperparameter Alpha Level.** In all graphs, the error bars depict the standard error of the mean.

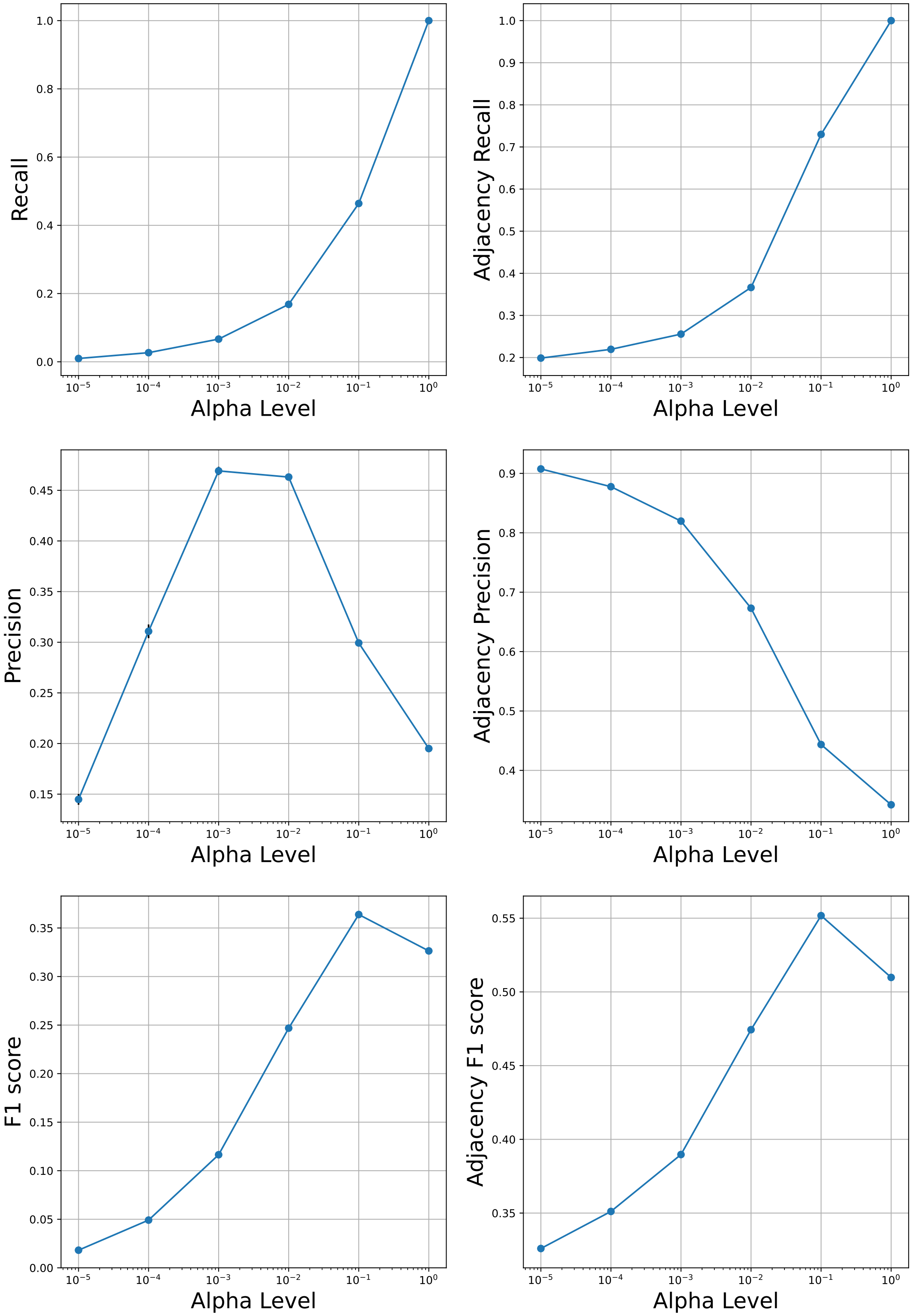

**Supplementary Figure 32: Detailed performance curves of Mixed-PCMCI over simulated fMRI from Macaque_Full network for varying values of its hyperparameter Alpha Level.** In all graphs, the error bars depict the standard error of the mean.

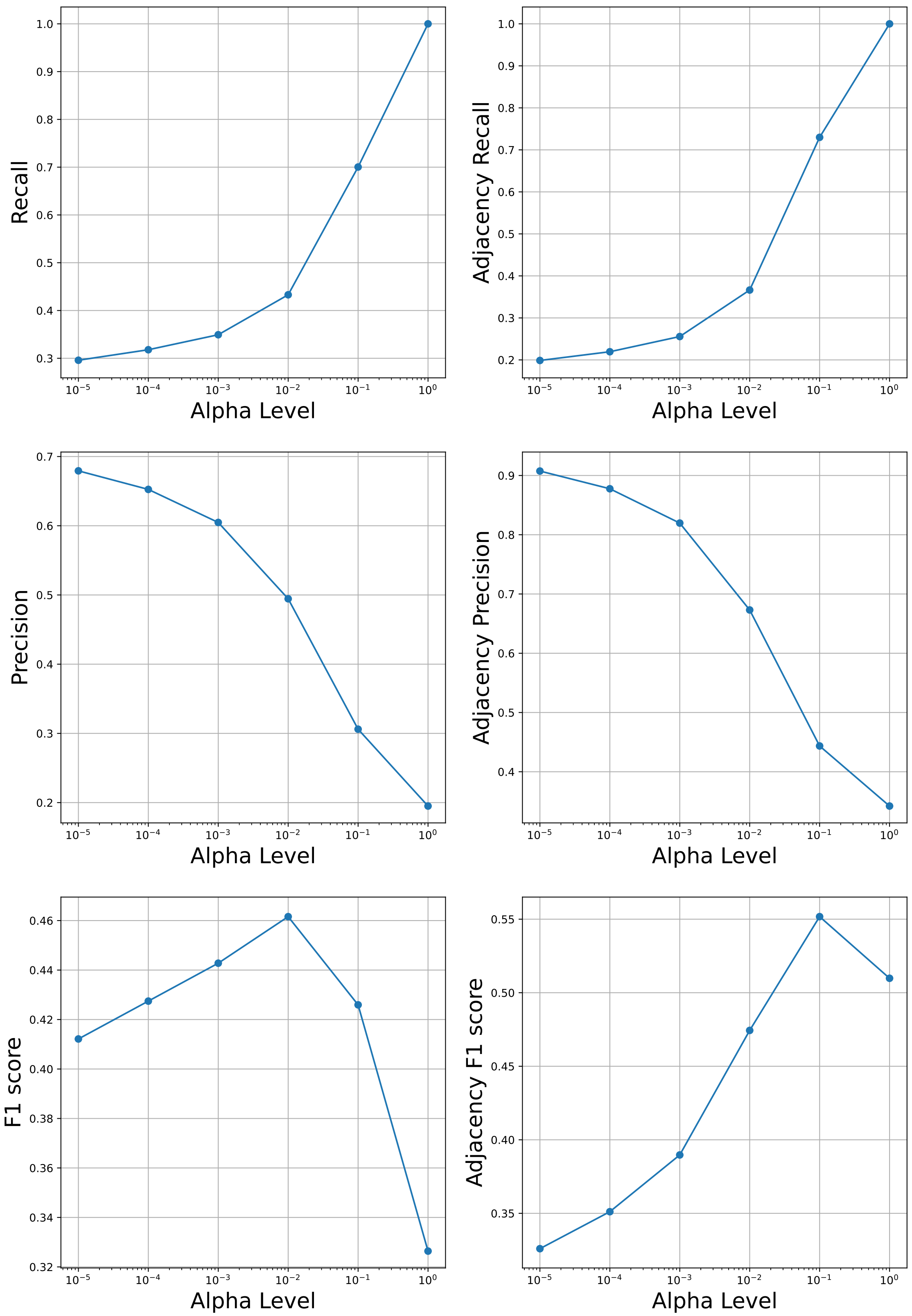

**Supplementary Figure 33: Detailed performance curves of CaLLTiF over simulated fMRI from Macaque_Full network for varying values of its hyperparameter Alpha Level.** In all graphs, the error bars depict the standard error of the mean.

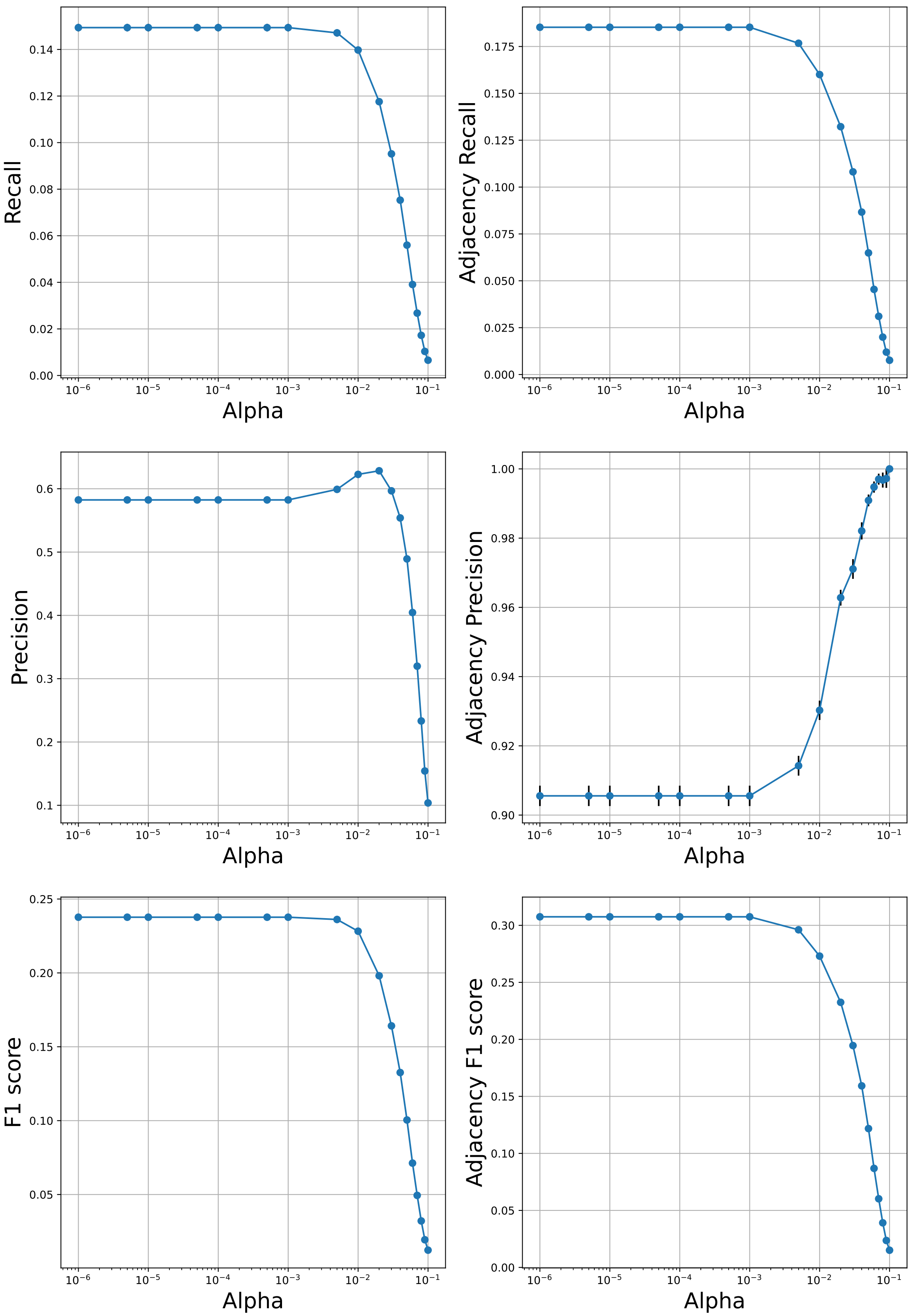

**Supplementary Figure 34: Detailed performance curves of VARLiNGAM over simulated fMRI from Macaque_Full network for varying values of its hyperparameter Alpha.** In all graphs, the error bars depict the standard error of the mean.

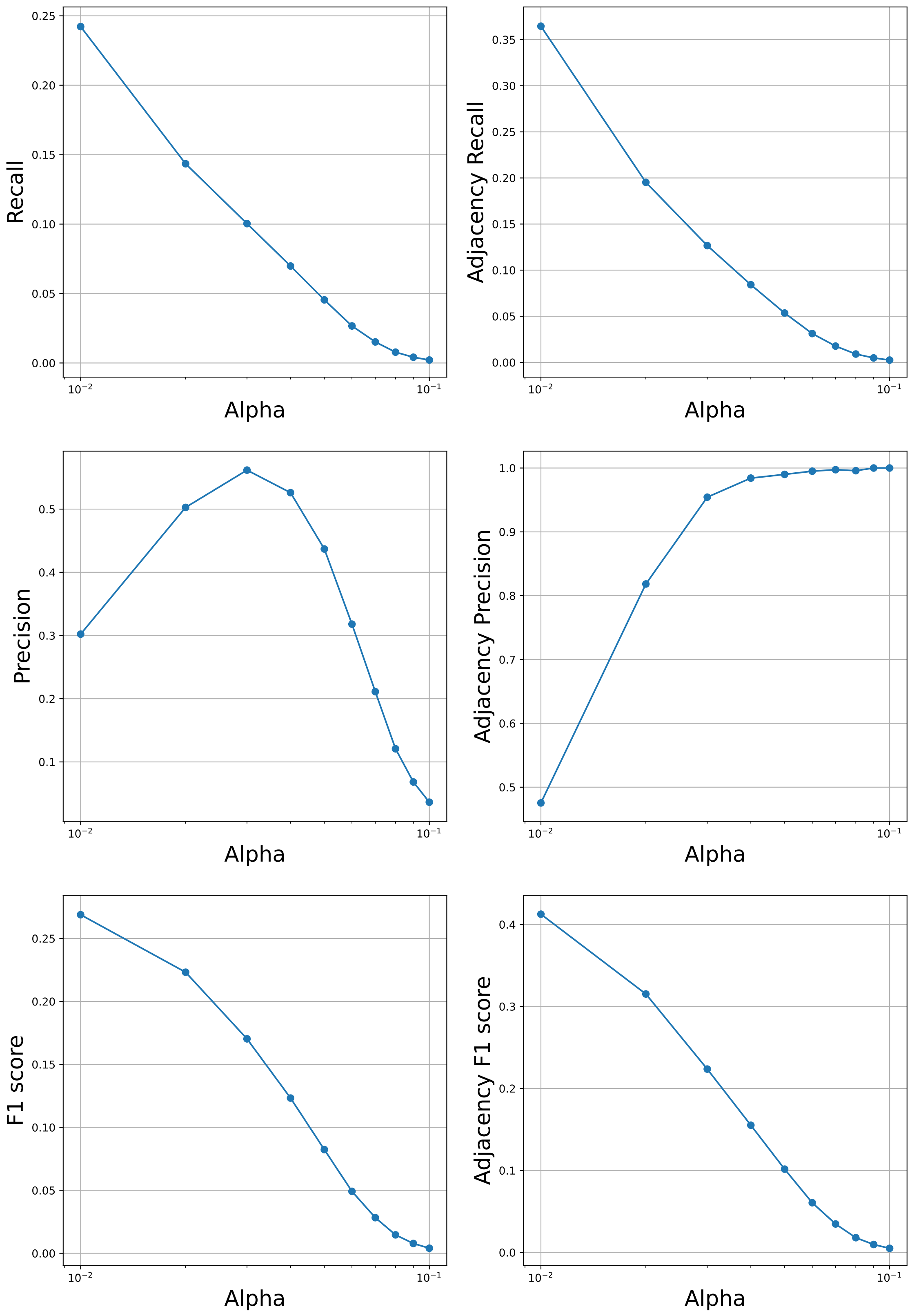

**Supplementary Figure 35: Detailed performance curves of DYNOTEARS over simulated fMRI from Macaque_Full network for varying values of its hyperparameter Alpha.** In all graphs, the error bars depict the standard error of the mean.

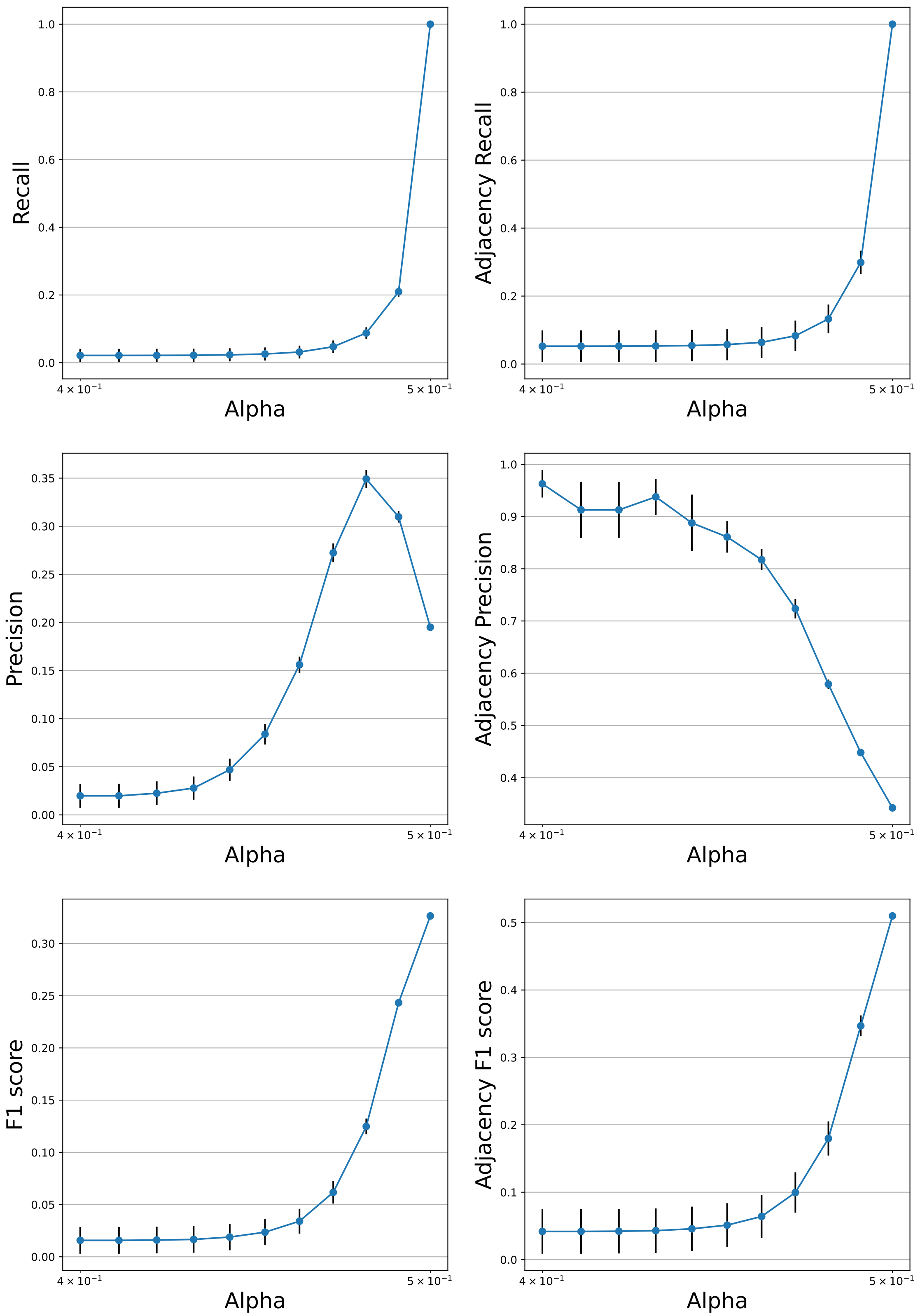

**Supplementary Figure 36: Detailed performance curves of MVGC over simulated fMRI from Macaque_Full network for varying values of its hyperparameter Alpha.** In all graphs, the error bars depict the standard error of the mean.

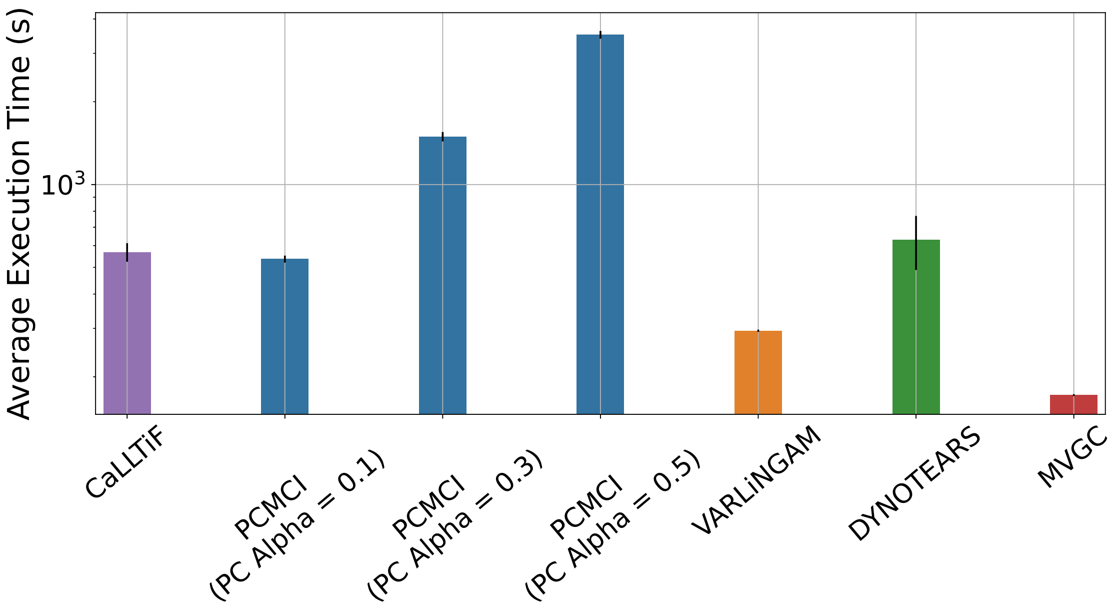

**Supplementary Figure 37: Mean execution times of CaLLTiF and state-of-the-art alternatives on 15 repetitions of Macaque_Full data**. The 'PC Alpha' hyperparameter in PCMCI significantly influences execution time, and is hence set to 3 different values for comparison. Note the logarithmic scale on the y-axis. Error bars represent one standard deviation.

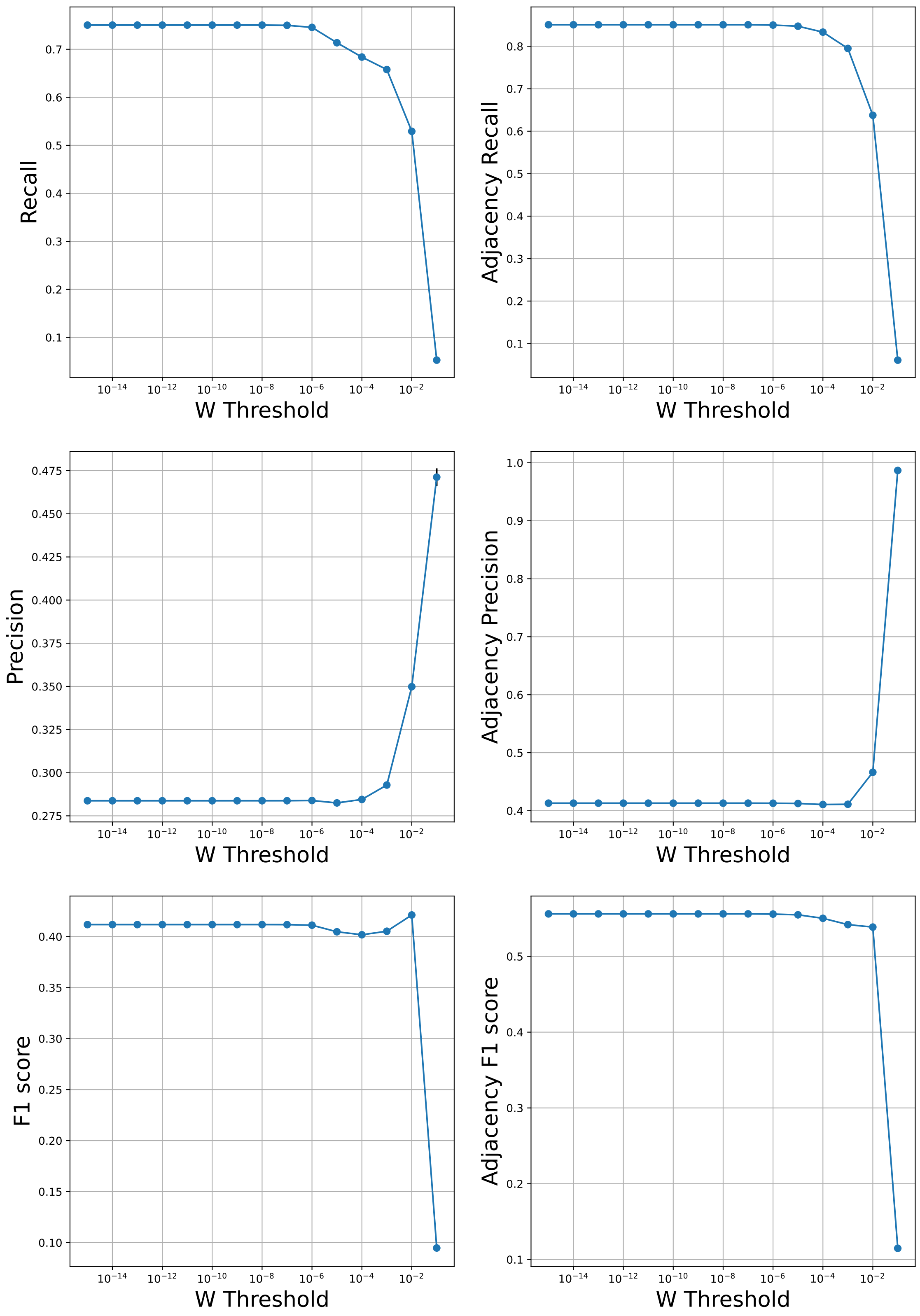

**Supplementary Figure 38: Detailed performance curves of NTS-NOTEARS over simulated fMRI from Macaque_Full network for varying values of its hyperparameter W Threshold.** In all graphs, the error bars depict the standard error of the mean.

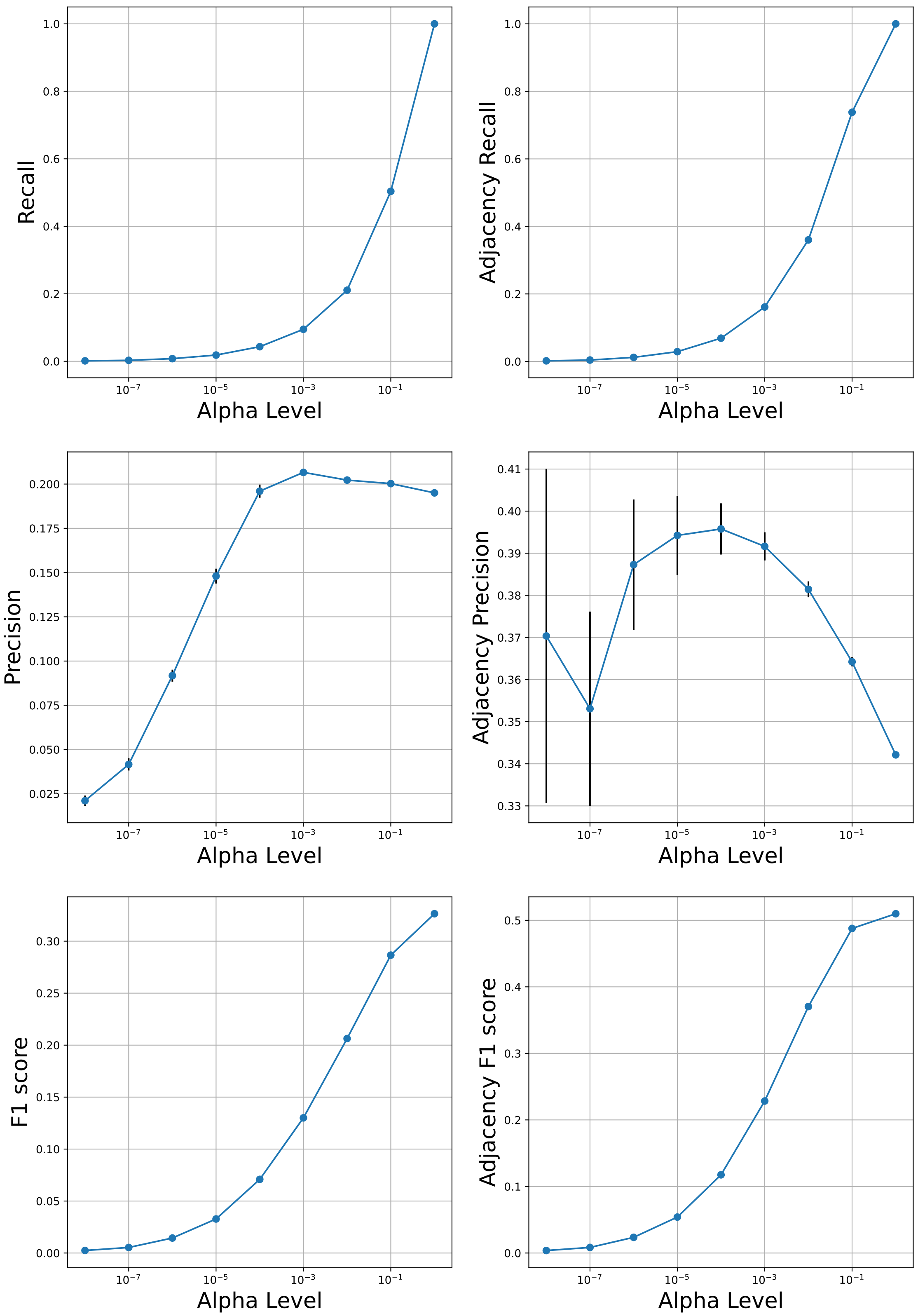

**Supplementary Figure 39: Detailed performance curves of PCMCI over simulated fMRI from Macaque_Full network when using ground-truth parent sets for each conditional independence test.** In all graphs, the error bars depict the standard error of the mean.

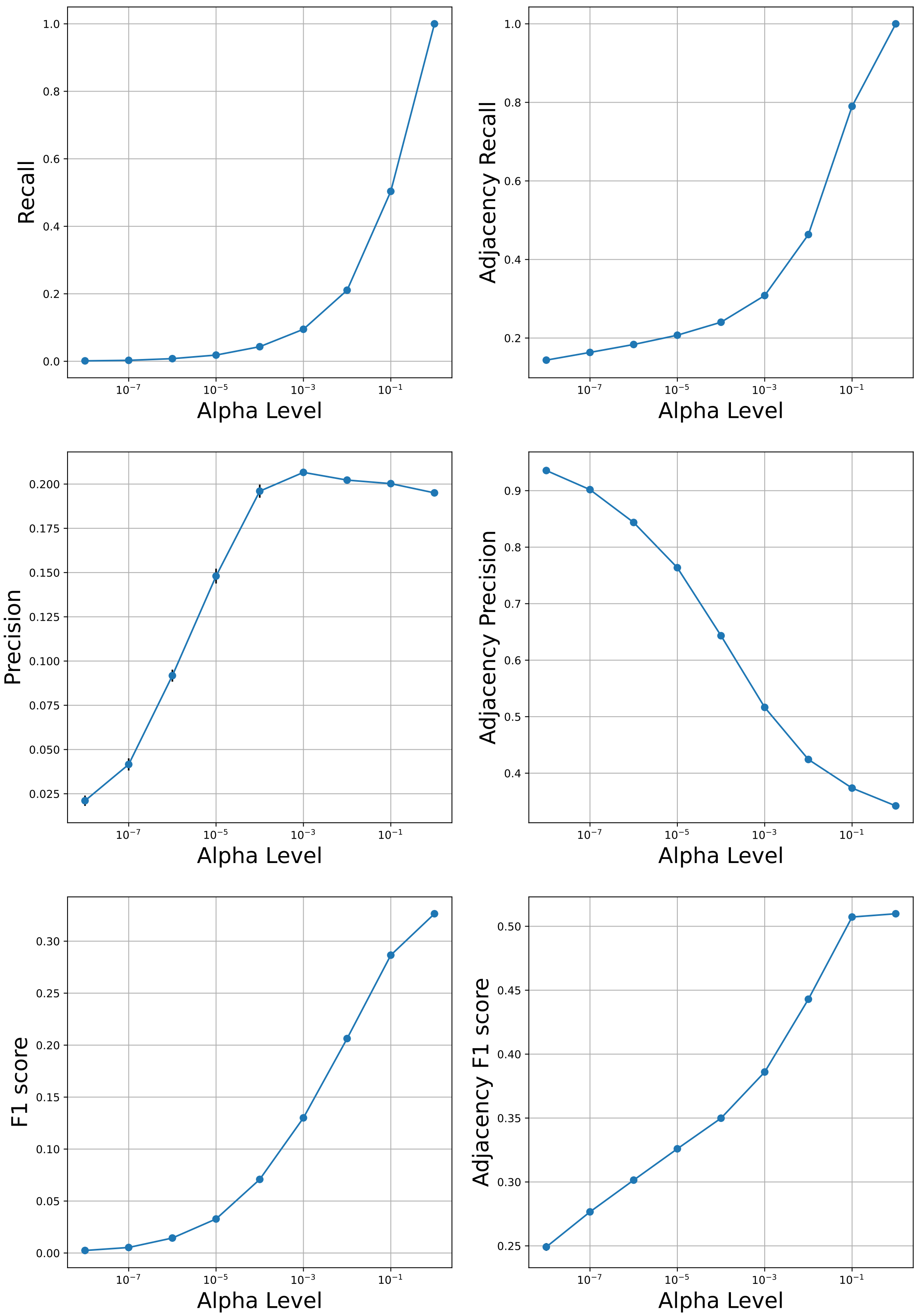

**Supplementary Figure 40: Detailed performance curves of Mixed-PCMCI over simulated fMRI from Macaque_Full network when using ground-truth parent sets for each conditional independence test.** In all graphs, the error bars depict the standard error of the mean.

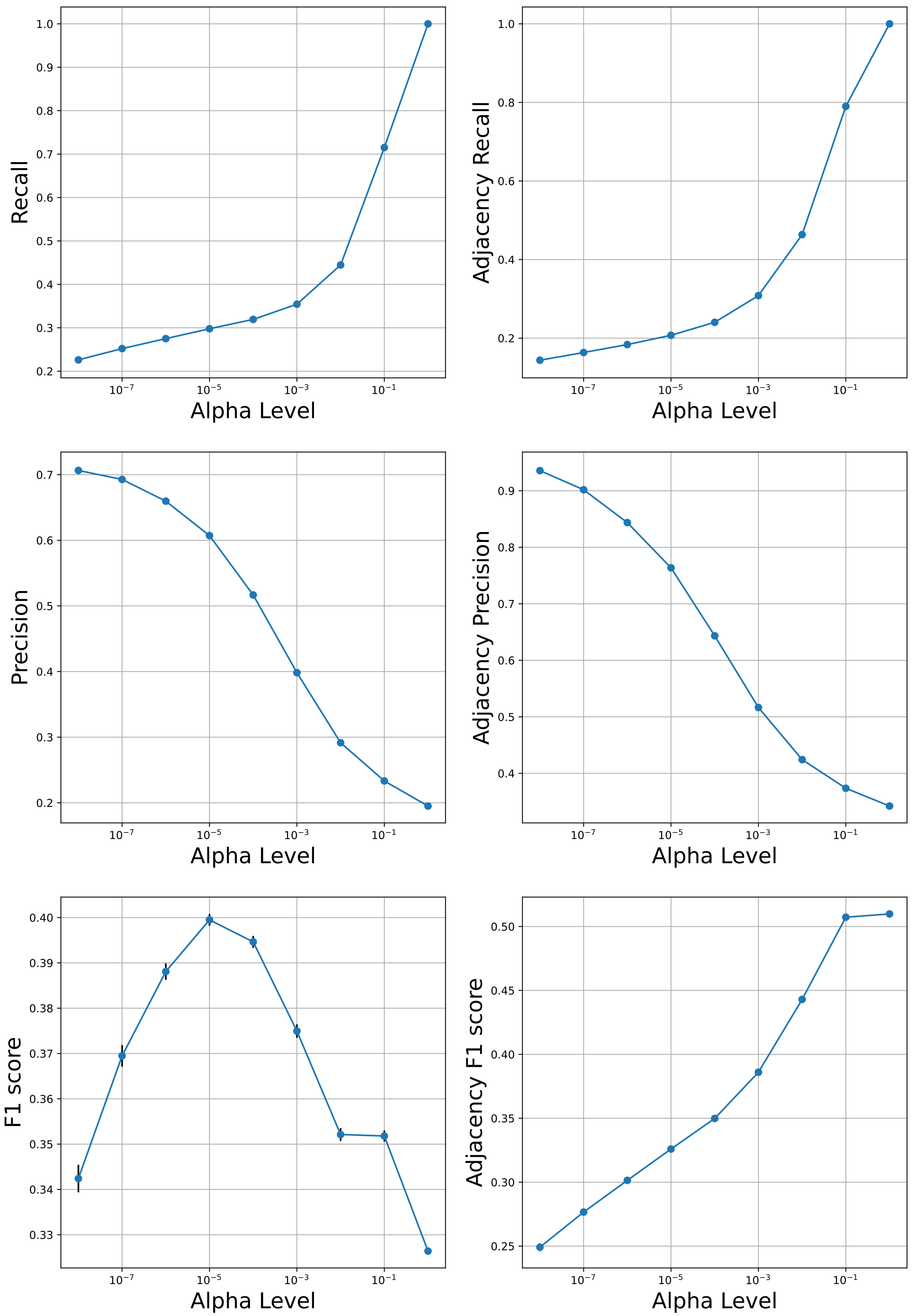

**Supplementary Figure 41: Detailed performance curves of CaLLTiF over simulated fMRI from Macaque_Full network when using ground-truth parent sets for each conditional independence test.** In all graphs, the error bars depict the standard error of the mean.

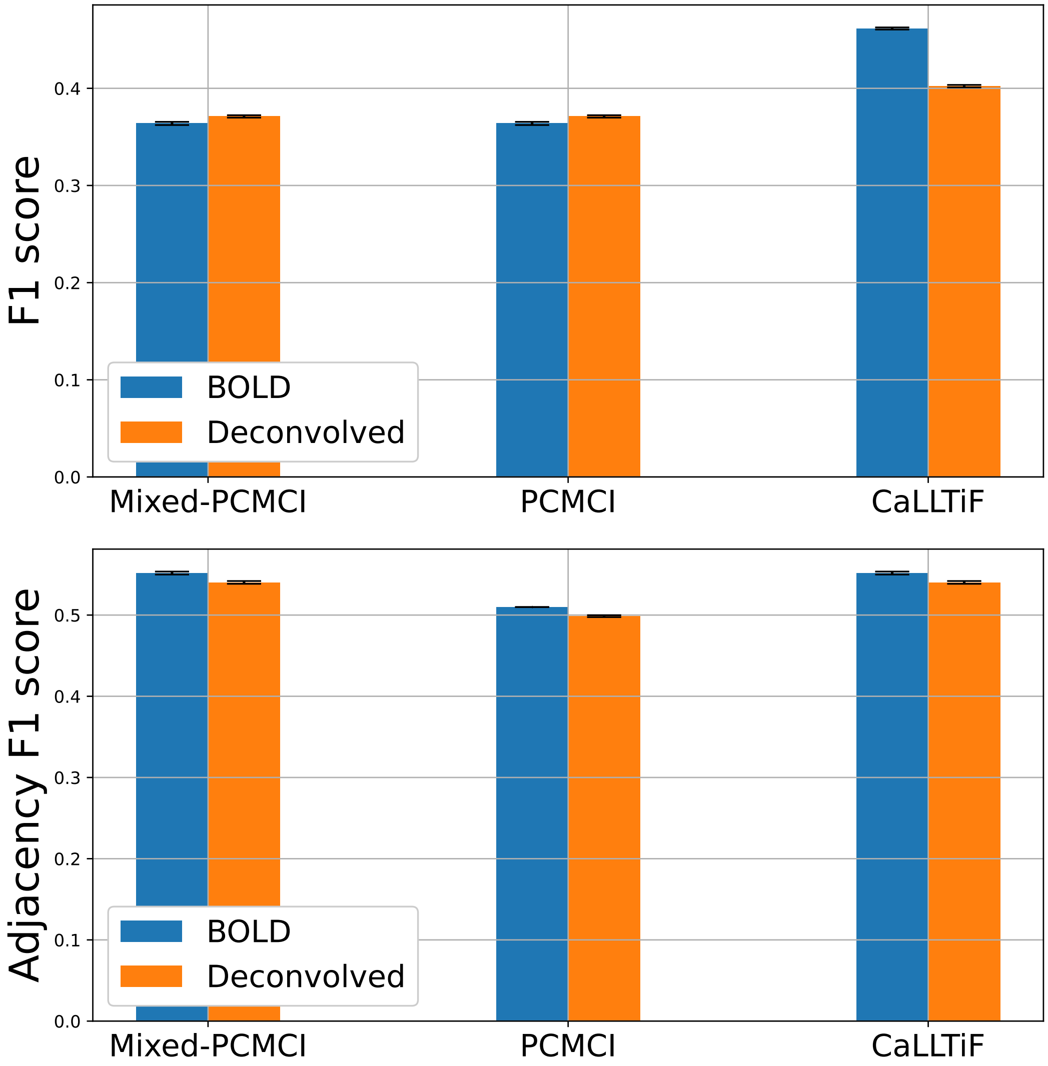

**Supplementary Figure 42: Effect of HRF deconvolution on causal discovery.** Bar plots show comparison between F1 scores and adjacency F1 scores of CaLLTiF, PCMCI, and Mixed-PCMCI when applied to simulated fMRI from the Macaque_Full network with and without deconvolution with a generic hemodynamic response function (HRF). In all graphs, the error bars depict one standard deviation.

### Supplementary Figures for Resting-State Human fMRI from HCP

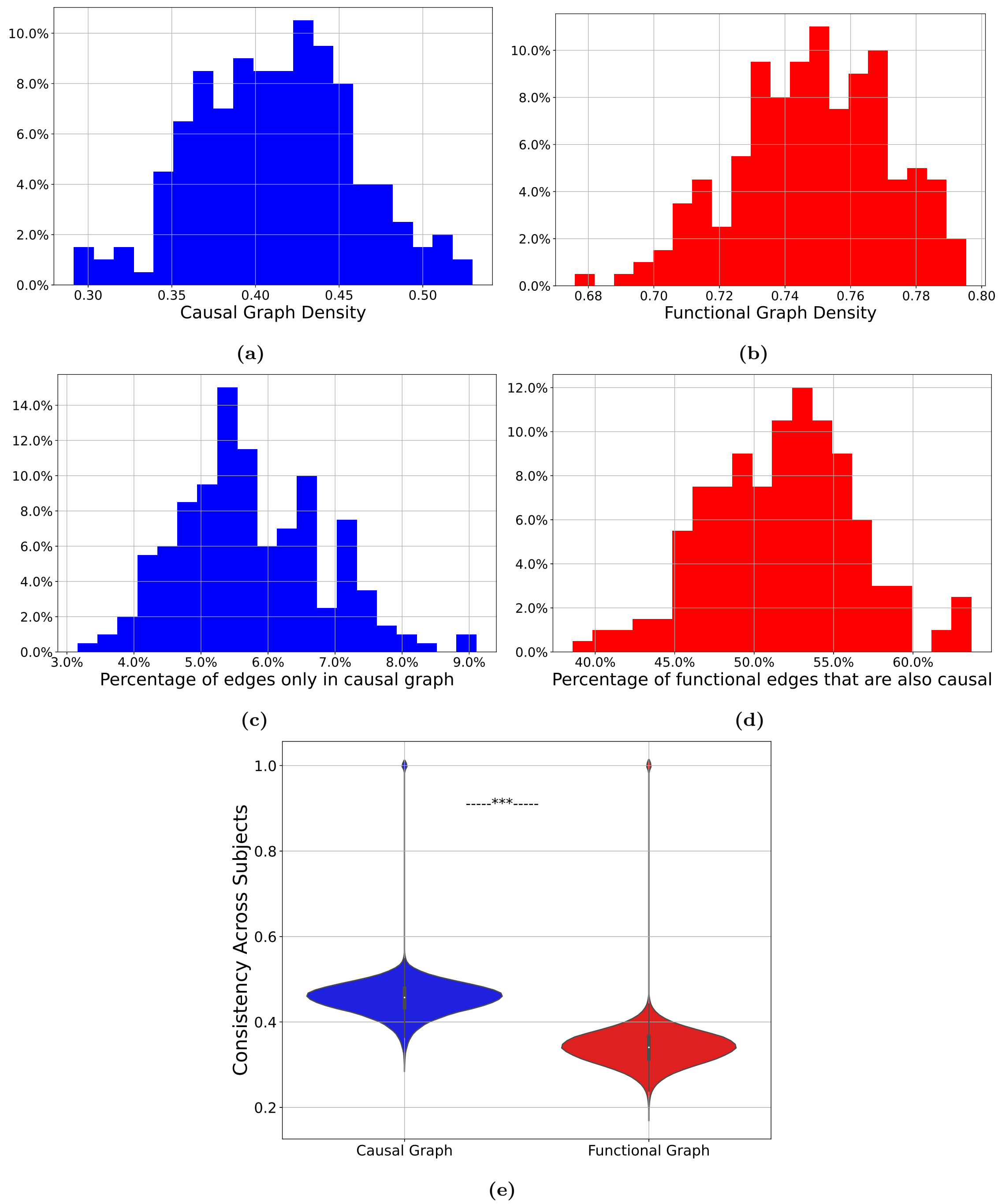

**Supplementary Figure 43: Causal graphs are sparse and more consistent than functional connectivity (FC) graphs.** (a) Distribution of the density of edges in causal graphs learned by CaLLTiF. (b) Similar to (a) but for FC. (c) Distribution of the percentage of edges that only exist in CaLLTiF but not in FC. (d) Distribution of the percentage of edges that exist in both causal and functional graphs. (e) Distributions of graph consistencies (correlation coefficients) across subjects. $^{***} = p < 0.001$, one-sided Wilcoxon signed-rank test.

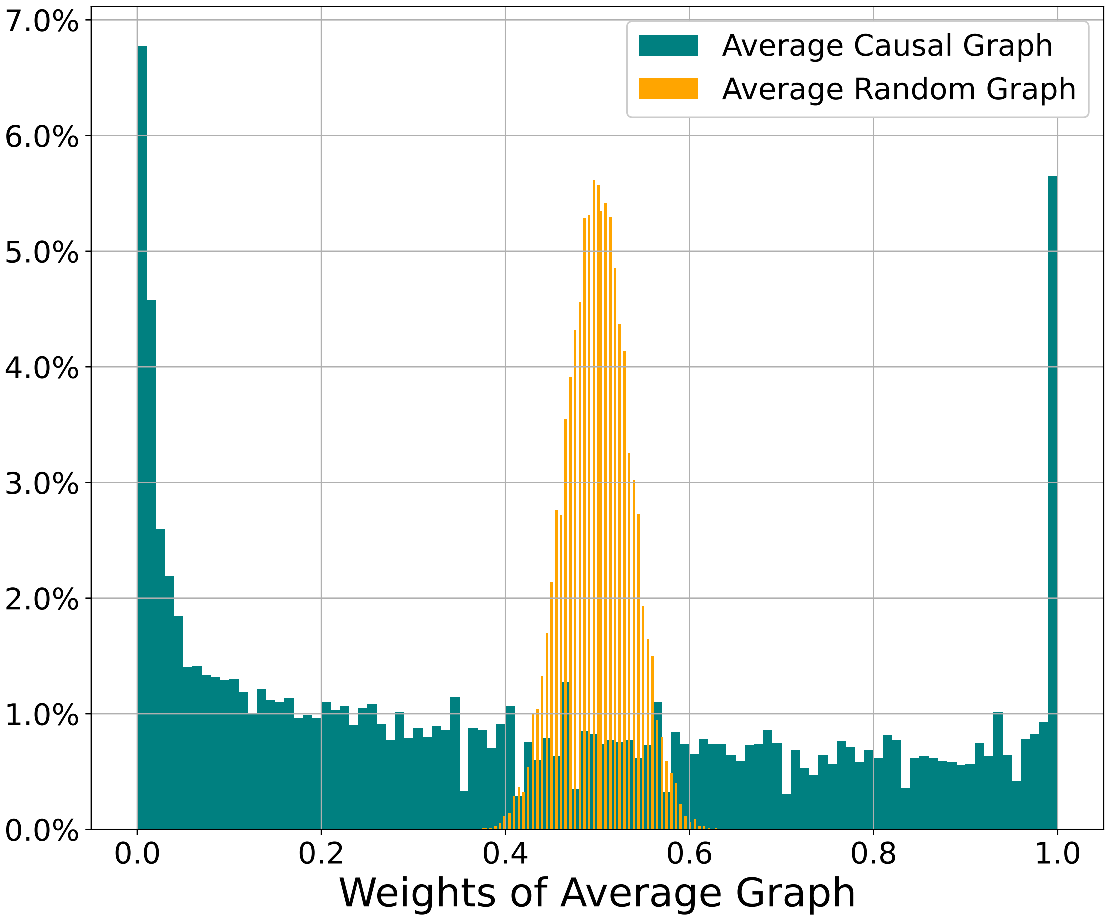

**Supplementary Figure 44: Distributions of edge weights of the average causal graph (computed for 200 HCP subjects) and corresponding weights of the average of 200 random binary graphs.** The randomized surrogate is computed based on data from 200 subjects, with the weight distribution of the average random graph, which is computed based on data from 200 random binary matrices from the Bernoulli distribution with $p = 0.5$. The average graph based on real data is significantly more bimodal ($p = 0$, Kolmogorov-Smirnov test), indicating the presence of evidence in the data on whether each edge exists or not.

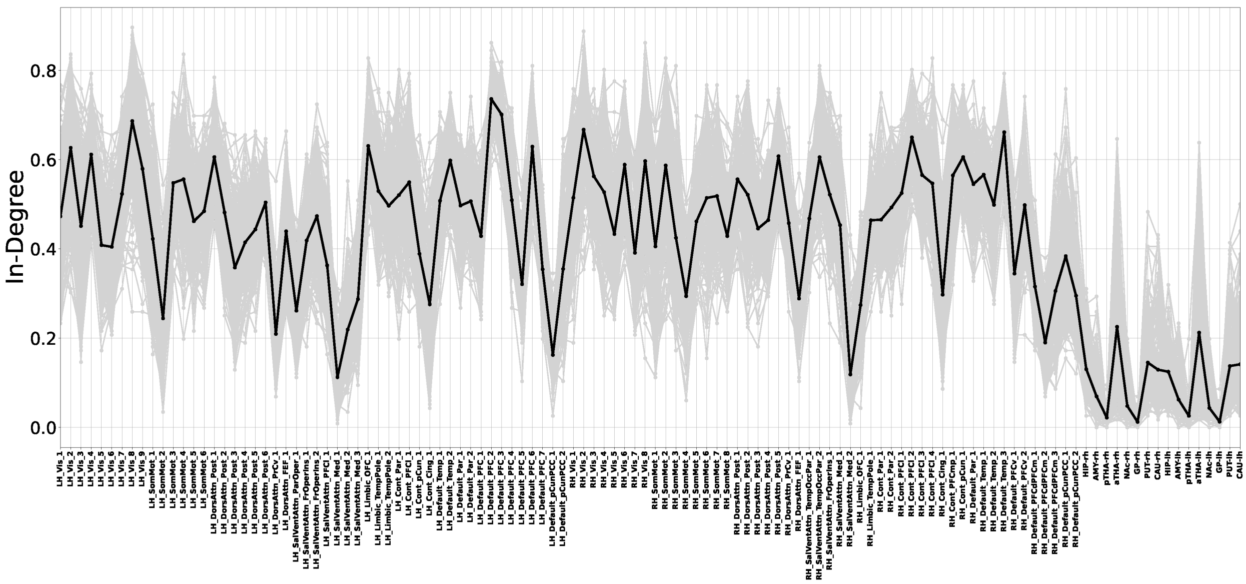

**Supplementary Figure 45: In-degree for all nodes (horizontal axis) and all subjects (gray lines) in causal graphs learned by CaLLTiF over human fMRI data from HCP.**

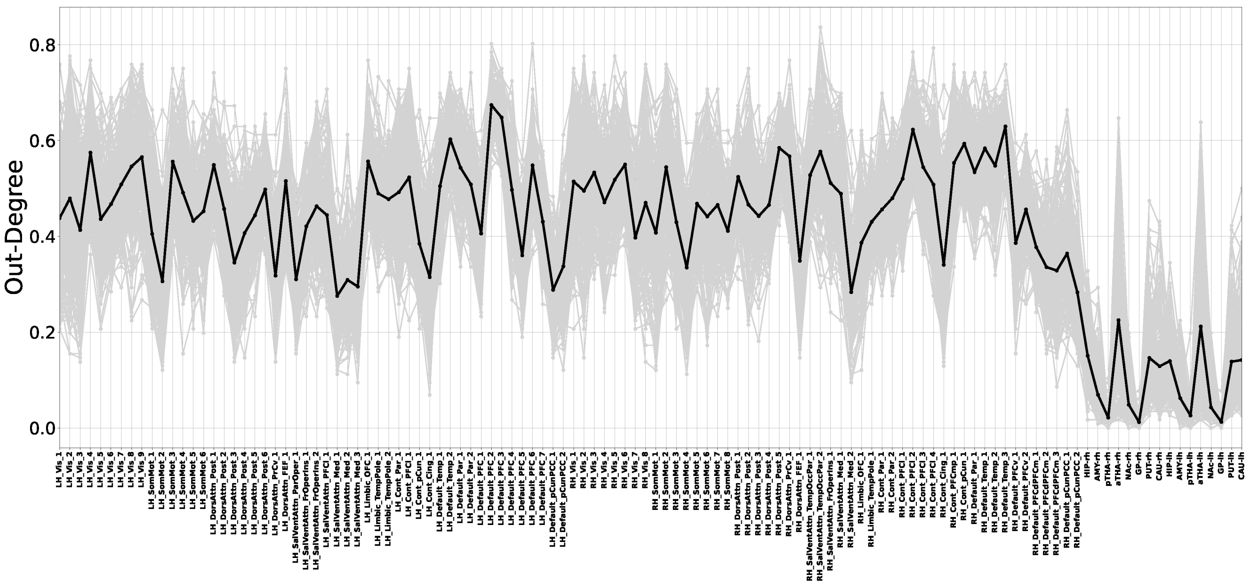

**Supplementary Figure 46: Out-degree for all nodes (horizontal axis) and all subjects (gray lines) in causal graphs learned by CaLLTiF over human fMRI data from HCP.**

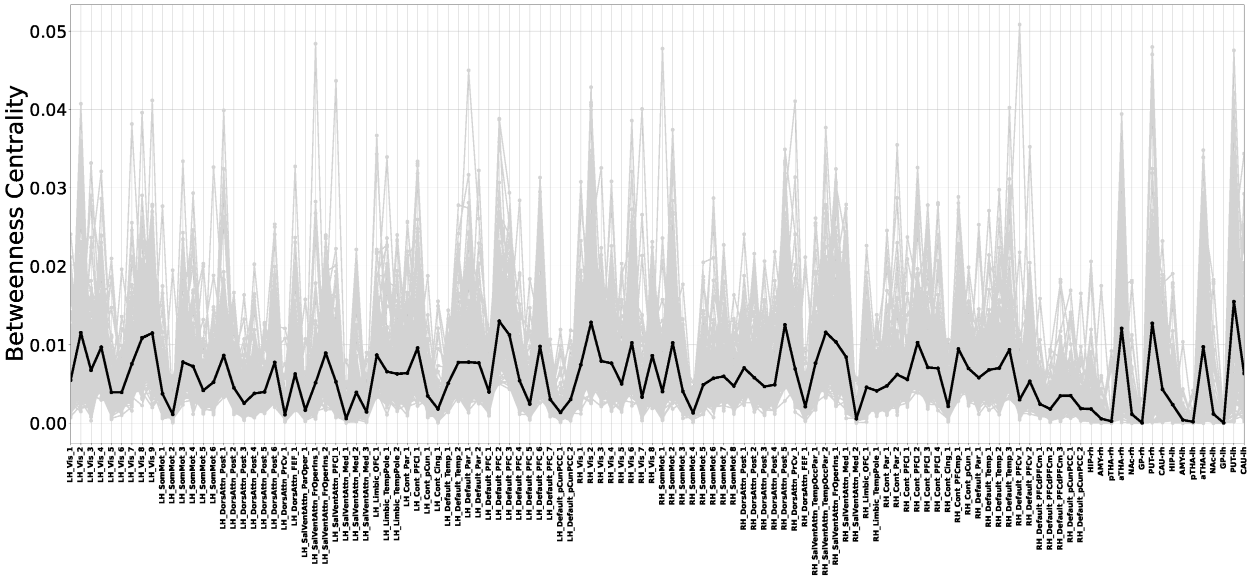

**Supplementary Figure 47: Betweenness centrality for all nodes (horizontal axis) and all subjects (gray lines) in causal graphs learned by CaLLTiF over human fMRI data from HCP.**

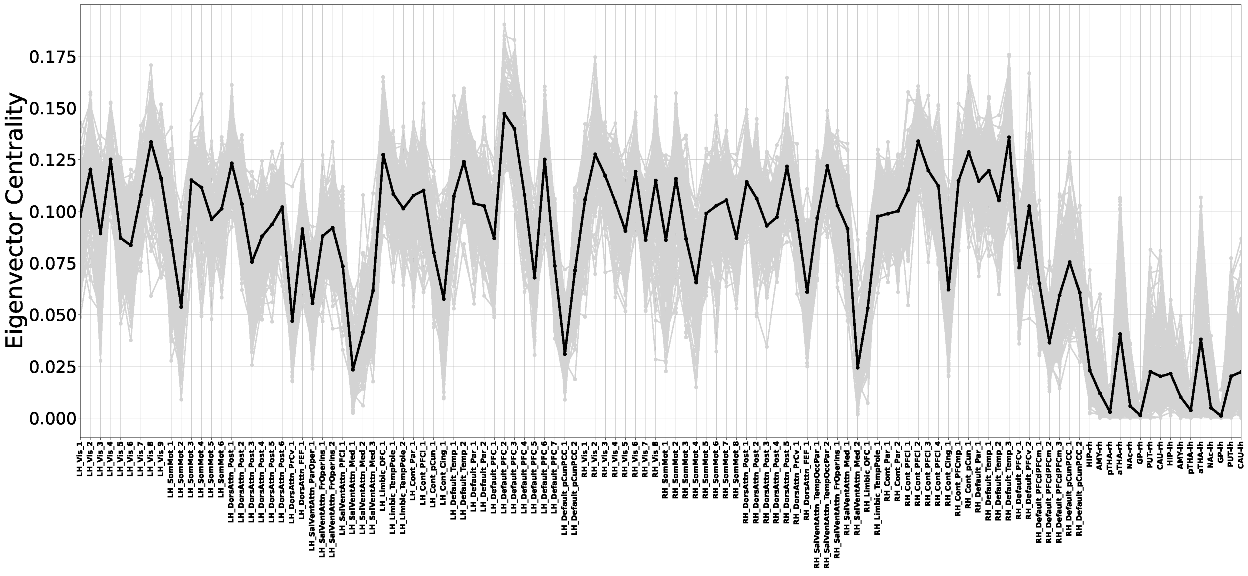

**Supplementary Figure 48: Eigenvector centrality for all nodes (horizontal axis) and all subjects (gray lines) in causal graphs learned by CaLLTiF over human fMRI data from HCP.**

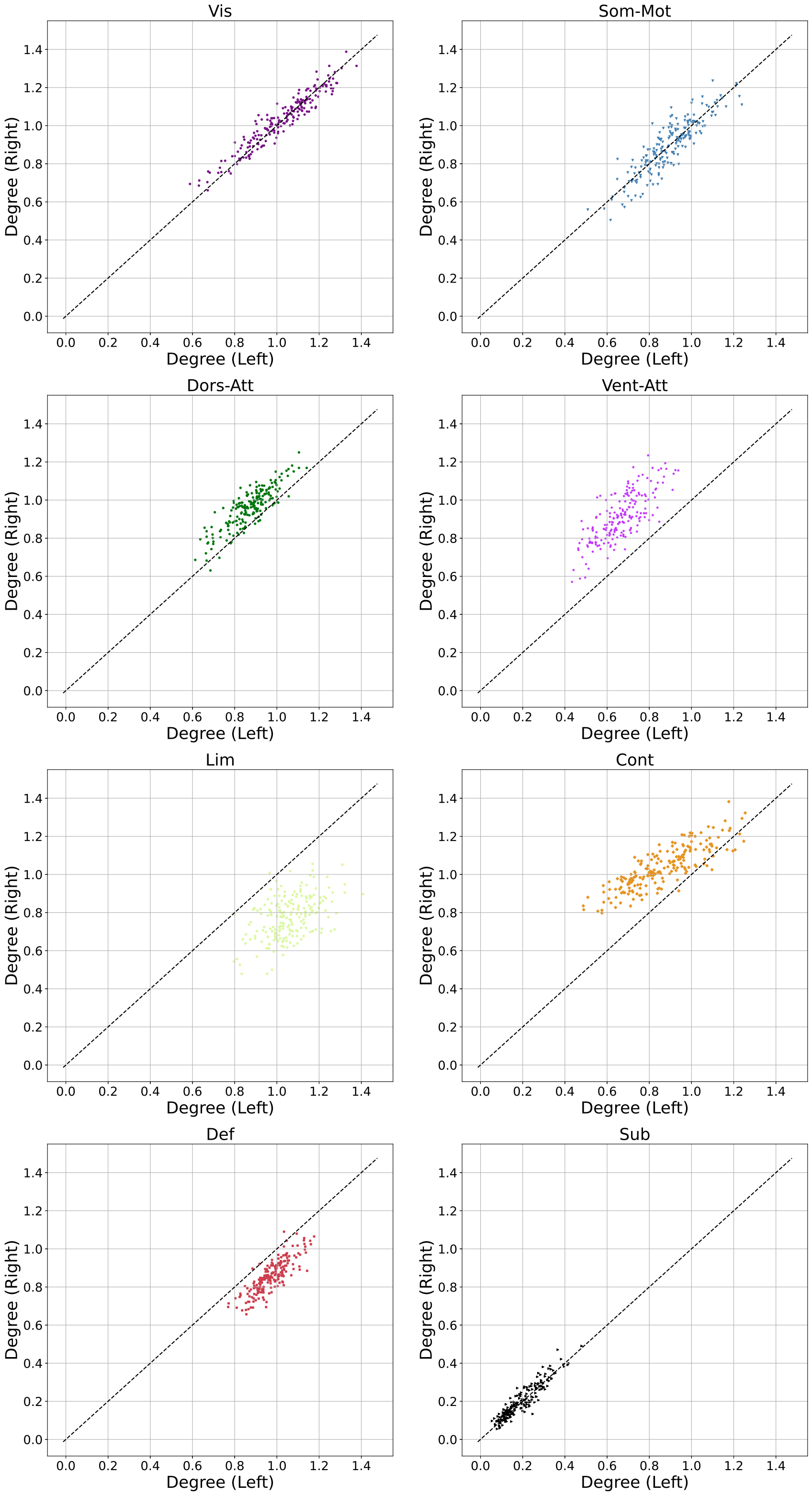

**Supplementary Figure 49: Hemispheric (a)symmetry of nodal degrees in causal graphs learned by CaLLTiF over human fMRI data from HCP.** This is the same as Figure 8c in the main text except that different subnetworks are shown in distinct panels for better visualization.

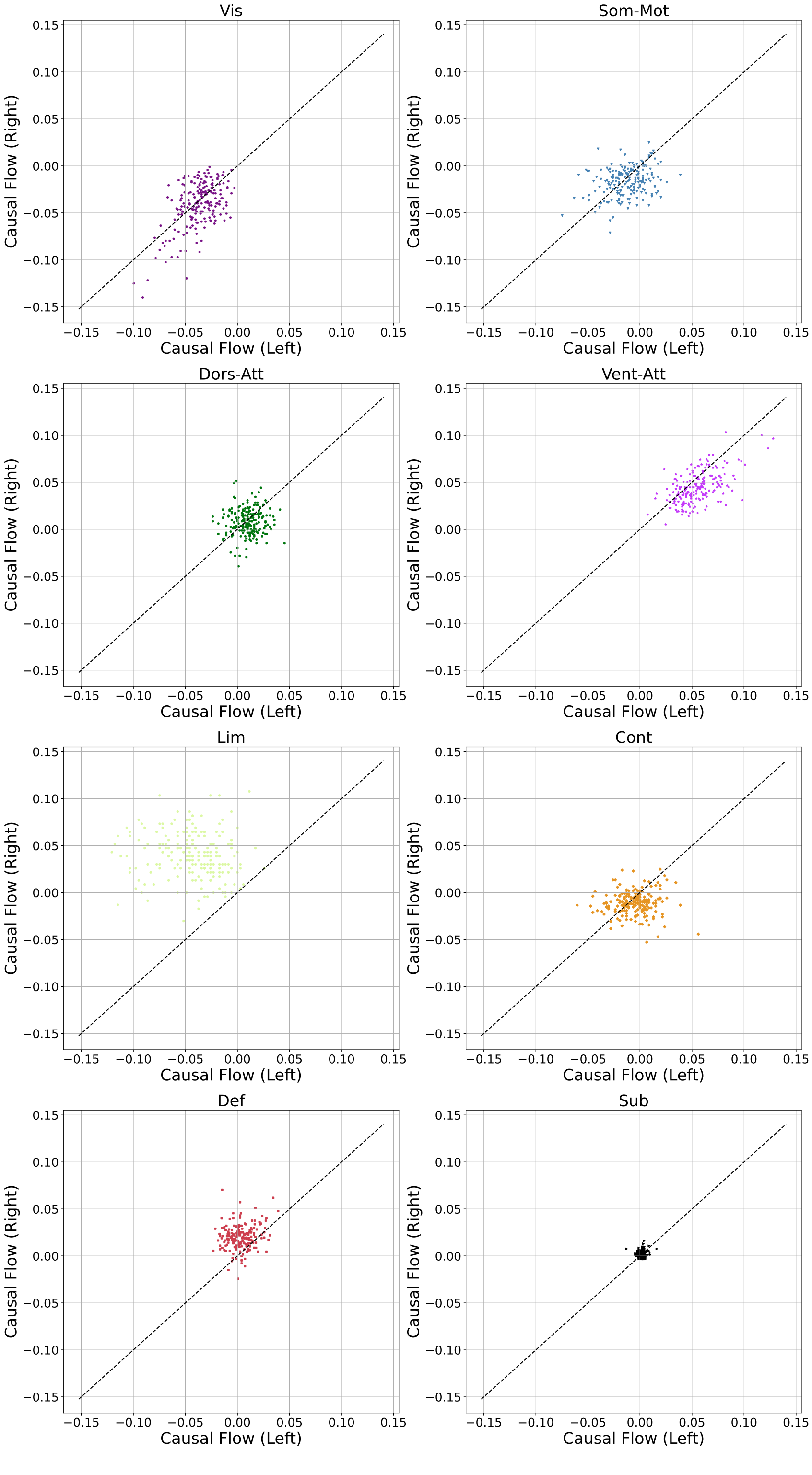

**Supplementary Figure 50: Hemispheric (a)symmetry of causal flows in causal graphs learned by CaLLTiF over human fMRI data from HCP.** This is the same as Figure 8d in the main text except that different subnetworks are shown in distinct panels for better visualization.

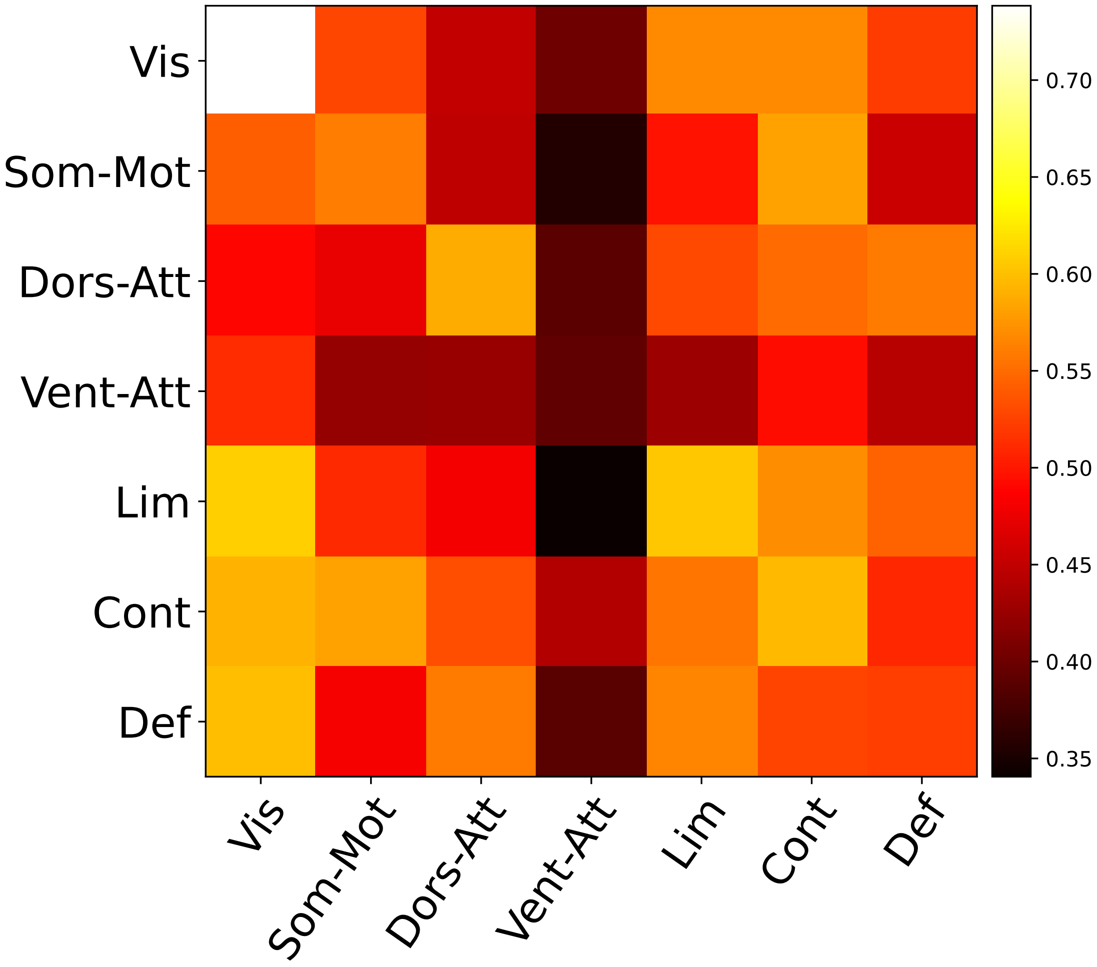

**Supplementary Figure 51: The average subnetwork graph, computed as the mean of subnetwork graphs of all the subjects.** In the subnetwork graph of each subject, the weight of an edge from subnetwork $i$ to $j$ is the number of nodes in subnetwork $i$ that connect to nodes in subnetwork $j$, normalized by the number of all possible edges between these subnetworks.

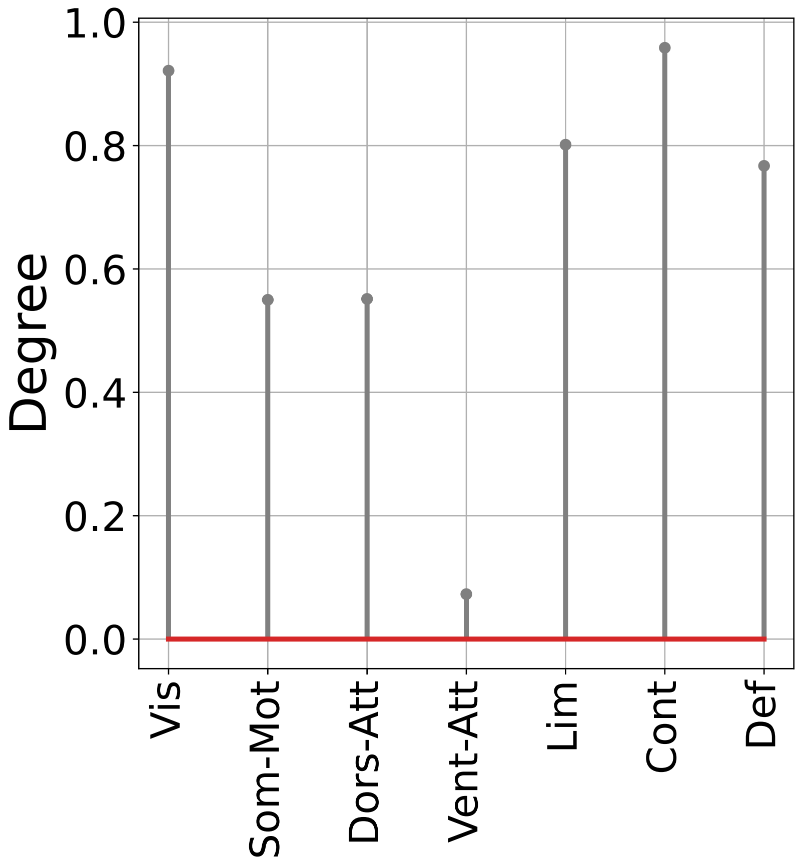

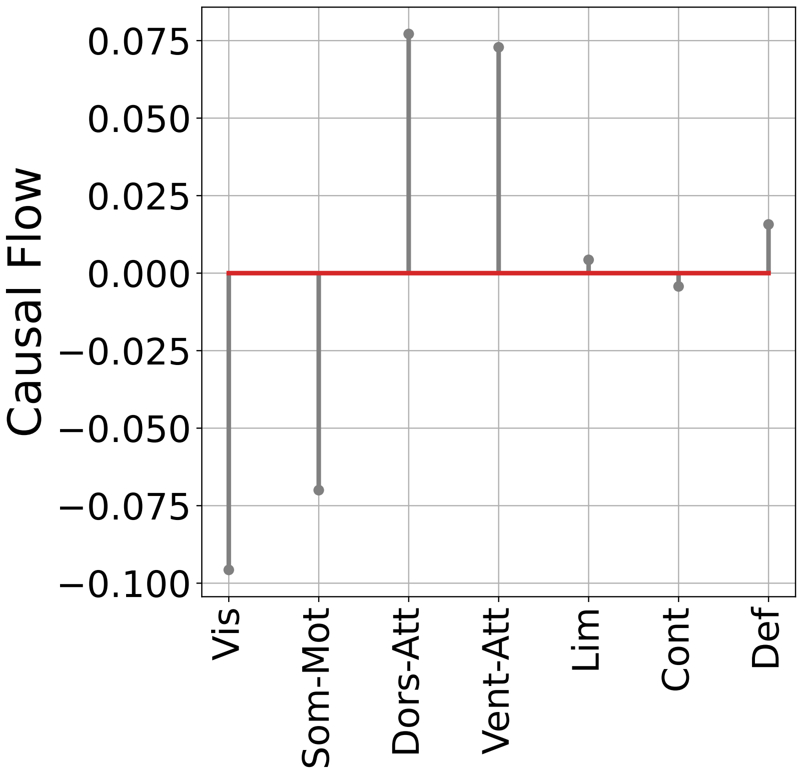

**Supplementary Figure 52: Nodal degree and causal flow for each node in the average subnetwork graph in Supplementary Figure 51**

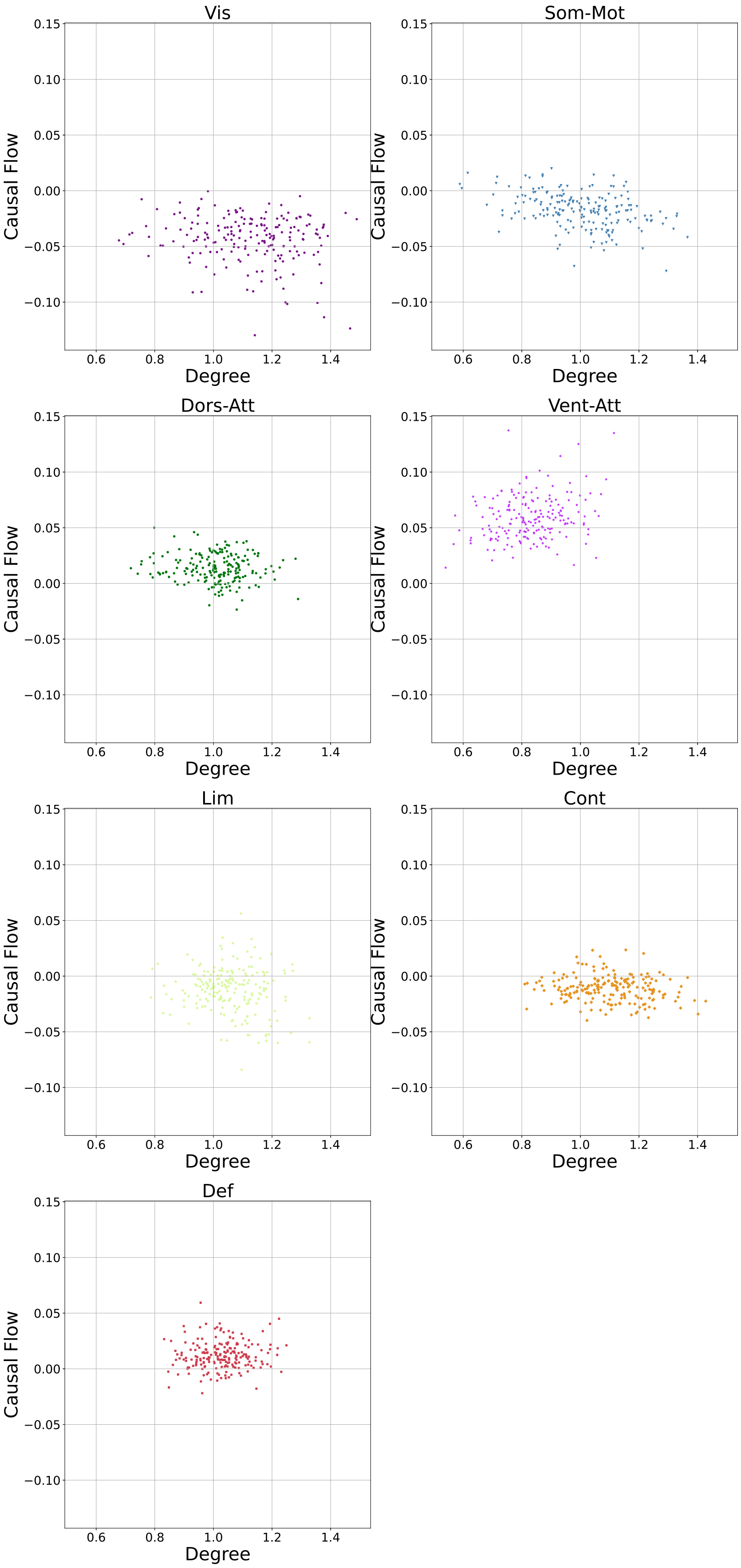

**Supplementary Figure 53: Joint distribution of degree and causal flow for (hyper-)nodes of subnetwork graphs for each subject.** Each dot in each panel represents one subject and different panels show different functional subnetworks. This is the same as Figure 5b in the main text except that different subnetworks are shown in distinct panels for better visualization.

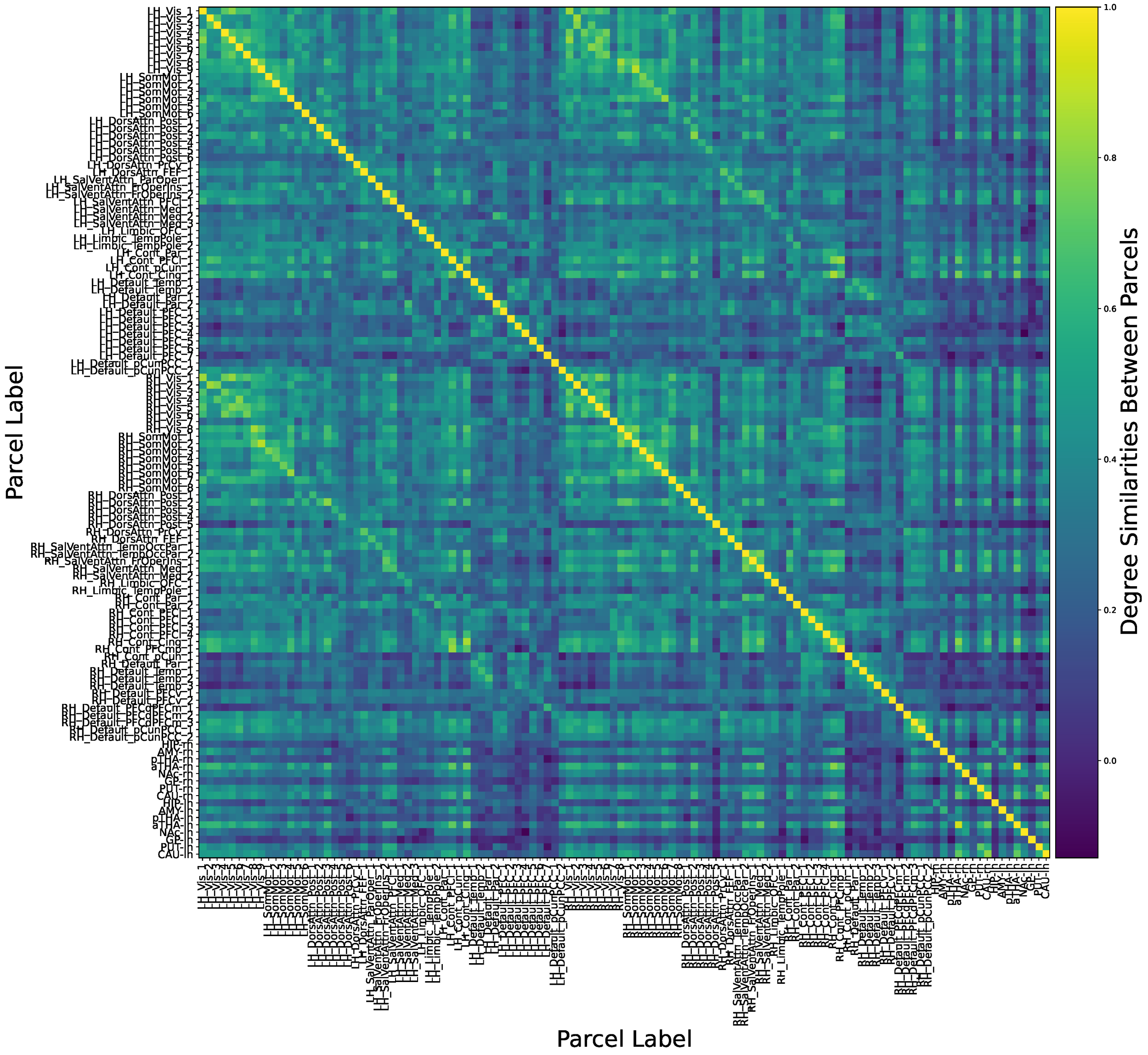

**Supplementary Figure 54: Degree similarities between each pair of nodes (parcels) in causal graphs learned by CaLLTiF over human fMRI data from HCP.** Each cell shows the correlation coefficient between nodal degrees of the respective two nodes across different subjects.

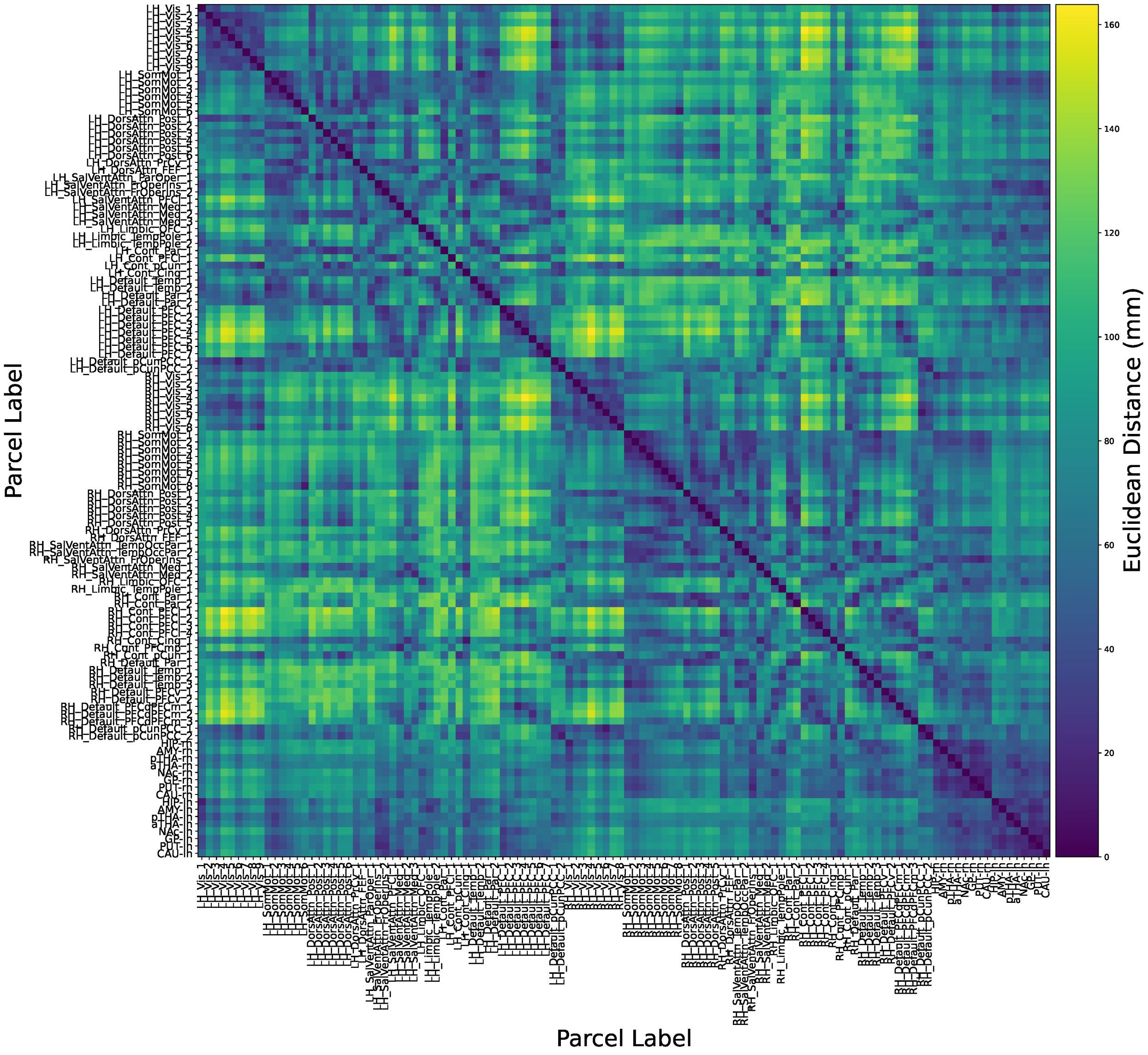

**Supplementary Figure 55: Pairwise spatial (Euclidean) distances between all nodes (parcels) of the causal graphs learned by CaLLTiF over human fMRI data from HCP.**

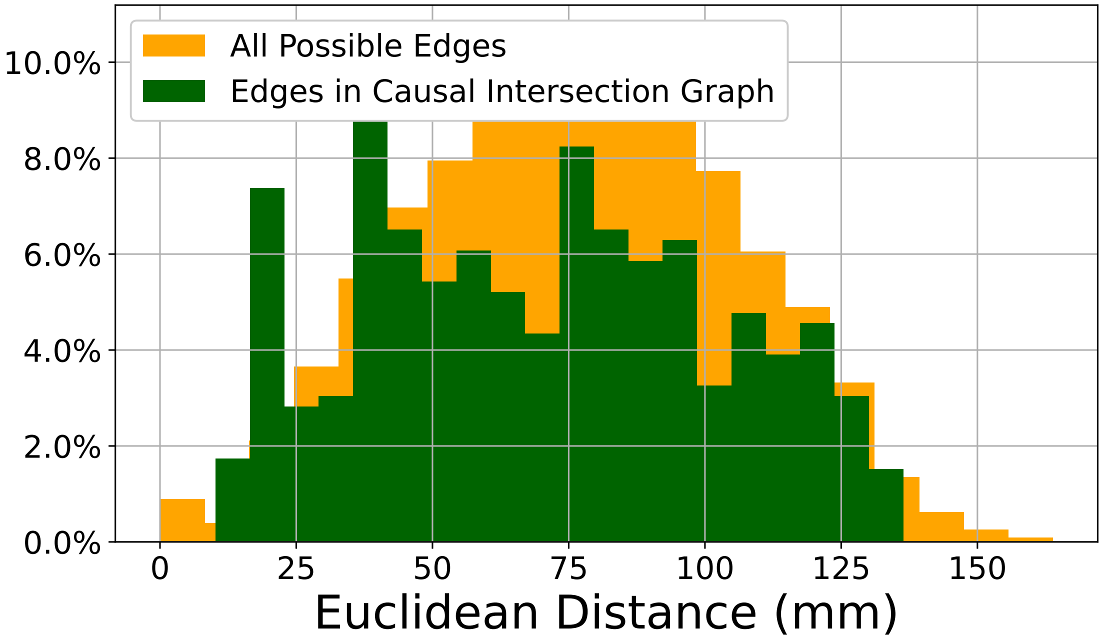

**Supplementary Figure 56: Distribution of Euclidean distances for each possible pair of nodes, compared to the distribution of Euclidean distances only for links present in the intersection of all subject-wise causal graphs learned by CaLLTiF over human fMRI data from HCP.**

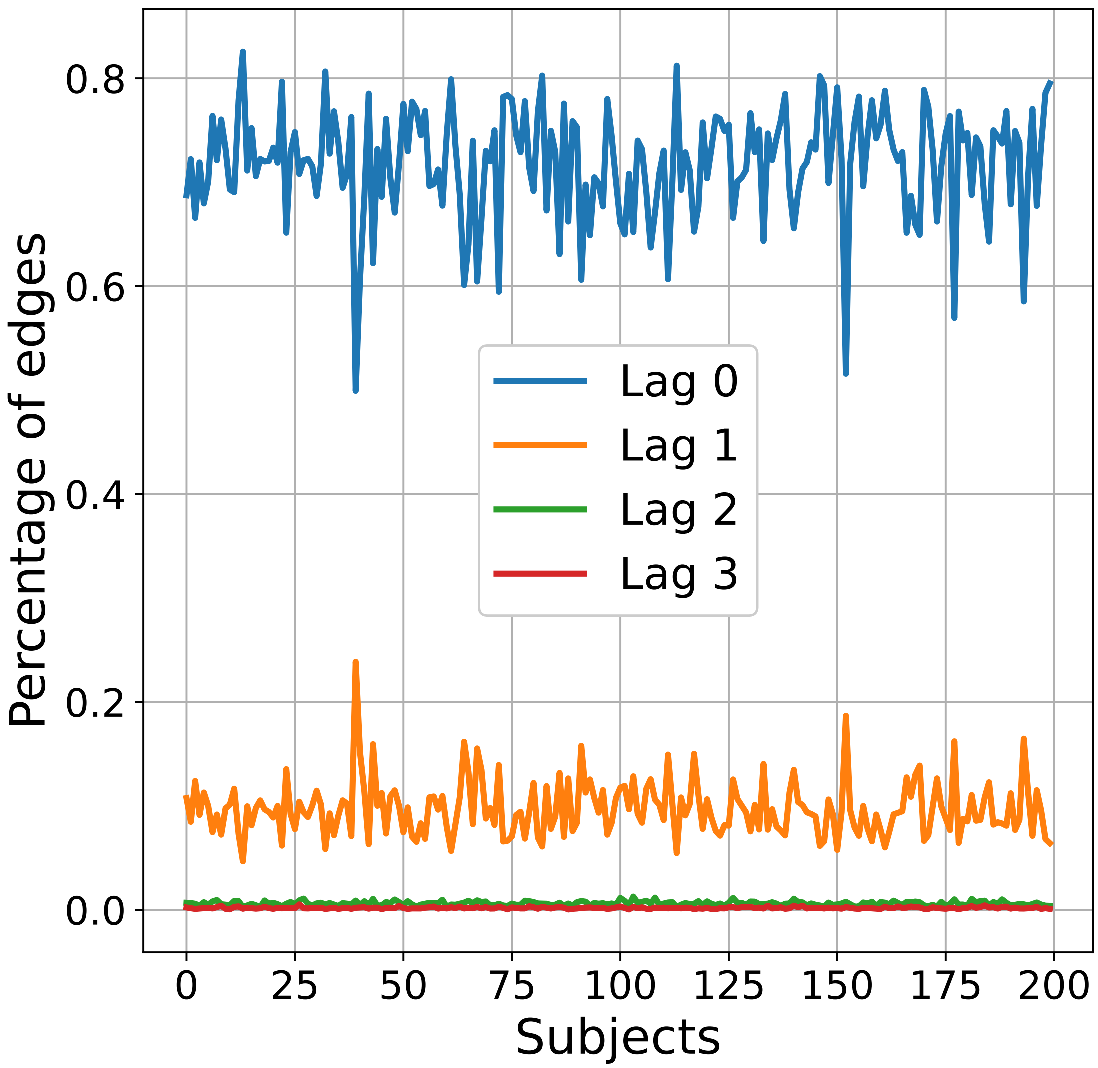

**Supplementary Figure 57: Percentage of edges present in each lag of the causal graphs learned by CaLLTiF over human fMRI data from HCP, shown separately for each subject.**

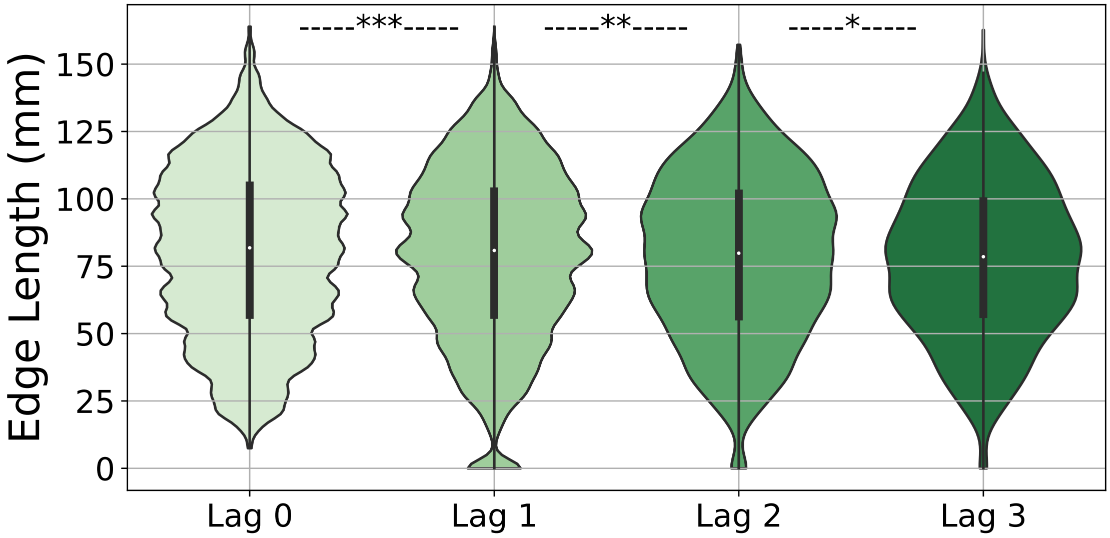

**Supplementary Figure 58: Comparison between Euclidean edge length of CaLLTiF edges at different lags.** Each violin plot shows the distribution of Euclidean edge length (parcel distance between edge endpoints) for edges existing in the subgraph corresponding to each lag. $^{*} = p < 0.05$, $^{**} = p < 0.01$, $^{***} = p < 0.001$, one-sided Wilcoxon rank-sum test.

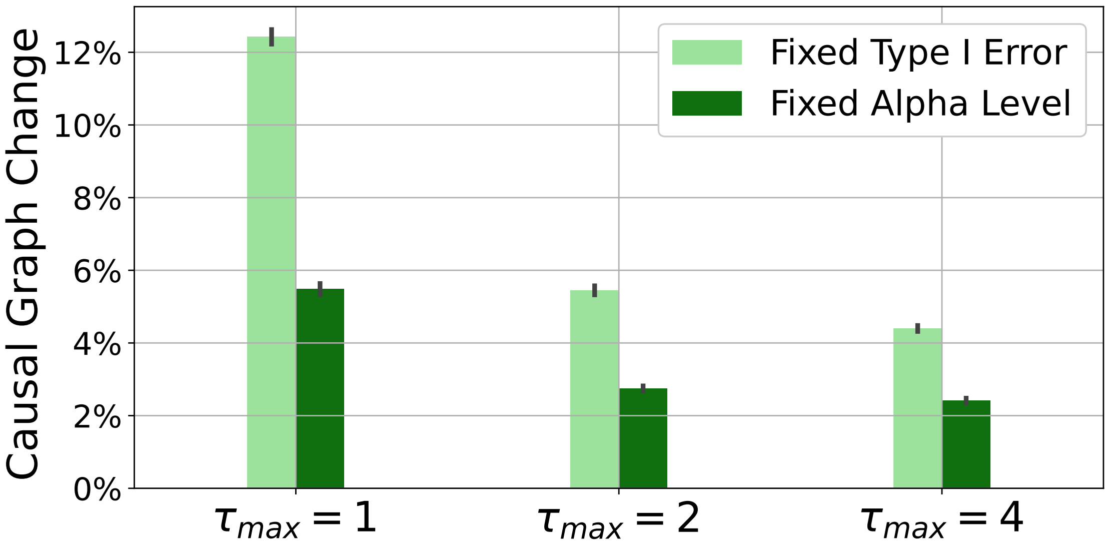

**Supplementary Figure 59: Sensitivity of CaLLTiF to $\tau_{\max}$ on HCP data.** Bar plots show mean and 1 s.e.m. of percent differences between the graphs computed using CaLLTiF with $\tau_{\max} = 1$, 2, and 4 compared to graphs computed with the nominal value of $\tau_{\max} = 3$. The difference is computed as the percentage of absolute change in the corresponding binary graphs computed for 15 subject. The 'Fixed Type I Error' and 'Fixed Alpha Level' cases correspond to when Eq. (4) in the main text is followed and ignored, respectively.

**Table 1: Names of regions in Schaefer 100x7 atlas (cortical parcels) and Melbourne Scale I atlas (subcortical parcels)**

| Parcel Number | Short Name |
|---|---|
| 1 | LH_Vis_1 |
| 2 | LH_Vis_2 |
| 3 | LH_Vis_3 |
| 4 | LH_Vis_4 |
| 5 | LH_Vis_5 |
| 6 | LH_Vis_6 |
| 7 | LH_Vis_7 |
| 8 | LH_Vis_8 |
| 9 | LH_Vis_9 |
| 10 | LH_SomMot_1 |
| 11 | LH_SomMot_2 |
| 12 | LH_SomMot_3 |
| 13 | LH_SomMot_4 |
| 14 | LH_SomMot_5 |
| 15 | LH_SomMot_6 |
| 16 | LH_DorsAttn_Post_1 |
| 17 | LH_DorsAttn_Post_2 |
| 18 | LH_DorsAttn_Post_3 |
| 19 | LH_DorsAttn_Post_4 |
| 20 | LH_DorsAttn_Post_5 |
| 21 | LH_DorsAttn_Post_6 |
| 22 | LH_DorsAttn_PrCv_1 |
| 23 | LH_DorsAttn_FEF_1 |
| 24 | LH_SalVentAttn_ParOper_1 |
| 25 | LH_SalVentAttn_FrOperIns_1 |
| 26 | LH_SalVentAttn_FrOperIns_2 |
| 27 | LH_SalVentAttn_PFCl_1 |
| 28 | LH_SalVentAttn_Med_1 |
| 29 | LH_SalVentAttn_Med_2 |
| 30 | LH_SalVentAttn_Med_3 |
| 31 | LH_Limbic_OFC_1 |
| 32 | LH_Limbic_TempPole_1 |
| 33 | LH_Limbic_TempPole_2 |
| 34 | LH_Cont_Par_1 |
| 35 | LH_Cont_PFCl_1 |
| 36 | LH_Cont_pCun_1 |
| 37 | LH_Cont_Cing_1 |
| 38 | LH_Default_Temp_1 |
| 39 | LH_Default_Temp_2 |
| 40 | LH_Default_Par_1 |
| 41 | LH_Default_Par_2 |
| 42 | LH_Default_PFC_1 |
| 43 | LH_Default_PFC_2 |

*Continued on next page*

Table 1 – *Continued from previous page*

| Parcel Number | Short Name |
|---|---|
| 44 | LH_Default_PFC_3 |
| 45 | LH_Default_PFC_4 |
| 46 | LH_Default_PFC_5 |
| 47 | LH_Default_PFC_6 |
| 48 | LH_Default_PFC_7 |
| 49 | LH_Default_pCunPCC_1 |
| 50 | LH_Default_pCunPCC_2 |
| 51 | RH_Vis_1 |
| 52 | RH_Vis_2 |
| 53 | RH_Vis_3 |
| 54 | RH_Vis_4 |
| 55 | RH_Vis_5 |
| 56 | RH_Vis_6 |
| 57 | RH_Vis_7 |
| 58 | RH_Vis_8 |
| 59 | RH_SomMot_1 |
| 60 | RH_SomMot_2 |
| 61 | RH_SomMot_3 |
| 62 | RH_SomMot_4 |
| 63 | RH_SomMot_5 |
| 64 | RH_SomMot_6 |
| 65 | RH_SomMot_7 |
| 66 | RH_SomMot_8 |
| 67 | RH_DorsAttn_Post_1 |
| 68 | RH_DorsAttn_Post_2 |
| 69 | RH_DorsAttn_Post_3 |
| 70 | RH_DorsAttn_Post_4 |
| 71 | RH_DorsAttn_Post_5 |
| 72 | RH_DorsAttn_PrCv_1 |
| 73 | RH_DorsAttn_FEF_1 |
| 74 | RH_SalVentAttn_TempOccPar_1 |
| 75 | RH_SalVentAttn_TempOccPar_2 |
| 76 | RH_SalVentAttn_FrOperIns_1 |
| 77 | RH_SalVentAttn_Med_1 |
| 78 | RH_SalVentAttn_Med_2 |
| 79 | RH_Limbic_OFC_1 |
| 80 | RH_Limbic_TempPole_1 |
| 81 | RH_Cont_Par_1 |
| 82 | RH_Cont_Par_2 |
| 83 | RH_Cont_PFCl_1 |
| 84 | RH_Cont_PFCl_2 |
| 85 | RH_Cont_PFCl_3 |
| 86 | RH_Cont_PFCl_4 |
| 87 | RH_Cont_Cing_1 |
| 88 | RH_Cont_PFCmp_1 |
| 89 | RH_Cont_pCun_1 |

*Continued on next page*

Table 1 – *Continued from previous page*

| **Parcel Number** | **Short Name** |
|---|---|
| 90 | RH_Default_Par_1 |
| 91 | RH_Default_Temp_1 |
| 92 | RH_Default_Temp_2 |
| 93 | RH_Default_Temp_3 |
| 94 | RH_Default_PFCv_1 |
| 95 | RH_Default_PFCv_2 |
| 96 | RH_Default_PFCdPFCm_1 |
| 97 | RH_Default_PFCdPFCm_2 |
| 98 | RH_Default_PFCdPFCm_3 |
| 99 | RH_Default_pCunPCC_1 |
| 100 | RH_Default_pCunPCC_2 |
| 101 | HIP-rh |
| 102 | AMY-rh |
| 103 | pTHA-rh |
| 104 | aTHA-rh |
| 105 | NAc-rh |
| 106 | GP-rh |
| 107 | PUT-rh |
| 108 | CAU-rh |
| 109 | HIP-lh |
| 110 | AMY-lh |
| 111 | pTHA-lh |
| 112 | aTHA-lh |
| 113 | NAc-lh |
| 114 | GP-lh |
| 115 | PUT-lh |
| 116 | CAU-lh |
